# The *Spirogyra* genome: signatures of shared and divergent division and differentiation

**DOI:** 10.1101/2025.10.09.681428

**Authors:** Elisa S. Goldbecker, Deepti Varshney, Anja Holzhausen, Tatyana Darienko, Hong Zhou, Armin Dadras, Lukas Pfeifer, Thomas Pröschold, Enrique Lopez-Gomez, Elke Woelken, Charlotte Permann, Tassilo Erik Wollenweber, Kerstin Becker, Stefanie König, Franz Hadacek, Fay-Wei Li, Ivo Feussner, Noe Fernandez-Pozo, Andreas Holzinger, Florian Maumus, André Marques, Henrik Buschmann, Klaus von Schwartzenberg, Jan de Vries, Stefan A. Rensing

## Abstract

Zygnematophytes emerged as the unexpected closest algal relatives of land plants despite their simple body plans, raising questions about the morphogenetic toolkit present in the last common ancestor of land plants and algae. Genomic analyses have revealed that zygnematophytes are cellular giants, sharing homologous frameworks for several phytohormones, secondary metabolites, and key morphogenetic and transcriptional regulatory processes. Zygnematophytes fall into five orders, each of which has charted its own evolutionary path. Here, we have sequenced a contiguous genome of *Spirogyra pratensis*, the eponymous representative of Spirogyrales and a classical model system for evolutionary cell biology in the green lineage. Building on this genome, we transcriptionally profiled the tractable life cycle of *Spirogyra* and its responses to a bifactorial gradient of light and temperature. Our data highlight the activation of quiescence and homeostatic programs. Yet what stands out most in *Spirogyra* is its spiral chloroplast—undulating intracellularly and abscising during mixed phragmoplast formation and furrowing. Leveraging the genome in tandem with co-expression network analyses, we describe the molecular underpinnings of the unique cytokinetic processes that govern both cell and plastid division. We find that *Spirogyra* deploys a molecular program characteristic of Phragmoplastophyta, yet lacks the deeply conserved plastid division machinery found in other archaeplastid plastids.

## Introduction

*Spirogyra* is a Zygnematophyceae, the streptophyte algal class sister to land plants^1–3^. Land plants (embryophytes), the organisms that constitute the entire macroflora on land, share a last common ancestor with Zygnematophyceae about 600 million years ago^4,5^. Zygnematophyceae are thus key for inferring the molecular toolkit of early land plants. Genomic exploration of streptophyte algae has revealed a suite of signature genes of land plants^6–12^, salient to phytohormone biology^13–18^, specialized metabolism^19–21^, plant–microbe interaction^22–24^, light signaling^25,26^, and more. Here, we show that *Spirogyra* has a divergent genetic complement, bringing about its unique biology.

The genus *Spirogyra* belongs to the Spirogyrales^27^, is comprised of ∼400 species^28,29^, and distributed worldwide with corresponding ecological amplitude. This rapidly proliferating alga is often found in blooms of nutrient-rich water bodies. This alga is characterized by uniseriate filaments with one to several spiral chloroplast(s) (Fig. 1a-c). The filaments occasionally branch to form lateral rhizoids. Sexual reproduction of *Spirogyra* is by conjugation of unflagellated cells.During conjugation in *Spirogyra*, two filaments align side by side, and the vegetative cells of both filaments form papillae toward each other, which subsequently fuse. The protoplasts of each cell move to each other to fuse and form zygotes (heavy-walled hypnospores called zygospores). The life cycle of the alga is haplontic and the meiosis occurs just prior to germination.

**Figure 1:**
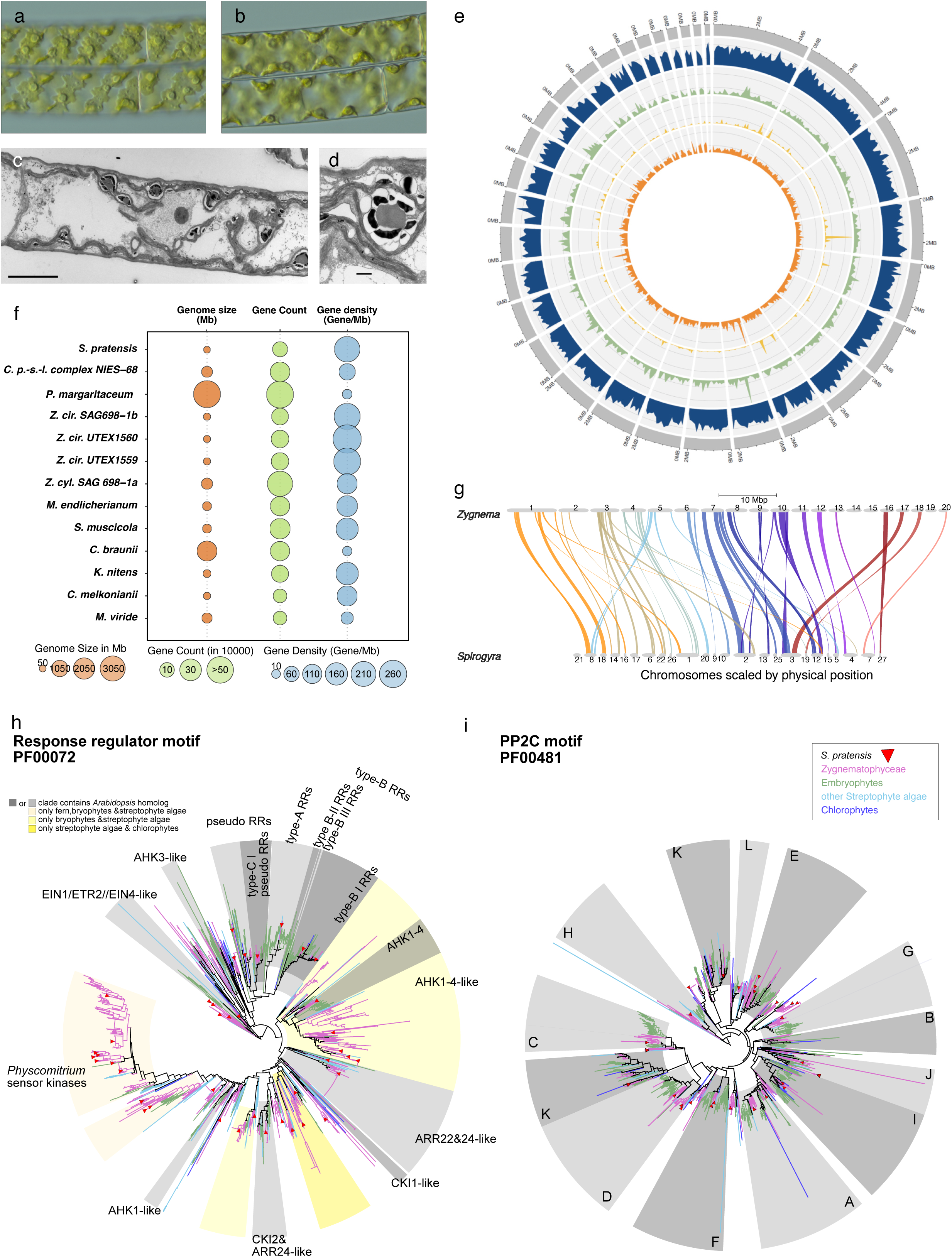
The *Spirogyra* genome. **a** and **b**: light microscopy images of the same filament of *Spirogyra pratensis* (MZCH10213) and detail of their spiral chloroplast; **c**: Transmission electron microscopy of *S. pratensis*, overview of vegetative cell, scale bar: 10 µm; **d** detail of chloroplast with pyrenoid, scale bar: 1µm; **e**: Distribution of the main classes of sequence types in *S. pratensis* genome with a 100 kbp sliding window. From outer to inner circle: Genes (blue); Transposable elements (green), the main satellite DNA repeat TR-128bp (light orange) and all repeats (dark orange). Note absence of clustering and relative uniform distribution of genes and repeats along the scaffolds. **f**: bubble plot showing annotation metrics of several streptophyte algae. **g**: synteny plot comparing *Zygnema* and *Spirogyra*; the chromosomes are gradient colored by the *Zygnema* chromosome numbers. **h** and **i**: response regulator (PF00072) and PP2C (PF00481) domain trees. Motifs were extracted from protein sequences using hmmer and aligned using mafft, phylogenetic trees were calculated using iqtree2 (SH-aRLT bootstrap 1,000); proteomes were downloaded from Genome Zoo (Table S15); clades were defined based on *Arabidopsis* genes; PP2C classes were annotated based on Xue et al.^40^.

*Spirogyra* is a classical model for cell biological research and has spawned important discoveries in the plant sciences. The mode of cell division in *Spirogyra* is enigmatic as it combines the ancestral cleavage mechanisms with the apomorphic cell plate-phragmoplast mechanism^30,31^. This discovery placed the class Zygnematophyceae safely in the group of Phragmoplastophyta^32,33^. Characteristic for *Spirogyra* cells are the spirally wound chloroplasts. Each cell has one or several left-handed ribbon-shaped chloroplasts with several pyrenoids (Fig. 1 a-d); the chloroplasts are located just beneath the plasma membrane. The plastids are not movable as in *Mougeotia* or *Mesotaenium*, but remain fixed in this parietal position, even though their shape and alignment is an expression of the physiological state of the alga. The spiral plastid is under tension, contracting when experimentally removed from the periplasm. First steps have been made to genetically modify *Spirogyra*.^34^

We here present the genome of *Spirogyra pratensis* MZCH10213. We find that this alga has an unusually small genome, packing 11,797 protein-coding genes on 55 Mbp. Comparative genomic analyses reveal the loss of plastid division genes, while we recover co-expression of cell division and environmental response hubs.

## Results

### *Spirogyra* has a compact and gene dense genome

The *S. pratensis* genome was sequenced to 74x coverage and assembled into 31 scaffolds, which sum up to 55.44 Mbp, the longest scaffold being 4.49 Mbp (Fig. 1e). The N50 and N90 values are 2,195,869 bp and 1,234,611 bp respectively. The predicted assembly size was found to be 45 Mbp by k-mer analysis with the best k-mer value = 21 (Fig. S1). Structural gene annotation predicted 12,264 genes, with 11,797 protein-coding. The *Spirogyra* genome has a high gene density of 221.2 genes/Mbp, resembling another filamentous Zygnematophyceae species with a small genome, *Zygnema circumcarinatum*^10^ (Fig. 1f; Table S6). Despite the relatively low gene number, Benchmarking Universal Single-Copy Orthologs (BUSCO; viridplantae_odb10) showed high completeness for the *Spirogyra* annotation (C:94.6% [S:88.2%, D:6.4%], F:1.2%, M:4.2%, n:425) (Fig. S2); reads from RNA-seq mapped at an average rate of 85.6% (Table S2).

MEGAN analysis of the scaffolds found the highest percentage of hits among Viridiplantae, as expected (Table S5), and no evidence for contamination (see Methods). Hits against other taxonomic groups occurred exclusively in non-genic repetitive regions of the genome. The assembly (hap1) contains a significant amount of simple sequence repeats (SSRs) i.e. tandem repeats^35^ and low-complexity sequences. A soft SSR masking yields 19% coverage and an aggressive run masks over 47% of the assembly.

As expected for a small genome, dispersed repeats contribute ∼13.1% of the assembly (hap1). While most repeats remain unclassified (Table S4), the few bona fide transposable elements (TEs) detected include Copia- and Gypsy-type LTR retrotransposons (class 1 TEs), and hAT and putative Plavaka DNA transposons (class 2 TEs; Fig. 1e). Plavaka elements have so far been reported only in fungal genomes^35^, yet we detect them in various Archaeplastida—red algae, chlorophytes, and streptophytes (algae and moss). Multiple sequence alignment of their transposase domain indicates highly conserved residues shared with the CMC (CACTA, Mirage, Chapaev) clade^36^ of DNA transposons. Phylogenetic analysis placed plant Plavaka candidate sequences with fungal Plavaka elements in a distinct cluster within the CMC clade, supporting Plavaka as a separate superfamily (Fig. S3A). The data suggest Plavaka was present in the Archaeplastida last common ancestor (LCA) and was later lost in land plants (Fig. S3B). Among *S. pratensis* Gypsy-type TEs, one chromodomain-containing family (R226) shares ∼50% RT domain identity with oomycete homologs, suggesting a recent horizontal gene transfer (HGT) from an oomycete (Fig. S4). A second family (G72) shows >50% identity to fungal sequences, especially mucoromycotan fungi, while more divergent homologs are found in other eukaryotes. (Fig. S4). Taken together, several components of the TE repertoire in *S. pratensis* appear to originate from HGT events.

We identified a main family of a satellite DNA repeats with a monomer length of 128 bp showing a dispersed distribution across all scaffolds (Fig. 1e). This satellite repeat showed an array size average length of 2 kbp and inter-array spacing of 256 kbp. The pattern of satellite DNA distribution resembles plant species with repeat-based holocentromeres^37,38^, aligning with the longstanding notion^39^ that *Spirogyra* has holocentric chromosomes. Yet, further validation of centromere composition in is required in order to determine the centromere type.

### Transcription-associated proteins: evidence for lineage-specific gene loss, and for expansion of two component signaling

Transcription associated proteins (TAPs) include transcription factors (TFs) and transcriptional regulators (TR). All *S. pratensis* proteins were classified into 138 different TAP families using TAPScan^40^ v4. *S. pratensis* has 544 TAPs (TF = 289; TR = 202; putative TAPs = 53) accounting for 4.6% of the protein-coding genes (Table S7, Fig. S5). The number of TAPs in *S. pratensis* is lower than in land plants, resembling Zygnematophyceae species such as *Zygnema circumcarinatum* strains UTEX 1559 (n = 576), UTEX 1560 (n = 560) and SAG 698-1b (n = 528). Other Zygnematophyceae have larger complements—*Penium margaritaceum* (n = 1,323), and *Spirogloea muscicola* (n = 1,183)— likely due to duplication events^8,9^. The higher number of TAPs reported before for the strain *Spirogyra pratensis* UTEX 928 (n = 2,622), from which the present strain S. pratensis MZCH10213 derives, is likely to be due to the draft-stage of the *S. pratensis* UTEX 928 genome^22^ riddled with (near-)identical proteins (Figs. S6-8).

The two *S. pratensis* genomes (i.e. MZCH10213 and UTEX 928), despite their different absolute numbers of TAPs, have a similar pattern of absence of gene families: the TF families CRF, CudA, DBP, GeBP, HD_DDT, HD_PINTOX, HD_TALE_BEL, HD_TALE_KNOX2, LFY, MYB-4R, NZZ, RF-X, Rel, Runt, SAP, TEA, ULT, VARL, VOZ, bHSH, and the TR families Aux/IAA, Dicer, GIF, Jumonji_PKDM7, OFP, PcG_EZ, ZPR are absent from both, although present in all or several of the other Zygnematophyceae genomes (Table S7). *Spirogyra* likely experienced lineage-specific loss of TAP genes. To scrutinize this, the 1KP^117^ species tree was used as basis of CAFE5 analysis^41^. *S. pratensis* showed expansion in 4 and contractions in 46 of TAP families, the latter including AP2, bHLH, bZIP, MYB-related.

We noted that the fraction of genes classified as “no family found” is comparatively high in *S. pratensis* (n = 277; 2.3%; Table S8). In other Zygnematophyceae the fraction ranges from 0.7% to 1,6%, with the exception of *S. muscicola,* 2.6%). This classification occurs if genes contain one or more domains used for TAP classification, but in a combination that does not allow to assign a family. For example, 71 with only DEAD/helicase_c (not full Dicer), 20 with PP2C (but not DNC for TF DBP), and 35 with response_reg (unassigned to ARR-B/GARP families). Of 130 WD40-containing proteins, only 9 map to known families like FIE or MSI. Such cases likely occur due to domain swapping or transposon events.

To trace domains, we computed trees for the most abundant domains in this category (DEAD, PP2C, Helicase_C, Response_reg, WD40). For DEAD box (PF00270), Helicase C (PF00271) (Fig. S9) and PP2C (PF00481; Fig. 1i) most clades harboring Arabidopsis proteins also contain one or a few Spirogyra proteins, there are no clades that harbor a high number of Spirogyra proteins – hence, there is no evidence for lineage-specific expansion. For WD40 (PF00400; Fig. S8) there are some clades in which there are many *S. pratensis* and in general Zygnematophyceae sequences, such as the one harboring AT4G38480 (ABA-HYPERSENSITIVE DCAF 1; ABD1) in which there are three *A. thaliana* and 12 *S. pratensis* proteins. Phylogenetic analysis of Response_reg domains (PF00072) reveals clades lacking Arabidopsis, containing i) fern, bryophyte, and streptophyte algae, ii) bryophyte and streptophyte algae, or iii) streptophyte and chlorophyte algae sequences (Fig. 1h). This includes a clade containing three *P. patens* and eight *S. pratensis* proteins (Fig. S6). Two of the three *P. patens* proteins are annotated as histidine sensor kinases (Pp3c16_14810V3.1 and Pp3c27_2730V3.1; v3.3 models). Analysis of co-expressed genes during *P. patens* development in MAdLandExpression (https://peatmoss.plantcode.cup.uni-freiburg.de/easy_gdb/index.php) showed five co-expressed genes with a correlation score of 90% or more for Pp3c16_14810V3.1, related to signal transduction, cysteine biosynthesis, and sulfate transport. A total of 38 genes are co-expressed with Pp3c27_2730V3.1, showing functions related to transmembrane transport, transcription regulation, signal transduction, cysteine biosynthesis, photosynthesis, photosynthetic electron transport chain, biosynthesis and development. While type-B response regulator motifs are present in chlorophytes, streptophyte algae (including *S. pratensis*) and embryophytes, type-A RRs are only present in embryophytes, in *Klebsormidium nitens* as well as in both sampled *Spirogyra pratensis* strains (Fig. S7). PROTEIN PHOSPHATASE 2C (PP2C) motifs for all except class B are present in *S. pratensis* (Fig. 1 i). The lineage specific expansion of response regulator containing genes hints at an elaborate two component system that, together with the PP2C containing proteins, might convey stress signals in *S. pratensis*, while the expanded WD40 proteins might convey lineage-specific protein-protein interaction functions.

### Abundant tandem and proximal duplicates, but no relationship with expanded domains

To assess gene duplication events in *S. pratensis,* we used the doubletrouble tool^42^ to classify genes into four duplication categories: segmental duplication (SD), tandem duplication (TD), proximal duplication (PD), and dispersed duplication (DD) (Table S10, Fig. S10). While there are many SD detectable in the two genomes known to have been subject to whole genome duplications (*P. patens* and *S. muscicola*), this type of duplication is absent or of very low abundance in the other species. *S. pratensis* shows a relatively high number of TD and PD as compared to most other Zygnematophyceae, and a relatively low number of DD. Among the above-mentioned 130 WD40 domain containing genes (Table S16), 15 were classified as TD (n = 4) or PD (n = 11), for the 40 response_reg domain (Table S17), 6 genes as TD (n = 4) or PD (n = 2). Iterated statistical testing against random control samples shows that these genes are not enriched for proximal (TD and PD) duplication events (p > 0.5, Chi squared test). Hence, there is no apparent relationship between proximal duplication events and expansion of these genes (Table S16 & S17).

### Cell wall-related enzyme toolkit: species-dependent varying redundancy in Zygnematophyceae makes *Spirogyra* an optimal organism for studying cell wall evolution

We annotated all CAZymes (glycosyltransferases, GTs; glycosylhydrolases, GHs; polysaccharide lyases, PLs; carbohydrate-binding modules, CBMs; carbohydrate esterases, CEs; enzymes with auxiliary activities, AAs; Table S26-28), focused on cell wall-related enzymes and inferred homology to functionally described plant enzymes (Figs. S11-15). As in other Zygnematophyceae^10^ homologs for biosynthesis and remodeling of most cell wall polysaccharide structures, except for rhamnogalacturonan-II, are present. There is often a species-dependent higher or lower multiplication in homolog numbers which does not follow any obvious trend. For some polysaccharide groups, namely heteromannan, and homogalacturonan, *Spirogyra* contains only the basic enzymatic toolkit (one homolog per defined backbone-/sidechain-linkage type) for biosynthesis. Xyloglucan biosynthesis genes are far less numerous than in embryophytes. The glycosyltransferases IRX9 and IRX14, which form a complex with IRX10 for xylan backbone synthesis, are present as single copies. In contrast, GUX1-5 and XAT homolog numbers are like those in embryophytes, unlike most streptophyte algae. ESK1 homologs, which acetylate xylan, are found in all Zygnematophyceae but not in other streptophyte algae. Genes involved in AGP galactan core biosynthesis (GALTs, HPGTs, GALT31A) show low redundancy, and no close homologs of methylation-related enzymes (IRX15/15L, GXMT1-3, AGM1-2) are found in *Spirogyra* or other streptophyte algae. As there are descriptions of *O*-methyl sugars in *Spirogyra* and other streptophyte algae^43,44^, these hint towards presence of more but currently unknown enzymes catalyzing methyl-decoration of cell wall polysaccharides.

### The tractable life cycle of *Spirogyra*

In liquid airlift culture, *S. pratensis* remained mainly in the vegetative stage. Transfer to agarised medium induced scalariform or lateral conjugation at high frequencies after about 6 d (see time lapse in Supplemental Video S1). Zygospores were formed in the recipient cell and turned brownish after ∼14 d (Fig. 2a, 2b, S16). Following several weeks of drying at ambient air, the remaining vegetative cells died and dark mature zygospores could be harvested mostly still located in the recipient gametangial cell. Cold-treated (4°C, dark) mature zygospores occasionally germinated after ∼28 days at 15°C or 25°C on fresh medium. They swelled, turned green, and formed filaments with spiral chloroplasts that emerged from the gametangial cell. Three germinating zygospores were subcultured and deposited as MZCH strains #762, #763 and #764.

**Figure 2:**
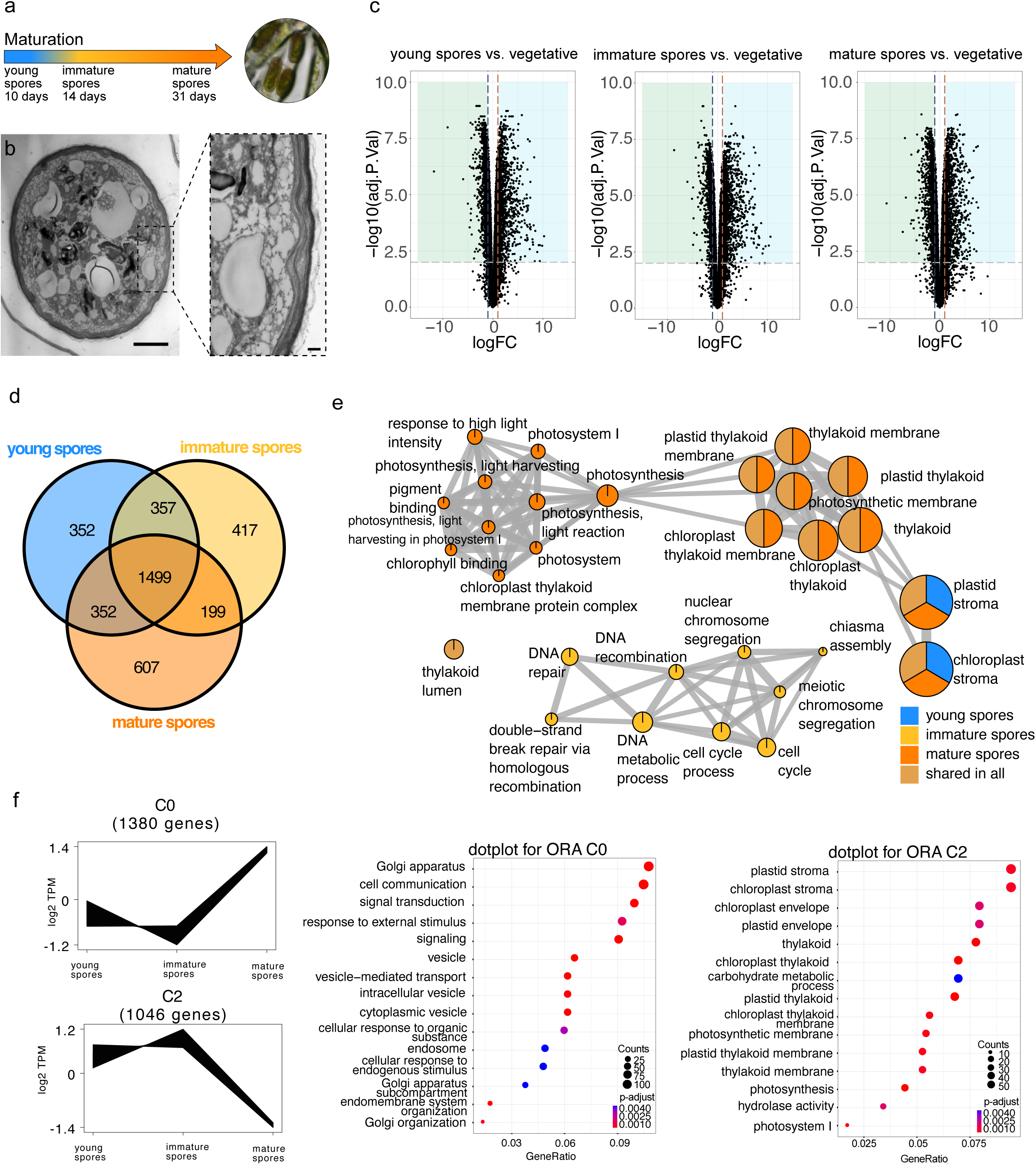
Transcriptional tracing of the life cycle of *S. pratensis*. **a**, Schematic of experimental design showing sampled zygospore stages b, Transmission electron microscopy images of young *Spirogyra* zygospore and detail of three-layered zygospore cell wall, scale bars: 5 µm, zoom-in: 500 nm. **c**, Volcano plots of differentially expressed genes between zygospore stages (young, immature, mature) and vegetative *Spirogyra* culture. Genes were considered differentially expressed at |log2Fold change| ≥1 and Benjamini-Hochberg corrected *P*≤0.01. **d**, Venn diagram of differentially expressed genes between the three zygospore stages. **e**, Biological theme comparison of Venn diagram intersections. “Shared in all”: in all three spore stages. “Young-, immature- and mature spores” contain differentially expressed genes exclusive to that life stage. **f**, Expression profiles and GO term enrichment of two clusters recovered by the *clust* algorithm.

We performed RNA-seq on samples enriched for different life cycle stages and compared gene expression across three zygospore stages (young, immature, mature) and vegetative filaments (Fig. 2a), revealing marked transcriptional differences between zygospore and vegetative stages in *Spirogyra* (Fig. 2c). The bulk of gene expression changes were shared between all spore stages (1,499 genes). This is to be expected since the enriched sets also contain vegetative cells or their remains. Yet, there were also large stage-specific gene expression changes (Fig. 2d). GO terms enriched in the shared zygospore specific differentially expressed genes (DEGs) were related to photosynthesis (Fig. 2e), likely reflecting a disassembly of the photosynthetic apparatus during zygospore development^45^. The signal for photosynthesis-related terms was even more pronounced in young and mature spore specific DEGs. Among DEGs specific for immature spores we found enriched GO terms like “meiotic chromosome segregation”, “DNA recombination”, and “cell cycle” pointing towards the timing of meiosis in the immature spore sample. Gene expression dynamic analysis across the three zygospore stages using clust^46^ recovered three clusters. Cluster C0 captures upregulation of genes during maturation and was enriched in GO terms related to signaling and Golgi mediated transport (Fig. 2f), suggesting dynamic intracellular reorganization; cluster C2 showed the downregulation of photosynthesis related genes during maturation, reflecting the controlled onset of quiescence.

### The molecular physiology of *Spirogyra* is dominated by light response

During plant terrestrialization, the earliest land plants were challenged by a unique set of stressors^47^. Enduring of stress by dormant zygospores is one avenue but also the vegetative cells need to adjust their physiology. To understand the molecular physiology of vegetative *Spirogyra*, we performed a bifactorial gradient experiment as previously described^26^. Briefly, *Spirogyra* was cultivated to apt density and spread across 42 twelve-well plates (2.5 mL of culture per well). The plates were placed on a bifactorial light and temperature gradient setup with light intensities ranging from 21 to 530 µmol photons s^−1^ m^−2^ (spectrum as in Dadras et al.^26^, Fig. S18) and temperatures from 8 °C to 29 °C; the experiment was repeated in three successive runs, yielding three independent biological replicates for each of the 42 conditions. We used pulse amplitude modulation fluorometry (PAM) to assess the maximum quantum yield of photosystem II (Fv/Fm) as a proxy for the gross physiology (Fig. 3a). No significant differences of the Fv/Fm to control conditions was observed under medium to high temperatures (17°C - 29°C) and low light conditions (21 µmol photons s^−1^ m^−2^), while values close to 0 were obtained in high light-stressed samples (530 µmol photons s^−1^ m^−2^; Fig. 3b). Overall, the increase in light intensity led to a stronger decrease in Fv/Fm than any changes in temperature, aligning with the phenotype: while *Spirogyra* cultures were still green under high temperature and low light (29°C and 21 µmol photons s^−1^ m^−2^) and low temperature and low light (8°C and 21 µmol photons s^−1^ m^−2^), they appeared bleached under high temperature and high light (29°C and 530 µmol photons s^−1^ m^−2^) and low temperature and high light (8°C and 530 µmol photons s^−1^ m^−2^; Fig. 3c). This might be a function of the chloroplast, unlike in other Zygnematophyceae, not being subject to light-induced movement.

**Figure 3:**
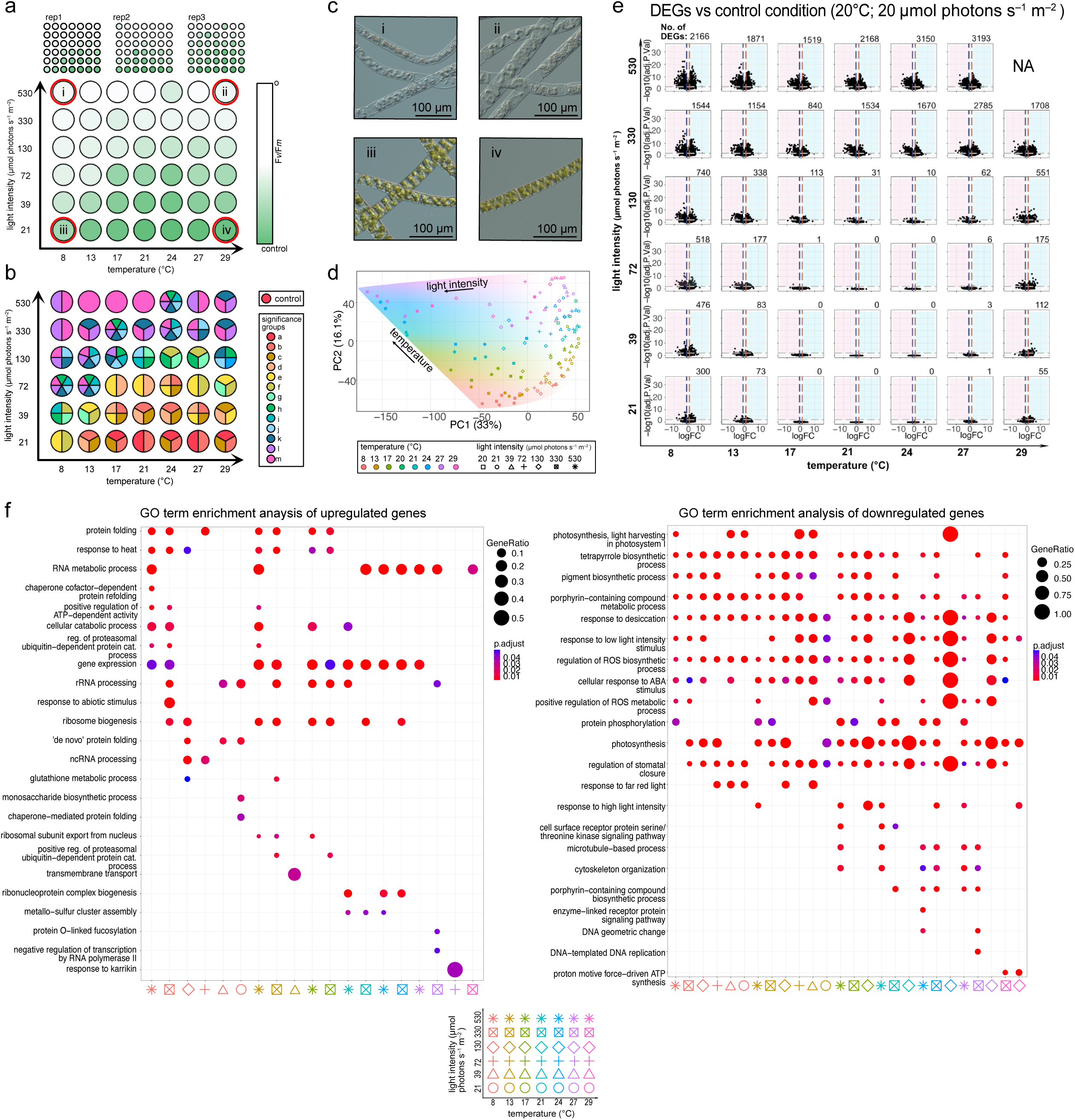
Response of *Spirogyra* to a bifactorial environmental gradient. **a,** The gross physiology of *Spirogyra pratensis* was assessed by measuring the maximum yield of photosystem II (Fv/Fm) after 65 h of treatment and in all three replicates of the gradient experiment. **b**, Significance groups were determined using Kruskal-Wallis test with Fisher’s least significant difference post-hoc test. Groups were considered significantly different at Bonferroni-adjusted p ≤ 0.001. **c**, Microscopy pictures show representative images of the *Spirogyra* cultures taken at 89 h from the most extreme conditions of the experimental setup. **d**, PCA plot of RNA-seq samples. **e**, Vulcano plots of differentially expressed genes. DEGs were determined by comparing samples from the gradient table against control samples grown at (20 °C and 20 µmol photons s^−1^ m^−2^). Genes were considered differentially expressed at |log2Fold change| ≥1 and Benjamini-Hochberg corrected p≤0.01. Numbers in the right corner of the volcano plots show the total amount of differentially expressed genes per condition. **f**, GO term enrichment of up-(left) and down-regulated genes (right). Enrichment shows redundancy removed biological process (BP) GO terms.

### Differential gene expression patterns reflect photophysiology and high light stress

To explore the molecular basis of the physiological differences, we performed global RNA-seq analysis across all conditions and biological triplicates, including controls, yielding a total of 129 distinct samples, of which 126 were usable (41 conditions plus controls) (Fig. S19). We obtained a total of ∼1.26 Tbp of 150 bp paired-end reads (on average 32.4 million reads per sample; ∼8.4 billion reads in total). These were mapped to the *Spirogyra* transcripts at a rate of 84%. Principal component analysis (PCA) of the RNA-seq data showed a clear separation of the samples along temperature and light intensity axes that closely matched PC1 and PC2 (Fig. 3d). The strong effect of light on the main PCs was confirmed using small multiple PCA analysis (Figs. S20-21). The strongest gene expression changes occurred under high light; further cold temperatures had a greater effect on DEGs than warm ones. This aligns with our photophysiological and phenotypic data identifying high light and cold as the most impactful stressors. While high light combined with high temperature likely caused the greatest disturbance, this extreme could not be fully assessed by RNA-seq (Fig. 3e, “NA”).

Upregulated genes across all conditions were enriched in GO terms related to RNA metabolism like “gene expression” or “RNA metabolic process” (Fig. 3f), mirroring activation of genes. Under 8°C to 17°C and high light conditions, we found GO term enrichment for “protein folding” and temperature responses, potentially reflecting protein folding stress. The cold and low light intensity condition 8°C and 21 µmol photons s^−1^ m^−2^ was the only low light condition that showed significant GO term enrichment among upregulated genes, which were “monosaccharide biosynthetic process” and “chaperone-mediated protein folding”. Heat specific GO terms in upregulated genes (27-29°C) included “protein O-linked fucosylation” and “response to karrikin”. Both GO terms reflect genes encoding enzymes for substrate methylation (nucleic acids) or fucosylation (proteins). Even though there are homologs of the plant specific protein O-fucosyltransferase SPINDLY (SPY) and its close paralog SEC in Spirogyra (Table S22, Fig. S22), the upregulated genes are more similar to the protein O-fucosyltransferases in glycosyltransferase family 68, playing important roles in posttranslational modification of NOTCH signal proteins in animals (Suppl. Table S20 &S22). GC-MS analysis of central metabolites of the gradient table samples revealed heat stress specific increases in amino acid (tryptophan, valine, isoleucine, leucin, alanine, proline and threonine), di- and trisaccharide sugars (sucrose, trehalose, raffinose) and citrate abundances (Extended Data Fig. 2), aligning with previous reports on same strain of *Spirogyra pratensis*^48^ and for *Chlamydomonas reinhardtii*^49^. The increases in specific amino acid abundances under high temperatures are likely not only due to protein degradation or decreased synthesis as they do not match the amino acid frequencies of the predicted *Spirogyra* proteome (Table S29).

By comparing the gene expression of our samples to control samples that were grown at 20°C and 20 µmol photons s^−1^ m^−2^, we recovered mainly stress specific responses in the DEGs. Overall, GO terms enriched among downregulated genes were associated with photosynthesis, capturing the light stress response that is overrepresented in the DEGs, aligning with previous findings^26,50^. Next to these general cell physiological readouts, we found links to key stress signaling cascades including “response to desiccation” and GO terms associated with ROS metabolism and “cellular response to ABA stimulus“, representing functions that are depleted in our experimental set-up. Under mid to high temperatures (17 – 27°C) and high light intensities (330 - 530 µmol photons s^−1^ m^−2^) genes related to “cytoskeleton organization” were downregulated. Genes downregulated under heat conditions (24 – 29°C) had GO terms enriched in ATP synthesis and DNA replication, reflected a stalled metabolism under heat.

Data can be explored at the MAdLandExpression atlas (https://peatmoss.plantcode.cup.uni-freiburg.de/easy_gdb/index.php) (Fig. S23).

### Co-expression analysis reveals molecular programs of *Spirogyra pratensis*

To examine the biological programs in *Spirogyra pratensis* we performed signed Weighted Gene Co-expression Network Analysis (WGCNA). We clustered the gradient RNA-seq data from 11,169 well-expressed genes into 29 co-expression modules (Fig. S25, Table S30)) and calculated their module-trait correlations to the experimental parameters light intensity, temperature and photophysiology (Fv/Fm) (Fig. 4a, Fig. S29). For example, modules red (Fig. 4c) and lightgreen correlated with high Fv/Fm values—indicative of unstressed and well-growing algae—and were enriched in GO terms for cell wall biogenesis (lightgreen), chromosome and cellular organization (red; Fig. 4b).

**Figure 4:**
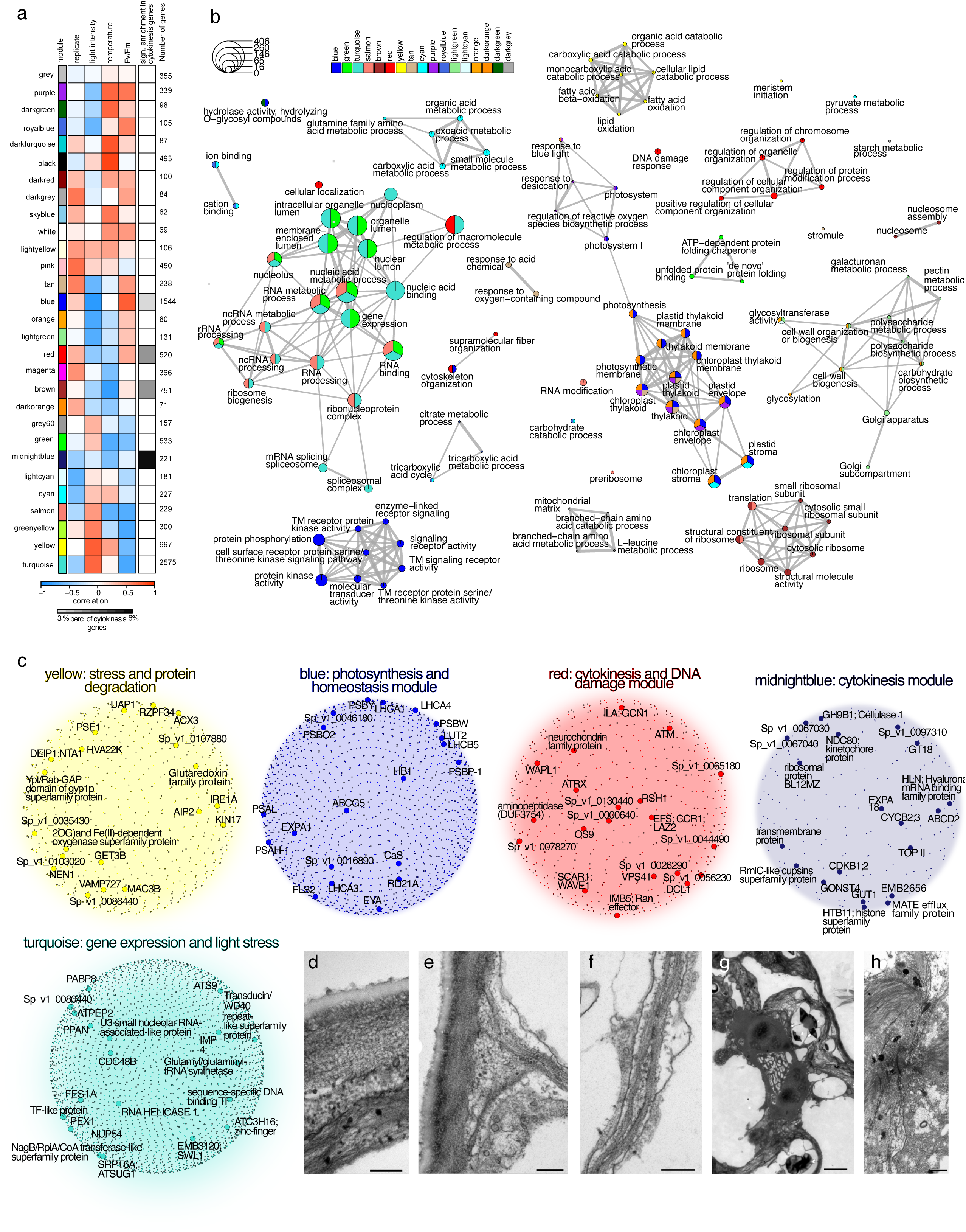
Weighted Gene Co-Expression Network Analysis of environmental gradient RNA-seq data. **a**, module-trait correlation of recovered WGCNA modules. The experimental parameters temperature, light-intensity and Fv/Fm were correlated to the eigengenes of co-expression modules. Correlations range from blue (strong negative correlation) to red (strong positive correlation). Enrichment for cytokinesis related genes was determined by testing the distribution of a cytokinesis related gene set within the modules using Pearson’s chi-squared test(χ2), p <0.05, df=1. **b**, GO-term based comparison of biological themes of the co-expression modules. **c**, Representative co-expression modules with highlighted hub genes. Hub genes were defined as the 20 genes with the highest connectivity within that module, and are highlighted. Annotations were transferred from a BLASTP-analysis against *Arabidopsis thaliana*. **d-h** Transmission electron microscopy of *Spirogyra pratensis* (MZCH10213) **d**. multilayer cell wall, scale bar: 250 nm; **e** connection of cross wall to external cell wall, scale bar: 500 nm; **f** Cross wall with intracellular space, scale bar: 500 nm; **g** nuclei after division in vegetative cell, scale bar: 2 µm; **h** young cross wall of vegetative cell after centripetal cell division, scale bar: 500nm.

Modules yellow and turquoise have a strong positive correlation with light intensity and a strong negative correlation with Fv/Fm values, indicative of genes important for high light stress responses; GO terms enriched for turquoise relate to RNA metabolism (Fig. 4b). The top 20 most connected genes (hubs) in module turquoise speak of intracellular responses, featuring: PETER PAN-LIKE PROTEIN (PPAN) and POLY-A BINDING PROTEIN 8 (PAB8) related to RNA metabolism^51,52^ reflecting the transcriptional response to the stress (Fig. 4c); peroxisome and chloroplast biogenesis and remodeling hubs peroxisome 1 (PEX1) or SNOW-WHITE LEAF 1 (SWL1)^53,54^(Table S69). Additionally chloroplast hubs, including CLP proteases (CLPPs) key for high-light acclimation and chloroplast protein quality control^55,56^, and the chloroplast control hub GENOMES UNCOUPLED 2 (GUN2)^57–59^ are part of the turquoise module.

Module yellow is enriched in genes for fatty acid and lipid metabolism (Fig. 4b). Hubs in yellow relate to the proteasome and ubiquitination: (i) the ubiquitin associated protein 1 (UAP1/FLOE2) that is also involved in dehydration response^60^ and is homologous to FLOE1, forming condensates potentially mediating the water status of germinating *Arabidopsis* seeds^61^; (ii) the universal ER stress sensor kinase INCREASED ORGAN REGENERATION 1 (IRE1)^62^ and the SNARE complex protein VESICLE ASSOCIATED MEMBRANE PROTEIN 727 (VAMP727)^63^ suggesting protein degradation under light stress (Table S70).

In contrast, the blue module has the strongest positive correlation with Fv/Fm and a strong negative correlation to light intensity, which indicates that this module is active under conditions in which the algae are in homeostasis - the GO terms enriched for blue mainly relate to signaling (Fig. 4b). The blue module has several hub genes related to photosynthesis, mirroring its strong positive correlation to Fv/Fm values and thus active and effective photosynthesis. The most strongly connected hub gene was identified as FLAGELLIN-SENSITIVE 2 (FLS2) by best BLASTP hits against *Arabidopsis* but did not fall into one orthogroup with Arabidopsis FLS2 (Table S71). Instead, it formed an orthogroup with only Zygnematophyceae species and might resemble an LRR-kinase hub conserved in Zygnematophyceaen algae^50^.

We compared the responses of *Spirogyra* with another member of the Zygnematophyceae algae, the single celled *Mesotaenium endlicherianum*, that was subjected to the same bifactorial gradient setup^26^ (Extended Data Fig. 1a, b). Most shared differentially regulated OGs occurred under high light and high temperatures, as these also yielded the most DEGs in both experiments (Extended Data Fig. 1c); the shared OGs had enriched GO terms associated with signaling (Extended Data Fig. 1d). The blue modules of both species showed high Jaccard similarity, with shared hub genes including orthologues of the calcium sensor CAS^64^ and the LRR-kinase hub described above (Extended Data Fig. 1e), stress signaling programs appear mediated by the same orthologs and upon the same environmental cues in the more than 500 million-years-divergent species.

### Divergence in *Spirogyra* cell division

For comparative analyses of cell division in *Spirogyra* (Fig. 4d-h), we manually curated a list of 288 genes for cytoskeleton and endomembrane transport, and well-characterized upstream regulatory cell division regulators. Most cell division genes are present in all streptophytes, showing their deeply conserved role in streptophyte cell biology^4,11,65^ (Table S9). Strikingly, a few cell division genes are absent from *Spirogyra* (or highly divergent), including the centromere/kinetochore-localized mitotic spindle check-point factors INCENP and MAD2; their absence is consistent with the lack of a defined centromere in *Spirogyra*^66^.

*Spirogyra* belongs to the Phragmoplastophyta and like the zygnematophyte *Mougeotia*, exhibits a hybrid form of cytokinesis combining phragmoplasts and cleavage furrows^31,67^. To trace cytokinetic regulators, we used a χ2 test assessing enrichment of orthogroups that contain the 288 cytokinesis related genes in the co-expression modules (Fig. 4a, Table S24)). Four modules are significantly enriched (p < 0.05) in cytokinesis genes, with midnightblue (6%), red (4%) and brown (4%) showing the highest enrichment. Midnightblue consists of 221 genes of which 13 were marked as cytokinesis related, of which homologs of cyclin B (CYCB2;3), the cyclin-dependent kinase B (CDKB2;3) and the glycosyl hydrolase 9B1 (GH9B1) were hub genes (Table S73). Additionally, it contained cytokinesis related hub genes that were not listed in our cytokinesis gene list like a member of the alpha-expansin family (EXPA18) that are involved in cell wall loosening and cell expansion^68,69^ or TOPOISOMERASE II (TOP II) involved in DNA replication^70,71^. Parts of the cytokinesis machinery were also found in other cytokinesis enriched modules, for example two orthologs of the PHRAGMOPLAST ORIENTING KINESINs POK1 and 2, necessary for the orientation of the cell division plane^72^, were found in the blue module. One ortholog was co-expressed with genes involved in cell wall biogenesis (GSLs, GLUCAN SYNTHASE-LIKE)^73^ and alignment of the sister chromatids (STRUCTURAL MAINTENANCE OF CHROMOSOMES 3, SMC3)^74^ while the other ortholog is co-expressed with the kinesin-like calmodulin-binding protein ZWICHEL (ZWI) and STRUCTURAL MAINTENANCE OF CHROMOSOMES 2B (SMC2B) (Suppl. Tables S2-3). Overall this highlights that *Spirogyra* ZWI and POKs are involved in cell division and aligns with previous finding of POK as a hub in co-expressed cell division genes of *Mesotaenium*^26^ — which we now can pinpoint as shared between filamentous and unicellular zygnematophytes.

Fission by cleavage furrows is best studied in unikonts where it is mediated by filamentous actin and type II myosin^75^. However, *Chlamydomonas*, which divides via cleavage furrow, lacks the unikont-specific myosin-II and while actin localizes to the division plane it is not required for fission^76^. *Spirogyra* has two actin orthologues that clustered into to the blue and magenta module. The actin ortholog of the blue module showed strong co-expression with GLUCAN SYNTHASE-LIKE 8 (GSL8) and AUXIN-INDUCED IN ROOT CULTURES (AIR9) that is important for division plane establishment^77^ hinting that it is likely involved in cytokinesis in *Spirogyra* (Suppl. Table S1). Interestingly, Zygnematophyceae have massive expansions in a plant-specific subfamily of glycosylhydrolases (GH5_14) that is (i) able to cleave 1,3-β-D-glucans like callose, which is produced by GSLs, and (ii) contains an actin-binding domain^78^, forming another link between actin and GSLs. The exact role of F-actin in *Spirogyra* cytokinesis remains to be determined microscopically and through detailed experimentation. Overall, the cell division machinery of *Spirogyra* shows evidence both for deeply conserved components and lineage specific divergence.

### Division of the name-giving spiral chloroplast: absence of key division genes

Streptophyte plastids are remarkable. These genetic compartments are under particular control by nuclear genes^17,79–81^ and division is tightly regulated^82^ featuring genes homologous to bacterial cell wall (murein) biosynthesis^83,84^ inherited from the cyanobacterial plastid progenitor during the primary endosymbiosis at the root of the Archaeplastida. Intriguingly, for dividing its unique spiral plastid *Spirogyra* appears to lack almost all plastid division genes, including those coding for FtsZ (Fig. 5a). FtsZ proteins constitute the inner division ring of plastids and bacteria^85,86^. Importantly, we find that MinD and MinE proteins, that are known to regulate FtsZ in plastid division, as well as the plastid division genes ARC5, ARC6 and PARC6 are lacking in *Spirogyra* (Fig. 5a). Only one ortholog of PDV1/2 could be recovered.

**Figure 5:**
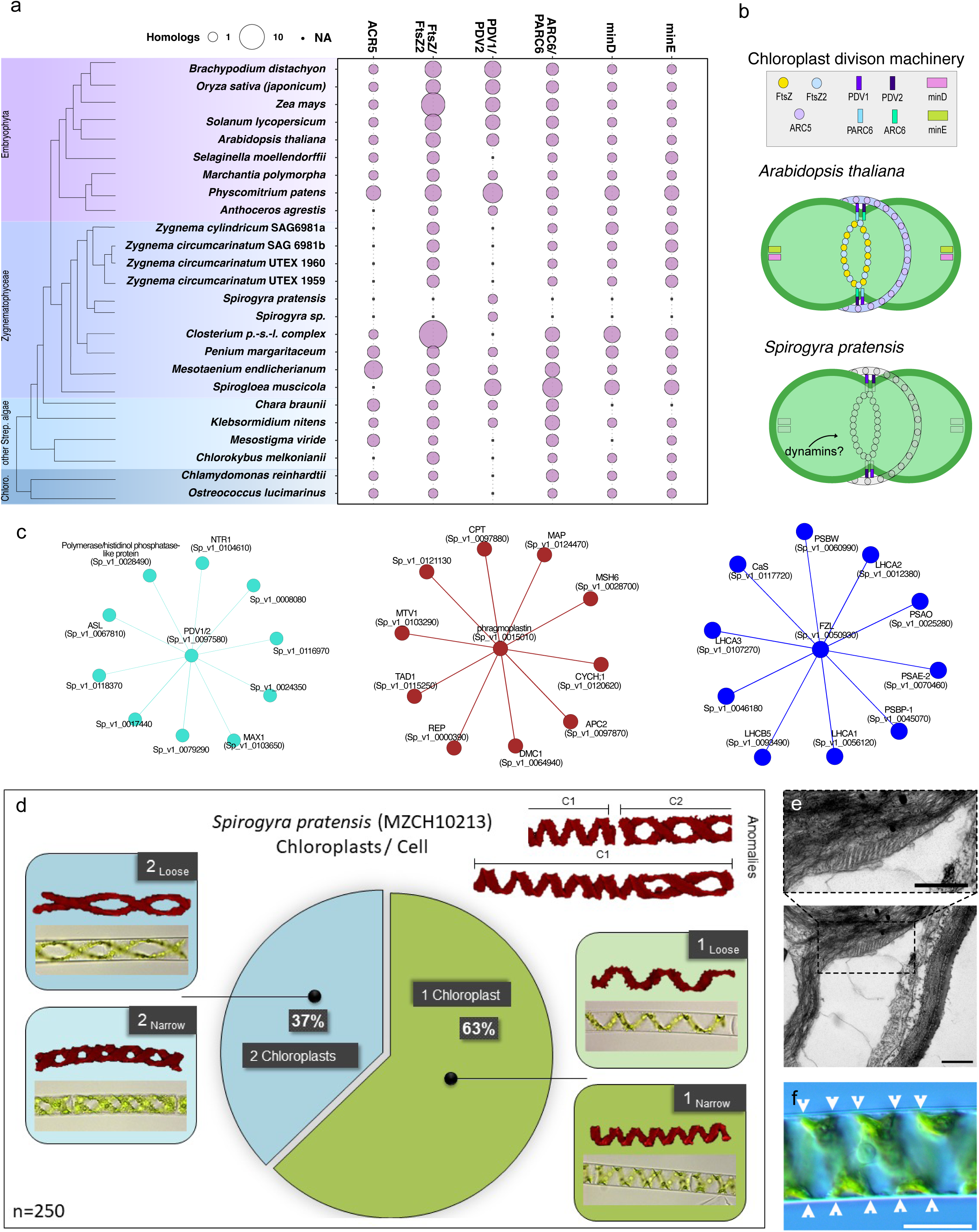
The canonical chloroplast division machinery is absent in *Spirogyra pratensis.* **a** Species tree with bubble plot showing the presence/absence of key plastid division genes in *Spirogyra pratensis* and other representatives along the green tree of life. The size of the bubbles corresponds to the size of the gene families in the respective organism. For the analysis potential orthologues were determined using BLASTP with *Arabidospis thaliana* proteins as queries. This was followed by multiple sequence alignment using mafft and construction of gene trees using iqtree2. **b** Schematic of the chloroplast division machinery with its key proteins FtsZ/FtsZ2, PDV1/2, PARC6 and ARC6 as well as minD and minE. The FtsZ proteins form an inner ring that is connected via PARC6/ARC6 and PDV1/2 to an outer ring formed by ARC5 proteins. minD and minE are localization factors that are necessary to position the the FtsZ ring. The lower panel shows the reduced chloroplast division machinery as present in *Spirogyra pratensis*. **c** Top 10 most co-expressed genes of the *Spirogyra* PDV1/2 and phragmoplastin orthologs. Annotations were transferred from a BLASTP search against *Arabidopsis thaliana* and orthogroups identified by orthofinder. Abbreviations: PDV1/2 = PLASTID DIVISION 1/2, ASL = ASYMMETRIC LEAVES2-LIKE, NTR1 = NTC-RELATED PROTEIN1, MAX1 = MORE AXILLARY BRANCHES1, CPT = CIS-PRENYLTRANSFERASE, MAP = MICROTUBULE ASSOCIATED PROTEIN, CYCH;1= CYCLIN H;1, MSH6 = MUTS HOMOLOG 6, APC2 = ANAPHASE PROMOTING COMPLEX 2, DMC1 = DISRUPTION OF MEIOTIC CONTROL 1, REP = Rab escort protein, MTV1 = MODIFIED TRANSPORT TO THE VACUOLE1, FZL = FZO-LIKE, LHC = LIGHT HARVESTING COMPLEX, PSBW = PHOTOSYSTEM II REACTION CENTER W, PSAO = PHOTOSYSTEM I SUBUNIT O, PSAE-2 = PHOTOSYSTEM I SUBUNIT E-2, CaS = Calcium sensing receptor, PSBP-1 = PHOTOSYSTEM II SUBUNIT P-1. **d** Light microscopy of *S. pratensis* chloroplasts. The majority of cells harbour a single spiral chloroplast. Chloroplasts can be observed in loose or narrow configuration. In rare cases double spiral and discontinuous forms can be found. **e** Transmission electron microscopy of *Spirogyra pratensis.* Detail of chloroplast with unidentified grid-like structure, scale bar: 500 nm. **f** Light microscopy showing the typical spirally arranged chloroplast of *Spirogyra pratensis* and possible links to the cell membrane or wall, scale bar: 20 µm.

*Spirogyra* PDV1/2 is co-expressed in the turquoise module, which is related to high light stress (Fig. 4c). Under its top 10 strongest co-expressed genes were orthologs of the cytochrome P450 MORE AXILLARY BRANCHES 1 (MAX1) involved in biosynthesis of the apocarotenoid-derived strigolactones^87^; here MAX1 probably reflects general stress-induced apocarotenogenesis that was recently shown to occur in zygnematophytes^50^ and potentially in Charophyceae that additionally stand out by expression of MAX1 homologs^88^. It further co-expresses with NTC-RELATED PROTEIN1 (NTR1), which is part of the spliceosomal complex and important for heat stress and high-light acclimation^89,90^ (Fig. 5c). Since PDV1/2 co-expresses with chloroplast signaling and transcription genes—not cytokinesis modules—it may serve a different role in *Spirogyra*. Unlike FtsZ-dependent plastid division in angiosperms, *Spirogyra* likely relies on dynamins. We identified eight dynamin-like proteins, with all Arabidopsis clades represented (Fig. S36), suggesting potential roles in plastid division (e.g., DRP5B; Fig. 5b). Six are in cytokinesis-related modules; notably, Sp_v1_0050930 co-expresses with photosystem genes (c) and likely corresponds to FZO-LIKE (FZL), involved in thylakoid morphology^91,92^. In addition, we observe that the phragmoplastin subgroup of dynamins has a clear ortholog in *Spirogyra* (Sp 0015010.1), suggesting that phragmoplast / cell plate function of *Spirogyra* cell division employs this conserved protein. This is corroborated by our co-expression data, as Sp_v1_0015010 (i) clusters in the brown module that is enriched in cytokinesis genes (Fig. 4a) and (ii) has several multiple cytokinesis and transport related genes under its top 10 strongest co-expressed genes (Fig. 5c). *Spirogyra crassa* chloroplasts are severed by the ingrowing septum during cell division^93^, what is compatible with our observation for *Spirogyra pratensis* (see Fig. 4h, Supplemental Video S1). Thus, *Spirogyra* must have a divergent mechanism of plastid division.

*Spirogyra* features large spiral chloroplasts and, intriguingly, cells of the moss *P. patens* that are subject to stoichiometric disturbances of FtsZ proteins display very large, undivided chloroplasts^94–96^. Microscopic analysis of *S. pratensis* (n=250) revealed that the majority of cells (63%) harbours only a single chloroplast (Fig. 5d). Chloroplasts can be observed in loose and narrow configuration; the number of spiral turns per chloroplast were found to be significantly different between cells with one or two chloroplasts. While cells with one chloroplast had an average of 6±1.4 turns, cells with two chloroplasts had only 2.3±0.5 turns. A linear regression analysis revealed that the number of chloroplasts is negatively correlated with the number of spiral turns (R^2^ = 0.69, *p < 0.001*, y= -3.80x + 9.84, Fig. S37). However, the calculated average surface area of the chloroplasts did not differ between cells with one or two chloroplasts (p = 0.84, Fig. S37), suggesting that they have a similar photosynthetic performance. In rare cases double spirals and discontinuity within the same cell can be observed. Taken together with the results mentioned above, chloroplast division apparently most often occurs in conjunction with cell division, yielding a single chloroplast per cell. Indeed, an iris-like closing of the cell wall can be observed during cytokinesis that might severe the chloroplasts, or is tightly regulated with its division (Supplemental Video S1). However, chloroplast division also occurs between cell divisions, yielding two chloroplasts per cell with varying shapes (Fig. 5).

## Discussion

The Zygnematophyceae are extremely diverse and have likely experienced an ancient reduction event, as evidenced by the loss of motile sperm in this lineage. Based on their sister relationship with land plants, one might naively assume that they have complex genomes that pay hommage to a large genetic complement with the last common ancestor of land plants and algae. Yet, the two thus far sequenced multicellular Zygnematophyceae—*Zygnema*^10^ and the here analyzed *Spirogyra*—have in fact small genomes. We currently assume that the lineage of Zygnematophyceae has undergone an ancient genome reduction and/or rampant gene loss^12,97^ concomitant with a streamlining of their cell biological toolkit^11,98^. Yet, *Spirogyra* has taken this loss to the extreme. While the streptophyte plastid division apparatus is largely conserved since its endosymbiotic origin, *Spirogyra* apparently can divide its chloroplast in the absence of most of the key division genes, including a loss of FtsZ. The absence of FtsZ function is further noteworthy because this filamentous protein was speculated to have a function in regulating plastid form^99^. How then is the helical shape of *Spirogyra* plastids brought about? It is noteworthy that unknown grid-like structures could be visualised by TEM (Fig. 5e), however to date we are unable to attribute their identity or function. It is possible that the shape of the plastid is not an intrinsic feature (mediated by filamentous proteins within the plastid) but is caused by attaching the plastid to some spiral entity of the membrane or cell wall (Fig. 5f)? Indeed, plasmolysis detaches the plastid from the plasma membrane and severely impairs plastid shape, even when turgor is re-established. An answer could be linked to the cellulose microfibrils of the plant cell wall, which often comprise spiral structures; it is conceivable that the spiral chloroplast of *Spirogyra* mirrors the spiral cell wall growth^100^. This is consistent with the chloroplast reorienting in older Spirogyra cells when growth stops: in many plant cells cellulose microfibrils are known to reorient when the cell has ceased elongation. The inability to re-arrange the chloroplast under high light, in contrast to other Zygnematophyceae, results in stress response under high light conditions.

*Spirogyra* is a classical experimental system. It stands out by having served as system for major discoveries in physiology and cell biology^30,31^. Transient transformation has been achieved^101^, and it remains the only filamentous Zygnematophyceae with a fully characterized life cycle in the laboratory. Thus, its potential for biotechnology cannot be understated. Our data, and comparisons across streptophytes^26,50^, show that conserved hubs might govern their biological programs (Extended Figure 1f). Now, with the genome at hand, the stage is set for using *Spirogyra* as a streamlined system for understanding zygnematophyte biology in a filamentous organism.

## Methods

### Growth of cultures

*Spirogyra pratensis* [Transeau, 1914] MZCH 10213 was obtained from the Microalgae and Zygnematophyceae Collection Hamburg^25^ (MZCH). This strain is a derivative of the strain #928 obtained from the Culture Collection of Algae at the University of Texas at Austin (UTEX)^102^. To obtain MZCH 10213 a culture of UTEX #928 was purified to remove microbial contamination. For biomass generation and maintenance, the alga was cultivated in 500 ml glass bottles (Schott) filled with 350 ml of liquid modified WHM medium^103^ and kept under sterile aeration (ca. 200 ml/min) for gas exchange and airlift. A climate cabinet (Rumed, 1602+) was used at a temperature of 25 °C and a photon flux of 30-40 µmol m^-2^ s^-1^ (fluorescent tubes); spectra are published in Dadras et al.^26^. The light-dark rhythm was 16h:8h. Prior to the transfer to solid medium for conjugation the medium was renewed twice a week.

### Developmental stages

For induction of conjugation and zygospore formation the cultivation was carried out in 92×16 mm Petri dishes on *Closterium*-Medium (C-medium)^104^ using a trace element solution according to Provasoli and Printer^105^. Medium was solidified with 1.5% agar (w/v). In half of the petri dishes the agar medium was covered by cellophane disks allowing better removal of plant material. For sufficient aeration Petri dishes were only partly sealed with two 1.5 cm strips of parafilm. Cultivation was carried out in an RUMED Typ 1602+ incubator (Rubarth Apparate GmbH, Germany) at 25 °C with a photon density of 40-50 µmol m^-2^ s^-1^ and a 16 h:8 h light-dark rhythm with a maximum duration of 31 days. Samples enriched with the following developmental stages were examined: (1) vegetative cells (no conjugating cells observed), 3 days old (2); young spores (also conjugating and non-conjugating cells present), 10 days old; (3) immature spores (also conjugating and non-conjugating cells present, on cellophane), 14 days old and (4) mature spores (also degenerating conjugating and non-conjugating cells present), 31 days. Developmental stages were microscopically analysed using an inverted microscope, Olympus, D / IX50. After harvesting at different time points plant material was transferred to 2 ml tubes and shock frozen in liquid nitrogen. Until RNA extraction the material was stored at -83°C.

### Stress experiments: Gradient table setup

The strain was cultivated in C-medium^104^ for an average of 14 days in glass flasks (SCHOTT, Germany) aerated at 80 µmol photons m^-2^ s^-1^ (spectra are published in Dadras et al.^26^). Filaments grown in 300 ml of this aerated culture were transferred in 50 ml C-medium in sterile glass tubes and fragmented with Ultra-Turrax (Typ TP 18/10, Janke & Kunkel GmbH, Germany) for 1 min. In total, 900 ml of aerated culture were used for the set-up of the experiment. 150 ml fragmented material was diluted in 1.5 l sterile C-medium. For the gradient table setup, the so-prepared algal suspension was distributed across 504 wells (42 twelve-well plates [tissue culture testplates 12 No. 92412, TPP, Switzerland]; 2.5 mL of culture per well). A 43^th^ plate was used as control and was placed in the culture room at 20°C. The plates were sealed with Surgical tape, Micropore™ tape (3M, Germany) for better gas exchange and keeping sterile condition. The 42 twelve-well plates were then placed on a table that generates a cross-gradient of temperature (8.6±0.5 °C to 29.2±0.5 °C on the x-axis) and irradiance (21.0±2.0 to 527.9±14.0 µmol photons m^-2^ s^-1^ on the y-axis). The temperature gradient was generated using a custom-made table (Labio, Czech Republic) equipped with true-daylight LEDs (sTube 2W 120 ver 11:11, Snaggi, Czech Republic, for spectrum see Dadras et al.^26^ and Fig. S18) set to a 16:8h L/D cycle (Light from 6 am to 22 pm, Central European wintertime). The *Spirogyra* biomass was exposed on the gradient table for 72 hours (for sampling for RNAseq and physiological measurements) and 89 hours (for detailed light-microscopical evaluation). The experiment was performed in triplicates.

### Stress experiments: Photophysiological measurements in tandem with the gradient table

For maximum-quantum yield measurements (*F_v_/F_m_*), the maxi version of the IMAGING PAM (ImagMAX/L, M-series, Walz, Germany) with an IMAG-K5 CCD camera, guided with the ImagingWinGigE (V2.32) software, was used. The 12-well plates containing grown *Spirogyra* biomass were dark-adapted for 10-30 min before the measurement. The measurements were performed without the lid. For the *F_v_/F_m_* measurement, a short saturation pulse (Intensity 3) was applied. The measurement settings of the IMAGING PAM were the following: measuring light 1, gain 3, damping 2, mean over AOI (area of interest) was turned off. No special SP-routine was applied to modify the signal to noise ratio of the chlorophyll fluorescence measurement.

Fv/Fm values from the gradient table experiment were analyzed in R (v4.3.1) using a Kruskal-Wallis test with Fisher’s least significant difference post-hoc test. Groups were considered significantly different at Bonferroni-adjusted P ≤ 0.001.

## Microscopy

### Differential interference contrast microscopy

Differential interference contrast (DIC) imaging was done for all replicates from the table with a Olympus BX-60 microscope (Olympus, Japan) with a JENOPTIK GRYPHAX ®PROKYON camera and the JENOPTIK GRYPHAX Software (version 2.2.0.1234) (JENOPTIK AG, Jena, Germany). The morphology of chosen conditions of *Spirogyra* cells that were 89 h on the table was analyzed.

### Bright field and confocal laser scanning microscopy

For determination of the chloroplast number and shape, cells were photographed using a Zeiss Axiovert 200 M light microscope (Carl Zeiss AG), equipped with a Axiocam HRc camera (Carl Zeiss A) and a Zeiss Axiovision software. Time-lapse imaging (Supplemental Video S1) was carried out on cells growing on agar plates containing WHM medium solidified with 0.75% agar (w/v). An Axiovert 35 inverted microscope Carl Zeiss AG) equipped with a EOS 77, 30 camera (Canon) was used to capture images every minute at a magnification of 200x or 400x. Irradiation was provided by the microscopés halogen lamp at a medium intensity.

Confocal laser scanning microscopy was performed with a Zeiss Pascal System und the control of a Zen 2009 software, excitation was generated with an argon laser at 488 nm, and emission was collected with a longpass filter (505 nm), z-stacks were generated and chloroplast autofluorescence was false coloured red according to Permann et al.^106^.

### Transmission electron microscopy

Transmission electron microscopy followed a standard chemical fixation protocol using 2% glutaraldehyde in 75 mM cacodylate buffer, pH=7, followed by 1 % OsO_4_ at 4°C, overnight according to Quader^107^ and Holzinger et al.^108^. Dehydration was achieved with a series of graded acetone concentrations (30–100%). Samples were finally embedded in epoxy resin according to Spurr^109^. Ultrathin sections were obtained with a ultramicrotome (Ultracut E, Leica-Reichert-Jung, Nußloch, Germany) and counterstained with 2% uranyl acetate followed by lead citrate^110^. Transmission electron micrographs were taken on a 906 E TEM (LEO, Oberkochen, Germany) at 100kV with a Wide-angle Dual Speed 2K-CCD camera from Tröndle Restlichtverstärkersysteme (TRS) operated by the software ImageSP-Professional + Panorama (MIA) also from TRS.

### Nucleic acid isolation

#### Genomic DNA extraction

Fresh algal filaments were harvested and sieved through a 100 µm test sieve. Residual fluids were removed by pulse centrifugation. A modified cetyltrimethylammonium bromide (CTAB) protocol of Porebski et al. (1997) was used to isolate genomic DNA. Samples were shock frozen in liquid nitrogen and ground in the presence of liquid nitrogen. 20 mL of 65°C preheated 2× CTAB extraction buffer (0.1 M Tris-base, 1.4 M NaCl, 0.02 M EDTA-2Na, 2% CTAB, 2% PVP), 40 µL ß-mercaptoethanol and 40 µL proteinase K were added. Samples were incubated at 65°C for 45 minutes with gentle mixing every 15 minutes. After cooling to room temperature (approx. 13 minutes), samples were centrifuged at 13,000 x g for 20 minutes and the supernatant was transferred to a new Falcon tube. The same volume of phenol/ chloroform/ isoamyl alcohol (∼ 20 µL) was added and centrifuged at 13,000 x g for 15 minutes. The upper layer was transferred to a new Falcon tube and the same volume of chloroform/ isoamyl alcohol was added and centrifuged for 15 minutes at 13000 x g. The upper layer was transferred into a new tube and 0.7 volume of ice-cool isopropanol (∼12 mL) was added, mixed by gentle inversion and incubated for 30 minutes at room temperature. The samples were then centrifuged for 15 minutes at 13000 x g, the supernatant was removed and the pellet was washed with 70% ethanol. After air drying (∼ 15 minutes), pellets were solubilised with 100 µL TE buffer and 10 µL RNAse stock solution (1 mg/mL) was added before incubation at 37 °C for 1 hour. After cooling to room temperature (∼ 13 minutes), samples were filled to 500 µL with TE buffer. The same volume of phenol/ chloroform/ isoamyl alcohol (∼ 20 µL) was added and centrifuged for 15 minutes at 13,000 rpm. The upper layer was transferred to a new Falcon tube, the same volume of chloroform/isoamyl alcohol was added and centrifuged for 15 minutes at 13,000 x g. gDNA was precipitated by adding 2 x volume of absolute ethanol and 1/5 volume of 5 M NaCl. Samples were gently inverted and stored overnight at -20°C. Finally, samples were centrifuged for 10 minutes at 13000 x g, the supernatant was removed and the pellet was washed with 70% ethanol. After air drying, pellets were resuspended in 100 µL TE buffer and stored at -20°C.

### RNA extraction

RNA from thallus material (control material), developmental stages and cytokinin treatments was extracted using the RNeasy ® plant mini kit (Qiagen, Germany) with minor modifications. Approximately 0.1 g of each sample was ground in liquid nitrogen. After grinding, samples were frozen in liquid nitrogen until all samples were prepared for extraction. Fresh RLC buffer was used for each extraction. The lysate was centrifuged at 18000 x g for 2 minutes. After addition of ethanol, and transfer to RNeasy Mini spin columns, samples were centrifuged for 25 seconds at 8000 x g. Same conditions were used for the RW1 and first RPE washes. After the second addition of RPE, samples were centrifuged for 2.20 minutes. 30-50 µL DEPC-treated water was used to elute RNA. DNase digestion was performed on each sample by adding 10 µL RDD buffer (a component of the RNase-Free DNase Set, used in the RNeasy Kit, Qiagen, Germany), 2.5 µL DNase I and ultrapure lab water (MilliQ) up to 100 µL. After incubation for 15 minutes at room temperature, 200 µL phenol / chloroform was added, samples were inverted and incubated for 10 minutes at room temperature followed by centrifugation at 12,000 x g for 15 minutes at 4 °C. The upper layer was transferred to a new tube, 500 µL of isopropanol was added, samples were inverted and incubated for 10 minutes at room temperature. Finally, samples were centrifuged at 12,000 rpm for 10 minutes at 4 °C, the supernatant was removed and the pellet was resuspended in DEPC-treated water before incubation at 55°C for 15 minutes. Samples were stored at -80 °C.

The quality of all gDNA and RNA samples were assessed by agarose gel electrophoresis, Nanodrop (ND-1000; Thermo Fisher) and fragment analyser (Agilent).

#### RNA extraction for gradient table experiments

After absorption measurements, the twelve-well plates were placed back on the table to let cells adjust to the table conditions again for a minimum of 5 minutes before harvesting them. For RNA extraction, 0.4 mL were taken from every well of the 42 twelve-well plates grown on the table after pipetting the cells up and down twice to homogenize them. In total, 4.8 mL liquid culture was taken per condition on the table (i.e., pooling 0.4 mL of each 12 wells per each of the 42 conditions). The samples were centrifuged for 5 min at 20 °C and 4000 rpm. The supernatant was removed and the pellet was frozen at -80 °C. To extract RNA, the GeneMATRIX Universal RNA Purification Kit (Roboklon, Cat. No. E3598, EURx Ltd., Sp. Z o.o., Poland) was used following the Plant Tissue RNA purification protocol provided by the manufacturer. For cell disruption, the frozen biomass was transferred in sterile disposable 1.5 ml homogenizer BioMasher (Nippi Inc., Japan) containing 300 µl lysis buffer and homogenized with the Pestle Motor PowerMasher II (Nippi Inc., Japan). The disruption lasted around 30-40 sec for complete destruction of cells followed by a sonication for 1 min. On-column DNAase digestion was applied for all RNA samples using DNAse I (Roboklon, Cat. No. E1345, EURx Ltd., Sp. Z o.o., Poland) following the manufacturer protocol.

### GC-MS analysis for central metabolites

For the central metabolite analysis by GC-MS, 2 mg lyophilized and homogenized algae material was extracted with 1 ml methanol/chloroform/water (129/50/25); to recover enough biomass, 4-6 adjacent samples were pooled to form nine groups for analysis. After 45 min shaking at 4°C, 0.5 ml water with allo-inositol as internal standard was added for phase separation. 450 µl of the upper phase was dried under a stream of nitrogen and derivatized with 15 µl methoxyamine hydrochloride in pyridine (30 mg ml^-1^) overnight at room temperature. Samples were silylated by addition of 30 µl N-methyl-N-(trimethylsilyl) trifluoroacetamide (MSTFA) for 1 h and analyzed by GC-MS (Agilent 7890B gas chromatograph coupled with 5977N mass selective detector) as previously described^111^. Data were analyzed with MS-DIAL^112^ 4.9.22 software together with internal and external spectral libraries^113,114^ for compound identification. Final data were visualized by VANTED 2.1 software^115^.

Differentially expressed genes (DEGs) were calculated per sample against control conditions (also see Materials and Methods section) and summarized into the same nine groups as for metabolite analysis. KEGG annotations were generated by eggNOG-mapper and mapped onto relevant KEGG pathways (map00020, map00052, map00250, map00260, map00280, map00290, map00330, map00380, map00400, map00500).

### Sequencing

#### Long read DNA HiFi sequencing

Long read WGS library was prepared with the Express Template Prep Kit 2.0 (PacBio) according to protocol version 06 (Procedure & Checklist – Preparing HiFi Libraries from Low DNA Input Using SMRTbell Express Template Prep Kit 2.0 (Version 06; June 2020)). Library preparation was started with 768ng genomic DNA in 50µl EB. The DNA was sheared using a Megaruptor2 (Diagenode) and long hydropores with settings for 30kbp aiming for ∼15kbp fragment size. Fragment size distribution was quality controlled by running a 0.1ng/µl dilution of the sheared sample on a Femto Pulse (Agilent) with FP-1002 run protocol (165 kbp). The shared sample was purified with 1X Ampure PB beads (PacBio), washed twice with 80% ethanol, and eluted in 46.4µl elution buffer (PacBio). DNA concentration was measured with Qubit dsDNA HS and 45.4µl (469ng) were used as input for the library preparation according to the manufactures’ instruction with the following modifications: End-repair reaction was incubated for 30min at 20°C. Ligation was performed overnight for 18h. Library concentration was measured with Qubit (Thermo Fischer) and size distribution was analyzed on a Fragment Analyzer (Agilent) with DNF-464 run protocol using a 1ng/µl dilution in TE. Library was size selected on a Blue Pippin (Sage Science) according to the protocol “Preparing HiFi SMRTbell Libraries from Ultra-Low DNA Input” (PacBio, Version 01; August 2020) and the following settings: 0.75% DF 3-10 kb Marker S1 – Improved Recovery, Marker S1, Cutoff 6,000-50,000bp. The size selected library was purified with AMPure PB beads (PacBio) and eluted in 11µl EB. Library concentration was measured with Qubit dsDNA HS (Thermo Fischer) and size distribution was analyzed with a Fragment Analyzer (Agilent). Sequencing primer (v5) and polymerase were bound to the library using the Sequel® II Binding Kit 2.2 (PacBio) and the library was sequenced on a Sequel II or Sequel Iie instrument using one 8M Sequel II SMRT Cell with a final on plate loading concentration of 41.5pM, 2h pre-extension and 30h movie time. Circular consensus (CCS) reads were generated with SMRT Link version 11 with min. predicted accuracy of 0.99 and min. 3 passes.

### Short read WGS sequencing

The short read WGS library was prepared with the DNA tagmentation based library preparation kit (Illumina) without PCR, with 500ng gDNA input. Library preparation was followed by a cleanup using SPRI beads (Beckman Coulter Genomics). After library preparation the library concentration was measured with Qubit (ssDNA assay kit, Thermo Fischer) and the library was then quantified using the Peqlab KAPA Library Quantification Kit and the Applied Biosystems 7900HT Sequence Detection System (Thermo Fischer). The library was sequenced on an Illumina NovaSeq6000 sequencing instrument with a paired-end 2×150bp protocol aiming for 60Gb.

### RNA-Seq

Each library was prepared using the Illumina® Stranded TruSeq® RNA sample preparation kit. Library preparation started with 100 or 500ng total RNA. After poly-A selection (using poly-T oligo-attached magnetic beads), mRNA was purified and fragmented using divalent cations under elevated temperature. The RNA fragments underwent reverse transcription using random primers. This was followed by second strand cDNA synthesis with DNA Polymerase I and Rnase H. After end repair and A-tailing, indexing adapters were ligated. The products were then purified and amplified (15 PCR cycles) to create the final cDNA libraries. After library validation and quantification (Agilent Tape Station), equimolar amounts of the library were pooled. The pool was quantified by using the Peqlab KAPA Library Quantification Kit and the Applied Biosystems 7900HT Sequence Detection System. The pool was sequenced on an Illumina NovaSeq6000 sequencing instrument with a PE100 protocol aiming for 30 million clusters per sample.

### Quality control and differential gene expression (DEG) analysis

The adapters of the paired-end RNAseq reads were trimmed using Trimmomatic (v0.39) (ILLUMINACLIP: adapter.fasta:2:30:10:2:True LEADING:26 TRAILING:26 SLIDINGWINDOW:4:20 MINLEN:36; for adapter sequences see Suppl. Table S3)^116^ and quality controlled using FastQC (v0.12.1) and MultiQC (v1.16)^117,118^. Trimmed and quality-controlled samples were pseudoaligned to the transcriptome using Kallisto. Kallisto’s length scaled transcript per million data was loaded into R (v4.3.1) using Tximport^119^ (v1.28.0), transformed to count data and aggregated to gene level. Counts per million (CPM) were calculated using edgeR^120^ (v3.42.4). Lowly expressed genes were filtered out (CPM>1 in 3 or more samples; 1111 genes were removed) and the filtered data was normalized within sample triplicates using the smooth quantile normalization of qsmooth^121^ (v1.16.6). Afterwards we used voom from the limma package^122^ (v3.56.2) to model the mean-variance relationship. To investigate outliers, we performed clustering based on Euclidian distances using the standard R stats package. Based on clustering results and FastQC scores (percentage of read alignment ≤50% and number of reads mapped ≤5 GB) we removed sample S1_37 (Fig. S19) and the corresponding replicates S2_27 and S3_37 for differential gene expression analysis. The spread of the data was investigated using principal components analysis using prcomp from the R stats package (Fig. 2d, Fig. S17, Fig. S20-S21). Differentially expressed genes were determined by comparing the remaining 41 treated samples against control condition using the limma package. Genes were considered differentially expressed at |log2Fold change|>1 and Benjamini-Hochberg corrected p≤0.01. All plots were generated using ggplot2 (v3.5.0)^123^ the R package tidyverse^124^ (v2.0.0) was used for data management. For differential gene expression analysis, RNA-seq samples obtained from the 41 conditions of the gradient table were compared to control conditions (position 29°C; 530 µmol photons s–1 m–2 had to be excluded for analysis due to poor RNA quality in one replicate) (Fig. 2e).

### Gene expression viewer

All gene expression datasets generated in this study, including Spirogyra zygospore development to vegetative filaments, responses to a gradient of light and temperature stress, cytokinin treatment, and osmotic stress experiments are available in the MAdLandExpression atlas (https://peatmoss.plantcode.cup.uni-freiburg.de). This expression atlas is the successor of PEATmoss^125^, extending the available expression data to other species of bryophytes and streptophyte algae. MAdLand Expression is based on EasyGDB^126^, and includes multiple interactive visualization methods to explore expression values, such as a line and bar plots, heatmaps, expression cards, replicates plot and a table containing the average expression values (Fig.S23), which can be downloaded in multiple formats. All available Spirogyra datasets are normalized to transcripts per million (TPM).

### Assembly and annotation

HiFi reads obtained by the sequencing process was subjected to assembly using Hifiasm^79^, available at https://github.com/chhylp123/hifiasm, for *S. pratensis* with the following command: hifiasm -o output.asm -t 40 reads.fq.gz. Preliminary assemblies were evaluated for contiguity and completeness with BUSCO^127^. We continued with the assembly with the highest score for RNA-seq mapping rate and BUSCO completeness.

### Genome size estimation using k-mer frequency

Genome size estimation for *S. pratensis* was performed by k-mer frequency analysis with the findGSE tool^128^ after counting k-mers with Jellyfish^129^.

### Repeat annotation

SSR: We used Tandem Repeats Finder (TRF) version 4.04 with two sets of parameters: 2 10 10 80 10 24 2000 (soft) and 2 3 5 80 10 20 2000 (aggressive). REPET: For de novo identification of dispersed repeats, we used the REPET package v2.5. We run the TEdenovo^130^ pipeline from REPET on the haplotype 1 assembly (parameters were set to consider repeats with at least 4 copies). We obtained a library of 218 consensus sequences that were filtered to keep only those that are found at least once as full-length copies in the assembly and we retained 149 of them. This library of consensus sequences was then used as a probe for whole genome annotation by the TEannot^131^ (REF) pipeline from the REPET. Threshold annotation scores were determined for each consensus as the 99th percentile of the scores obtained against a randomized sequence (reversed input, not complemented and masked using TRF with parameters 2 7 7 80 10 70 10). The library of consensus sequences was classified using PASTEC^132^ followed by manual curation. Phylogenetic analysis: To build the CMC clade tree, we searched for Plavaka homologs in the GenBank nr proteins database (NR) and the WGS assembly database for Rhodophyta and we retrieved CMC transposase sequences from Yuan and Wessler^36^. The transposase protein domains were aligned with MAFFT v7.475 and the phylogenetic trees were constructed with IQ-TREE version 1.6.12 (-m MFP+MERGE -bb 1000). To build the Gypsy tree, homologs of the *S. pratensis* RT sequences were searched against the whole NR database and filtered versions thereof to target or exclude the putative HGT donor (Mucoromycotina and SAR) and recipient clades (Viridiplantae below Tracheophyta). The sequences were then processed as above with MAFFT and IQ-TREE.

Satellite DNA repeat 128 bp arrays were annotated with TideCluster (https://github.com/kavonrtep/TideCluster). Average array size and inter-array spacing were calculated for all arrays (>500 bp).

### Contamination analysis

MEGAN6 analysis^133^ was conducted to investigate contamination in the genome of *S. pratensis* by initially splitting the genomic scaffolds into 1 kbp fragments which were queried against the NCBI non-redundant (nr) protein database (version date : 2023-07-28) using Diamond^134,135^ with an e-value threshold of 0.1 on European Galaxy Interface^136^. The resulting output was imported into MEGAN6 for analyzing taxonomic distribution according to NCBI taxonomy. Parameters of assignment to the last common ancestor (LCA) were set to default (minscore: 50; top percent: 10). Finally, scaffolds were summarized based on the percentage taxonomic designation of their contributing fragments to detect putative contaminants. Scaffolds with a percentage contribution greater than 0,5% in the case of bacteria, greater than 1.5% (fungi and SAR group) were analysed as follows. The distribution of the hits was visualized along the length of the scaffold using plotly package in R^137^; if clusters of the same taxonomic assignment were found the underlying region (Table S5, columns: S-U) was subjected to BLASTx and BLASTn searches and the results analysed. All data are summarized in Table S5; suspicious taxonomic contribution was detected primarily from fungi and SAR group organisms, in one case from bacteria. Analysis of BLAST results showed potential overlap with regions detected by the REPET analysis (Table S5, Column: U). After hard masking of scaffold using ‘N’ characters, BLAST searches of suspicious regions were repeated. In most of the cases, either top hits were detected from Streptophyta or no hits remained. Hence, no scaffolds or scaffold regions were detected to be due to contamination.

Contamination and repeat analyses demonstrate evidence for sequences closely related to SAR group/oomycetes. To distinguish a potential HGT event of the Spirogyra Gypsy from a contamination / endogenous oomycete, we performed PCR using oomycete-specific ITS primers for a locus found on *scaffold_20* region 4,500 to 4,810 (311 bp).

### Structural gene annotation

Structural gene annotation was performed using REAT^138^ (v0.6.1) and Minos^139^ (v1.8.0). We used the transcriptome, homology and prediction workflow of REAT to generate diverse sets of gene models that were consolidated into a final set using Minos. As a final step in both pipelines and at the end of all three REAT workflows, Mikado (v2.3.4) was used for scoring and picking gene models. The transcriptome workflow used Hisat2 (v2.2.1)^140^ to align short reads, Scallop^141^

(v0.10.5) and Stringtie^140^ (v2.1.5) to assemble transcripts, as well as Portcullis^142^ (v1.2.4) to determine splice junctions. It also uses Prodigal^143^ (v2.6.3) to call ORFs and Diamond^134,135^ (v2.1.8) to align gene models to homologous proteins. For the transcriptome workflow we used 41 short read RNA-seq samples that included different stress conditions and developmental stages (see Suppl. Table S2). The homology workflow uses Spaln^144^ (v2.4.7) to align cDNA and homologous proteins to the *Spirogyra* reference genome. For this workflow, we utilized available protein and annotations of representative streptophyte species: *Anthoceros agrestis* oxford^145^, *Arabidopsis thaliana* (v11)^146^, *Azolla filiculoides* (v1.1)^147^,*Chara braunii* ^148^, *Mesotaenium endlicherianum* (v2)^8,26^, *Marchantia polymorpha* (v6.1) ^149^, *Physcomitrium patens* (v3.3) ^150^, *Spirogloea muscicola*^8^ and *Zygnema circumcarinatum* SAG698-1b^10^. As there was no obvious choice for the SPALN species-specific parameter set, we ran the workflow three times with the parameters for Angiosp, Chlorspc and MossWorts (also compare^26^).

In the prediction workflow different *ab-initio* predictors were run, including Augustus^151^ (v3.4.0), Glimmer^152^ (v3.0.4), SNAP^153^ (v 2013_11_29) and CodingQuarry^154^ (v2.0). In total there were four runs of Augustus, three of which included differently weighted homology and RNAseq evidence, while repeat information was provided to all runs. The output of the different prediction tools was compared and combined using EVidenceModeler^155^ (v1.1.1). The gene model sets from transcriptome, homology and prediction workflow were consolidated into a final set using the Minos pipeline and afterwards filtered for repeat elements. Minos scores the input gene models and picks the best one based on external metrics that include: Kallisto^156^ (v0.44.0) transcript quantification, homology information using Diamond^134,135^ (v0.9.34), coding potential calculated by CPC2 (v2.0)^157^ and evaluation via BUSCO^127^ (v5.5.0). Repeat related gene models were determined by providing the predicted repeat elements file to Minos, which tagged them as repeat related using default parameters (>20% overlap with repeat elements). After tagging, we filtered them out. The nomenclature of gene models reflects their distance on the chromosomes, e.g. Sp_v1_0010010 and Sp_v1_0010020 are located next to each other (leaving a gap of 9 to accommodate future intervening models). Genome completeness was assessed using BUSCO (v5.5.0) using the viridiplantae_odb10 as reference.

### Functional annotation, homology assignment, domain-based and phylogenetic analyses Protein Sequence Data Sources for comparative analyses

The representative protein sequences of Zygnematophyceae (*Mesotaenium endlicherianum*, *Spirogyra pratensis* UTEX 928*, Zygnema circumcarinatum* SAG 698-1b, UTEX 1559, UTEX 1560, *Zygnema cf. cylindricum* SAG 698-1a*, Closterium sp.* NIES-68 (minus strain), *Penium margaritaceum),* additional streptophyte algae (*Chara braunii, Klebsormidium nitens, Mesostigma viride, Chlorokybus atmophyticus, Spirogloea muscicola),* bryophytes *(Marchantia polymorpha, Physcomitirium patens, Anthoceros agrestis),* vascular plants *(Brachypodium distachyon, Oryza sativa* spp. japonica*), Zea mays, Solanum lycopersicum, Arabidopsis thaliana, Selaginella moellendorffii)* and chlorophytes *(Chlamydomonas reinhardtii, Ostreococcus lucimarinus)* were downloaded from the Genome Zoo database^40^. Details are available in Table S15.

### Analysis of cell division genes

To compare the mode of cell division between streptophyte algae and land plants. We compiled the list of 288 Arabidopsis genes that are involved in cytokinesis (Table S9), focusing on genes required for phragmoplast and PPB function. we conducted reciprocal BLASTP searches (using version 2.12.0+) against *S. pratensis*, in addition to other streptophyte algae and land plants. The e-value threshold was set to 1e-04. Subsequently, the BLAST results were filtered according to Rost^158^ to retain only homologous sequences. The top hit was determined based on the bit-score.

The presence of plastid division genes was checked by using BLASTP with *Arabidopsis thaliana* queries against the proteomes of several viridiplantae species (see Protein Sequence Data Sources/Comparative genome analysis). Afterwards, sequences were aligned using mafft and phylogenetic trees were calculated using iqtree2 (-nt 5 -st AA -m TEST -msub nuclear -alrt 1000) (Fig. S30 – S35). Members of gene families were counted manually.

### FtsZ and dynamin

To identify the FtsZ proteins in *S. pratensis,* the protein sequence of plastid division protein FtsZ 2-2 precursor (CAB76387.1) from *Physcomitrium patens*^159^ was used as query in BLASTp and tBLASTn searches against the complete set of *S. pratensis* proteins and genome assembly, respectively. At the protein level, no significant hits were detected using a threshold of ≥30% sequence identity and ≥80 alignment length. Similarly, the TBLASTN search against the whole genome did not yield any matches. Since, FtsZ proteins are characterized by presence of Tubulin/FtsZ family, GTPase domain (PF00091) and FtsZ family, C-terminal domain (PF12327) all *S. pratensis proteins* were also screened with the HMMER software suite (v3.3.2) for using the gathering threshold (-cut_ga). No proteins were detected that contained both of these domains in *S. pratensis.* However, we identified six proteins containing GTPase domain (PF00091), we assume these proteins are not classified as FtsZ but as GTPases.

To identify dynamin-like/related proteins, the TAPScan v4 script was used to assign proteins to the dynamin protein family, characterized by the presence of the DYN1 domain^160^. Sequences were aligned using mafft and phylogenetic tree were computed using IQ-TREE^161^ 2 (-nt AUTO -s FILE -m MFP -bb 1000) and converted to unrooted using ITOL^162^ and annotated manually.

### Transcription associated proteins

TAPScan^40^ v4 was performed to investigate the evolution of transcription-associated protein (TAP) gene families in *S. pratensis* comparted to other streptophyte algae (Fig. S5). The resulting gene family counts were then used to calculate lineage-specific expansion and contractions of TAP families across species. For that, A species phylogeny tree was pruned from 1KP tree^3^ to include only species used in this study, then rooted on the chlorophyte node, and subsequently converted into ultrametric tree using the ape^163^ package in R. Gene family evolution analyses were conducted using CAFE^41^ v.5 with default (base) parameters. To avoid the potential bias in estimation of gene birth and death rates, TAP families with zero members across all species were excluded from the analysis.

Profiles for TAP domains PF00072 (Response-reg), DEAD(PF00270), Helicase_c (PF00271) and PP2C (PF00481) were obtained from the Interpro database. We used HMMER^164^ using default parameters and 25 proteomes of representative viridiplantae species as input (see Protein Sequence Data Sources/Comparative genome analysis) to obtain protein motif IDs. For the WD40 motif tree, a reduced set of species that only contained *Arabidopsis*, *Physcomitrium* and Zygnematophyceae species was used. Motif IDs were translated into sequences using esl-reformat from easel. Sequences were aligned using mafft and phylogenetic trees were computed using iqtree2 (-nt 5 -st AA -m TEST -msub nuclear -alrt 1000). Phylogenetic trees were analyzed using ITOL^162^ and annotated manually.

### CAZymes analysis

Carbohydrate-active enzymes were annotated using the dbCAN3 web server^165^ which annotates CAZymes by using three tools: HMMER search against the CAZy domain database (*E* value < 1e-15, coverage > 0.35), DIAMOND search against the CAZyme sequence database (*E* value < 1e-102) and HMMER search against the dbCAN-sub database of putative CAZy substrates (*E* value < 1e-15, coverage > 0.35). A majority vote of the respective CAZyme family assignment was performed. Furthermore, homologs of some cell wall related enzymes (BS1/DARX1, DUF231, DUF579, QUA2/TSD2, QUA3, CGR2, CGR3) were identified using BLASTP with the characterized sequences as query (*E* value of 1e-7) against the protein files as database. For CAZyme families with described cell wall biosynthetic or remodelling functionality, *Spirogyra* sequences were compared with a set of Viridiplantae species. For that, protein sequences of those families were aligned using MAFFT (in L-INS-i mode). IQTREE2 (-st AA -m TEST -bb 1000 -alrt 1000) or FASTTREE (LG+CAT, version 2.1.10) were used for inference of maximum-likelihood trees (see Table S27, Fig. S38-S58).

### WGCNA and BLAST annotation

Weighted genes co-expression network analysis was performed in R with the WGCNA package^166^ v1.73) using qsmooth normalized and log2 transformed count data from the RNA-seq samples of the gradient table experiment. We calculated the co-expression modules using the dynamic tree cut algorithm. We chose a signed network, “bicor” as correlation function, maximum portion of outliers as 0.05, a minimum module size of 30 and a soft threshold β =14 (Fig. S24). After manual investigation, we chose a merging cut-off of 0.25 (Fig. S25). We investigated the relationship between experimental parameters, gene significance and eigengenes of the co-expression modules to test how the modules relate to our experimental parameters (Fig. S26 - S29). Hub genes were determined by ranking the intra-modular connectivity (kWithin) and choosing the top 20 as cut-off. Networks were plotted using the R-package igraph^167^ (v 2.1.2) and the top 20 hub genes were highlighted and annotated with the results of a whole proteome best BLASTP analysis against *Arabidopsis thaliana* (TAIR 11) (BLASTP, default parameters). The annotation of hubs that are mentioned in the manuscript were confirmed with orthofinder results (see below); the annotation from the best BLASTP hit was used if the identified protein fell into the same orthogroup as the *Spirogyra* query. To extract the top 10 co-expressed genes of specific gene IDs the Topological Overlay Matrix calculated by the WGCNA package was exported into weighted edge lists by module using the exportNetworkToCytoscape function. Afterwards the edge weights connected to the queried gene ID were ranked and the top 10 edges with the highest weight were plotted using igraph.

### Identification of cytokinesis modules

Cytokinesis modules were identified from the WGCNA data by testing the modules for enrichment in cytokinesis-related genes. *Spirogyra* cytokinesis-related genes were identified by querying the orthofinder results with the aforementioned set of 288 cytokinesis genes (Table S9) and extracting the *Spirogyra* homologs from the shared orthogroups. We tested for enrichment using Pearson’s χ2 test (p<0.05, df=1, Suppl. Table S24).

### Orthofinder

We ran orthofinder^168^ to perform a genome wide gene family analysis in *Spirogyra*. For this we ran orthofinder twice, first without and then with a species tree as input using the following commands, run1: orthofinder.py -t 100 -a 6 -y -n run_2 -ft /Results_run_1 -s SpeciesTree_input.txt; run2: orthofinder.py -f /primary_transcripts -S diamond -M msa -A mafft - T fasttree -t 200 -a 6 -y -n run_1. Orthofinder was run with primary transcripts that were extracted using the supplied primary_transcripts.py.

### Functional enrichment

GO terms for *Spirogyra* were generated using eggNOG-mapper^169^ (v2.1.12) (emapper.py -m diamond --itype proteins --dmnd_iterate yes --dbmem --evalue 1e-10 --sensmode ultra-sensitive --tax_scope 33090 --dmnd_db /eggnog-mapper-data/viridiplantae.dmnd) and InterProScan^170^ (v 5.67-99.0) that were merged for further analysis. Using clusterProfiler^171^ (v4.8.2) and the enricher function, we performed GO term enrichment and biological theme comparison. We used different set of GO terms for the analysis: for the developmental analysis and WGCNA analysis the full set of GO terms was used (BP, MF, CC), while for analysis of the DEGs of the environmental gradient table a reduced set of GO terms was used to reduce the overwhelming signal of chloroplast related terms. For the reduced set we filtered for only BP terms and removed redundant terms using the simplify function of the clusterProfiler package.

### Differential expression analysis of orthogroups

To investigate evolutionary conservation of differentially expressed genes in stress responses, we used available RNA-seq data of the alga *Mesotaenium endlicherianum* that was generated using the same gradient table set-up and the same experimental conditions (see also Dadras et al.^26^). The raw reads were trimmed, quality controlled and DEGs were determined (refer to DEG analysis for tools and parameters). Compared to the *Spirogyra* experiment, there was no external control for the *Mesotaenium* dataset. Therefore, we used the samples grown at 21°C and 20 µmol m^-2^ s^-1^ of the gradient table as control. Differentially expressed genes from *Spirogyra* and *Mesotaenium* were annotated with orthogroups derived from the orthofinder run. Per experimental condition, the differentially expressed orthogroups between the two experiments were analysed and plotted using R.

### Jaccard distances and hub gene conservation

We used the results of the WGCNA analysis of Dadras et al.^26^ which was obtained under the same experimental conditions, and compared them to our WCGNA modules on orthogroup level. Similarities between modules were calculated using Jaccard similarity and clustered using the R-package pheatmap (v 1.0.12) and euclidian clustering. For the hub gene conservation analysis, we mapped the *Spirogyra* hub genes to *Mesotaenium* genes on orthogroup level and sorted them based on their intramodular connectivity rank. If the orthogroup contained multiple *Mesotaenium* genes, we continued with the *Mesotaenium* gene with the highest intramodular connectivity. The intramodular connectivity rank was normalized with the module size to obtain the top % rank.

### Detection of duplication events using double trouble

To classify paralogs into duplication types, doubletrouble tool^42^ was performed using the longest peptide isoform for each gene with default parameters for all the strephytophyte algae. doubletrouble classifies genes into segmental (SD), tandem (TD), proximal (PD), transposon-derived (TRD), and dispersed duplications (DD) (Fig. S10).

## Data availability

Raw data and assembly generated in this study were deposited into the NCBI SRA with the BioProject ID PRJNA1110855. The annotation is available via https://phycocosm.jgi.doe.gov/Spipra1/Spipra1.home.html. Gene expression data are available in MAdLandExpression (https://peatmoss.plantcode.cup.uni-freiburg.de).

## Acknowledgements

This project was carried out in the framework of MAdLand (https://madland.science, DFG priority programme 2237), SAR (RE-1697/18-1, 19-1 and 20-1; CharKeyS) and JdV (422691801, 440231723, 514060973, 528076711) are grateful for funding by the DFG; KvS and HZ acknowledge DFG funding within the CharMod project Schw687/13. JdV. thanks the European Research Council for funding under the European Union’s Horizon 2020 research and innovation programme (Grant Agreement No. 852725; ERC-StG “TerreStriAL”). Computational support of the Zentrum für Informations-und Medientechnologie, especially the HPC team (High Performance Computing) at the Heinrich Heine University is acknowledged and the Gesellschaft für wissenschaftliche Datenverarbeitung mbH Göttingen (GWDG) for providing excellent computational infrastructure. This work was supported by the DFG Research Infrastructure West German Genome Center (project 407493903) as part of the Next Generation Sequencing Competence Network (project 423957469). NGS analyses were carried out at the production site Cologne (Cologne Center for Genomics (CCG)) and the production site Düsseldorf (Genomics & Transcriptomics Labor (GTL)). This research was funded in part by the Austrian Science Fund (FWF) 10.55776/P34181 to AH. The authors thank Katrin Schnoor, Vera Schwekendiek and Atiqur Rahaman (University Hamburg) for support in an early phase of the project. The authors further acknowledge help of Frank Friedrich with TEM and thank Luc Guder for the time-lapse imaging of cell division and spore formation (both University of Hamburg). Julian Obwegeser, University of Innsbruck is thanked for help in data generation (chloroplast counting). We are grateful to M. Lorenz and T. Friedl for supporting us with access to the facilities of the Department of Experimental Phycology and SAG Culture Collection of Algae. E.S.G. and A.D. are grateful for being supported through the International Max Planck Research School (IMPRS) for Genome Science. NFP is grateful for funding by MICIU/AEI/10.13039/501100011033, ERDF/EU, and CSIC (PID2021-125805OA-I00 and JAE-PRE23-15). FM benefits from the support of Saclay Plant Sciences-SPS (ANR-17-EUR-0007) and the PlantBioinfoPF platform. To ensure data FAIRness, we utilized the DataPLANT personal assistance network and its tool stack, centered around the DataHUB^172^.

## EXTENDED DATA FIGURE LEGENDS

**Extended Data Figure 1.**
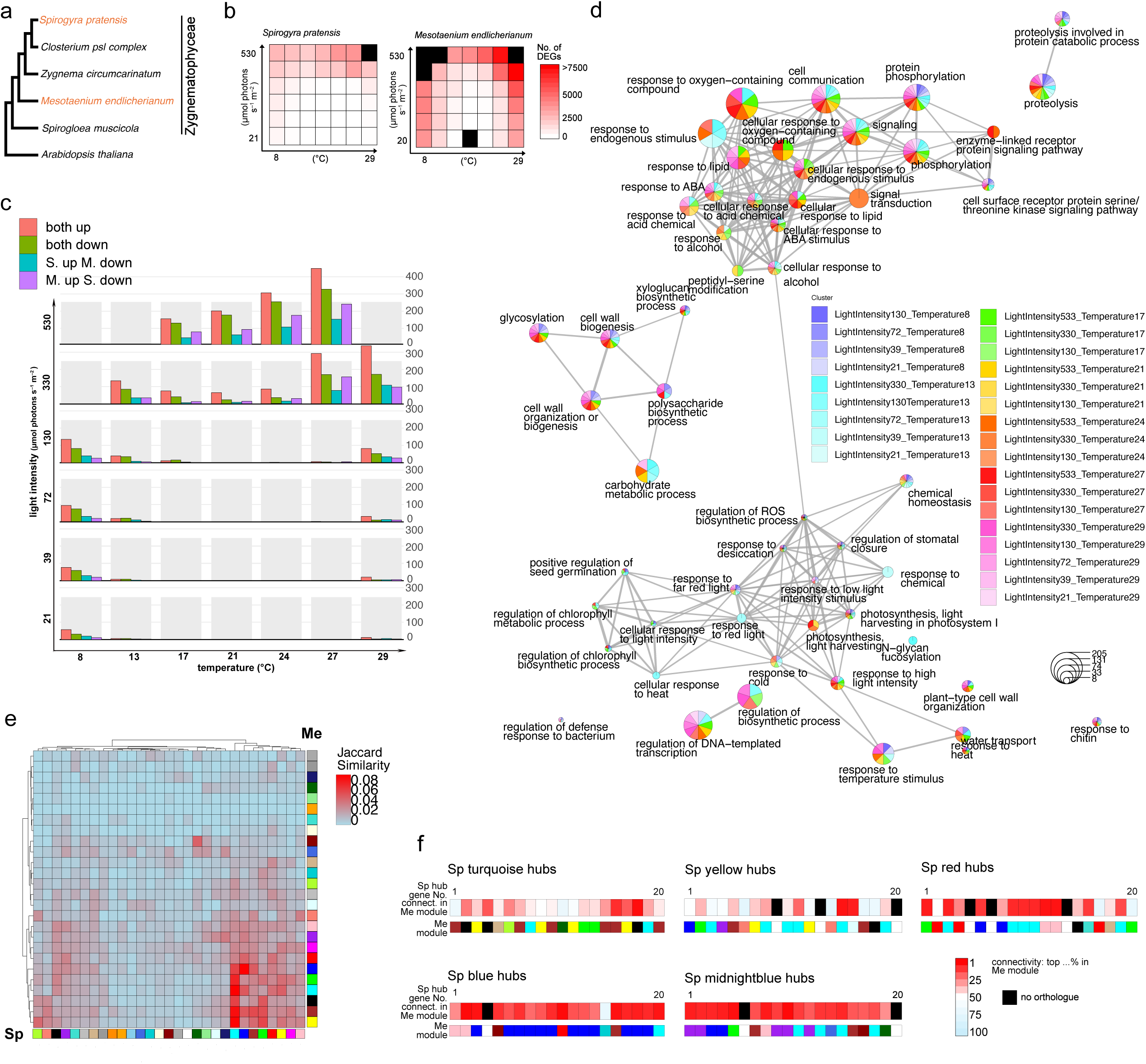
Shared stress hubs of zygnematophytes. **a** The phylogenetic relationship of the investigated Zygnematophyceae species *Spirogyra pratensis* and *Mesotaenium endlicherianum*. **b** Heatmap showing the number of DEGs of *Spirogyra pratensis* and *Mesotaenium endlicherianum* treated with the same environmental gradient table set-up. Black squares: no DEGs could be determined due to poor RNA quality. **c** Number of orthogroups (OGs) that are shared in the DEGs of both alga. Orthogroups were determined using orthofinder. **d** “Biological theme comparison” of “biological process” GO terms of shared differentially expressed orthogroups. Redundant GO terms were summarized using the “simplify” function of the R-package clusterProfiler. **e** Jaccard similarity comparing the WGCNA modules from *Spirogyra* of this study with the WGCNA modules of *Mesotaenium* recovered in the analysis of Dadras et al. 2023. **f** Conservation of hub genes between the analyses. The top 20 hub genes recovered in the *Spirogyra* modules “turquoise”, “yellow”, “red”, “blue” and “midnightblue” were tested for their connectivity within the *Mesotaenium* WGCNA analysis. Genes were compared on orthogroup level and the *Mesotaenium* gene with the highest connectivity measure from that orthogroup was chosen. Connectivity within *Mesotaenium* modules was normalized by dividing the connectivity rank by the module size.

**Extended Data Figure 2.**
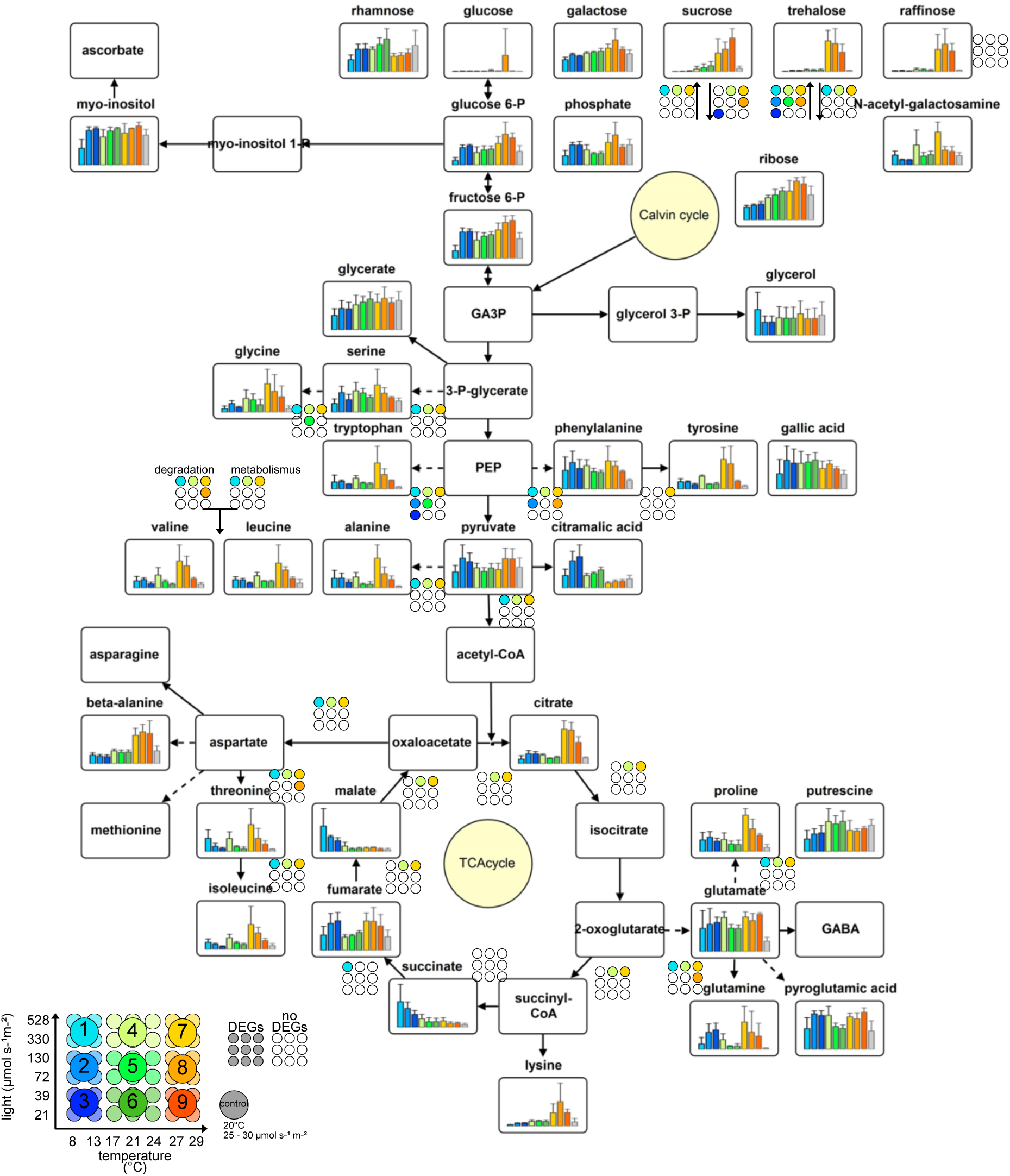
Central Metabolite analysis of gradient table experiment using GC-MS. Four to six samples were pooled into nine groups to recover enough biomass for GC-MS analysis. The bar graphs show the mean relative peak area (peak area/peak area of internal standard) of three biological replicates. Whiskers symbolize the standard deviation. Differentially expressed genes (DEGs) were calculated per sample against control conditions and summarized into the same nine groups. KEGG annotations were generated by eggNOG-mapper and mapped onto relevant KEGG pathways (map00020, map00052, map00250, map00260, map00280, map00290, map00330, map00380, map00400, map00500). DEGs are marked, if one or multiple genes in the conversion step(s) were differentially regulated compared to control conditions.

