## Supplemental Material for "The *Spirogyra* genome: signatures of shared and divergent division and differentiation"

**SUPPLEMENTARY MATERIAL, Goldbecker et al.**

**Supplementary Results and Discussion** Conserved molecular responses in two divergent Zygnematophyceae species

**Supplementary References** References of Supplementary Results and Discussion

**Supplementary Figures**

**Supplementary Figure S1** K-mer-based genome size estimation and assembly statistics.

**Supplementary Figure S2** BUSCO evaluation of the *Spirogyra pratensis* annotation.

**Supplementary Figure S3** Phylogenetic trees of Plavaka transposable elements.

**Supplementary Figure S4** Phylogenetic tree of the Gypsy reverse transcriptase domain showing a putative horizontal gene transfer.

**Supplementary Figure S5** TAP bubble plot.

**Supplementary Figure S6** Zoom into Resp_reg motif tree clade containing expansions in Zygnematophyceae.

**Supplementary Figure S7** Zoom into Resp_reg motif tree showing the clade of type-A RRs.

**Supplementary Figure S8** Zoom into WD40 motif tree clade containing expansions in Zygnematophyceae.

**Supplementary Figure S9** DEAD-box and Helicase C motif trees.

**Supplementary Figure S10** Gene duplication events.

**Supplementary Figure S11** Summary of the glycosyltransferases (GTs) found in Spirogyra.

**Supplementary Figure S12** Summary of the glycosylhydrolases (GHs) found in Spirogyra.

**Supplementary Figure S13** Summary of the carbohydrate-binding modules (CBMs).

**Supplementary Figure S14** Summary of the enzymes with auxiliary activity (AAs), pectate lyases (PLs) and carbohydrate esterases (CEs) found in Spirogyra.

**Supplementary Figure S15** Summary of CAZymes found in Spirogyra.

**Supplementary Figure S16** Light microscopic images of S. *pratensis* MZCH#10213 life cycle stages

**Supplementary Figure S17** Principal component analysis of RNA-seq data of developmental stages.

**Supplementary Figure S18** Light Spectrum of the gradient table set-up.

**Supplementary Figure S19** Hierarchical clustering of gradient table RNA-seq samples.

**Supplementary Figure S20** Small multiple PCA, impact of light intensity on RNA-seq data, gradient-table experiment.

**Supplementary Figure S21** Small multiple PCA, impact of temperature on RNA-seq data, gradient-table experiment.

**Supplementary Figure S22** Phylogeny of glycosyltransferases from family 41.

**Supplementary Figure S23** *S. pratensis* gene expression atlas at MAdLandExpression, showing expression of five genes along the temperature and light intensity gradient experiment.

**Supplementary Figure S24** Picking a soft threshold power based on scale free topology and mean connectivity.

**Supplementary Figure S25** Gene Cluster Dendrogram and Module Membership.

**Supplementary Figure S26** Gene significance for all genes split into WGNCA modules in relation to temperature.

**Supplementary Figure S27** Gene significance for all genes split into WGNCA modules in relation to Fv/Fm.

**Supplementary Figure S28** Gene significance for all genes split into WGNCA modules in relation to light intensity.

**Supplementary Figure S29** Module-trait relationship of WGCNA.

**Supplementary Figure S30** MinD phylogenetic tree.

**Supplementary Figure S31** MinE Phylogenetic tree.

**Supplementary Figure S32** ARC5 phylogenetic tree.

**Supplementary Figure S33** FtsZ phylogenetic tree.

**Supplementary Figure S34** PDV phylogenetic tree.

**Supplementary Figure S35** ARC6 and PARC6 phylogenetic tree.

**Supplementary Figure S36** Dynamin tree (unrooted).

**Supplementary Figure S37** Quantification of spiral turns and surface area of *Spirogyra* chloroplasts.

**Supplementary Figure S38** CAZyme (DUF231) ESK1 and MOAT1-4 tree.

**Supplementary Figure S39** CAZyme (GT47) XGD1, IRX7/F8H, IRX10/XYS1 and ARAD1+2 tree.

**Supplementary Figure S40** CAZyme (GT2) CesA, CsID, CsIQ, CsIB/E/G, CslO, CsiP, CsIK, CsIA, CsIL, CsIC and CSiN tree.

**Supplementary Figure S41** CAZyme (GH3) BXL1/BXL4 tree.

**Supplementary Figure S42** CAZyme (GT8) IRX8, GAUT8, GAUT1/7, PARVUS, GUX1-5 tree.

**Supplementary Figure S43** CAZyme (DUF579) IRX8, GAUT8, GAUT1/7, PARVUS, GUX1-5 tree.

**Supplementary Figure S44** CAZyme (GH31) AXY3 tree.

**Supplementary Figure S45** CAZyme (GT106) tree.

**Supplementary Figure S46** CAZyme (GT77) tree.

**Supplementary Figure S47** CAZyme (GT92) GALS1-3 tree.

**Supplementary Figure S48** CAZyme (GH27) AGAL2-3, APSE tree.

**Supplementary Figure S49** CAZyme (GT29) SIA1-2, GALT29A tree.

**Supplementary Figure S50** CAZyme (GT37) FUT1-9 tree.

**Supplementary Figure S51** CAZyme (GH79) GUS2 tree.

**Supplementary Figure S52** CAZyme (GT31) HPGT1-3, GALT31A and GALT2-6 tree.

**Supplementary Figure S53** CAZyme QUA2 and QUA3 tree.

**Supplementary Figure S54** CAZyme (GT34) XXT1-5 and GMGT1 tree.

**Supplementary Figure S55** CAZyme (GT14) GlcAT14A-E tree.

**Supplementary Figure S56** CAZyme (GT43) tree.

**Supplementary Figure S57** CAZyme (GT61) XAT/XAX tree.

**Supplementary Figure S58** CAZyme (GT116) RGGAT1 tree.

**Supplementary Results and discussion**

**Conserved molecular responses in two divergent Zygnematophyceae species**

We compared the responses of *Spirogyra* with another member of the Zygnematophyceae algae, the single celled *Mesotaenium endlicherianum*, that was subjected to the same bifactorial gradient setup^1^. We re-analysed the RNA-seq data of *Mesoteanium*^1^ to calculate DEGs and used the sample at 21 °C and 20 µmol photons s^–1^ m^–2^ as control. Despite physiological patterns suggesting that both algae prefer low light and warmer temperatures^1^, we observed major differences in the RNA-seq data: *Mesotaenium* had twice the number of DEGs than in *Spirogyra* (Extended Fig. 1a). To find conserved responses we compared the expression of orthogroups (OGs) between samples of the same condition (Extended Fig. 1b) and performed biological theme comparison (Extended Fig. 1c). Most shared differentially regulated OGs occurred under high light and high temperatures, as these also yielded the most DEGs in both experiments; GO terms associated with signaling like “cell communication” “signaling” and “response to ABA” were enriched in the shared differentially expressed orthogroups (Fig. 6d). Also GO terms connected to light sensing, photosynthesis and temperature sensing were enriched in both algae. Additionally, we found cell wall related terms like “cell wall biogenesis” or “glycosylation”. Two GRAS-type orthogroups (OG0009880 and OG0000447) were differentially expressed under high-light and heat in both algae: OG0009880 was downregulated, while OG0000447 was upregulated (Table S25). Stress signaling programs appear mediated by the same orthologs and upon the same environmental cues in the more than 500 million-years-divergent species.

We used Jaccard similarities and compared orthogroups between the co-expression modules of *Spirogyra* and *Mesotaenium*^1^. Spirogyra's turquoise module, linked to RNA metabolism and high-light stress, closely resembles at least six Mesotaenium modules, suggesting conserved processes split across modules. Its blue module, associated with photosynthesis and signaling, aligns with Mesotaenium's blue module. Many hub genes in Spirogyra’s blue module are also highly connected and recur in Mesotaenium’s blue module (Fig. 6e). These included orthologues of the calcium sensor CAS^2^ and the LRR-kinase hub described above. Among the highest connected *Mesoteanium* orthologs of blue hubs was a member of the ATP-BINDING CASETTE TRANSPORTER subfamily G (ABCG5) that radiated in land plants and of which many are involved in forming protective cuticle layers in response to stressors^3^. ABCG5 belonged to the top 2% most connected genes in the blue module of *Mesotaenium* hinting at an important and highly conserved function in both algae. Also, the midnightblue hubs of *Spirogyra* had a high connectivity within the *Mesotaenium* modules. Here, expectedly orthologues of the cell cycle regulators CYCB2;3 and CDKB1;2 were among the top 2 and 6% connected genes in *Mesotaeanium*. Additionally, we observed the midnightblue hub gene belonging to the MATE efflux carrier family and the hub gene GOLGI NUCLEOTIDE SUGAR TRANSPORTER 4 (GONST4) that transports GDP-L fucose in *Arabidopsis*^4^ among the highest connected orthologues in *Mesotaenium* (3 and 6% respectively).

**Supplementary Figures**

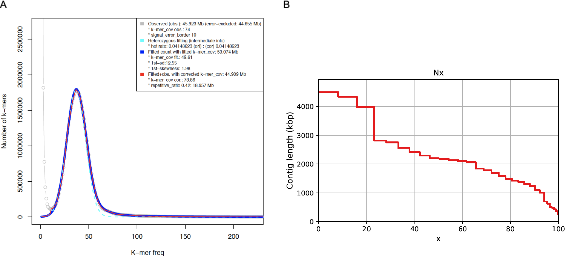

**Supplementary Figure S1: Assembly statistics of the *Spirogyra* genome.** (a) K-mer-based genome size estimation and (b) assembly statistics.

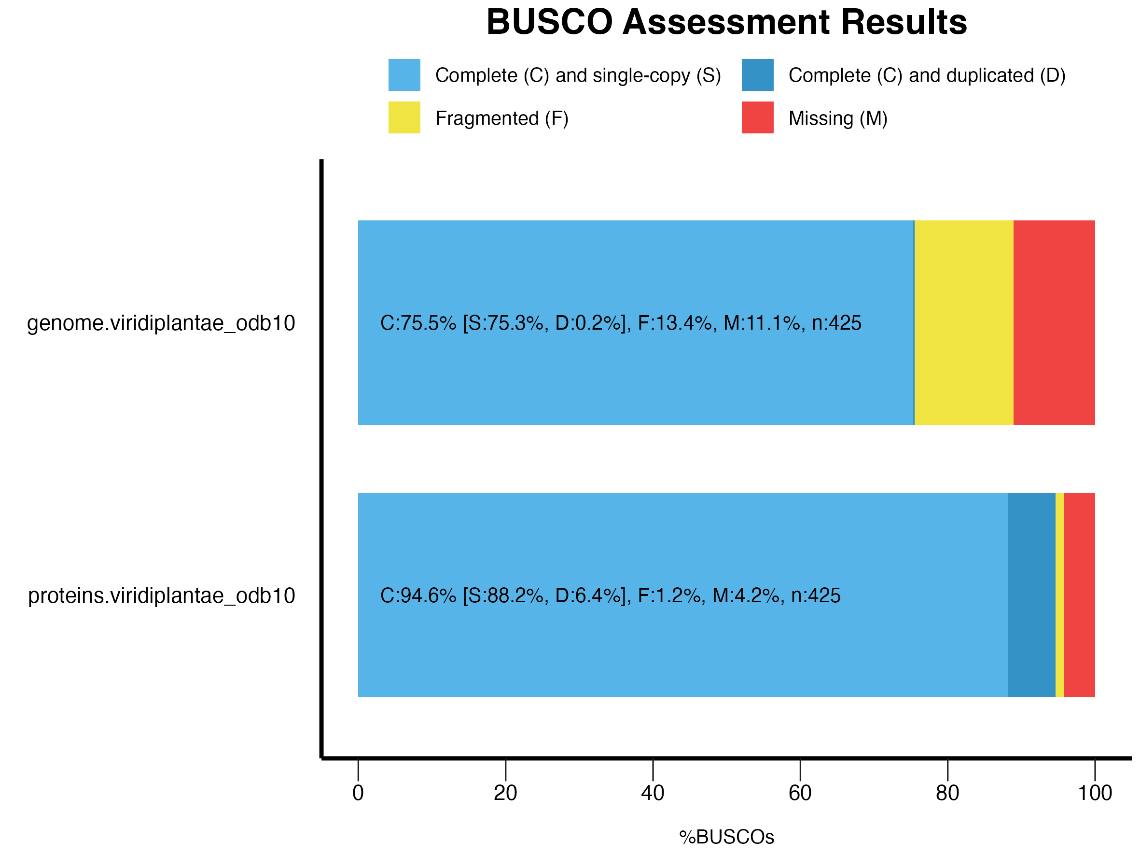

**Supplementary Figure S2: BUSCO evaluation of the *Spirogyra pratensis* annotation.** BUSCO was run using viridiplantae_odb10 as reference with the assembled genome and annotated proteins of *Spirogyra*.

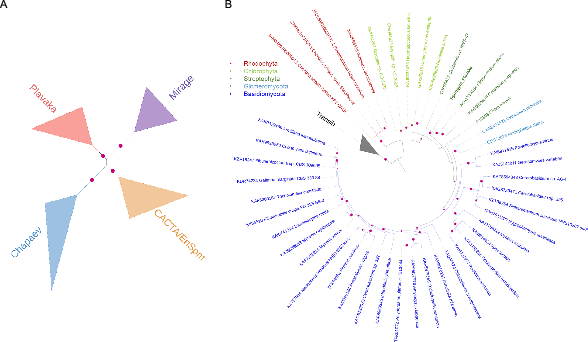

**Supplementary Figure S3:** **Phylogenetic trees of Plavaka transposable elements.** A: CMC clade phylogenetic tree. B: Plavaka transposase tree. The magenta circles indicate bootstrap values above 70%. The Spirogyra sequence is in bold. Tree was rooted on Transib outgroup.

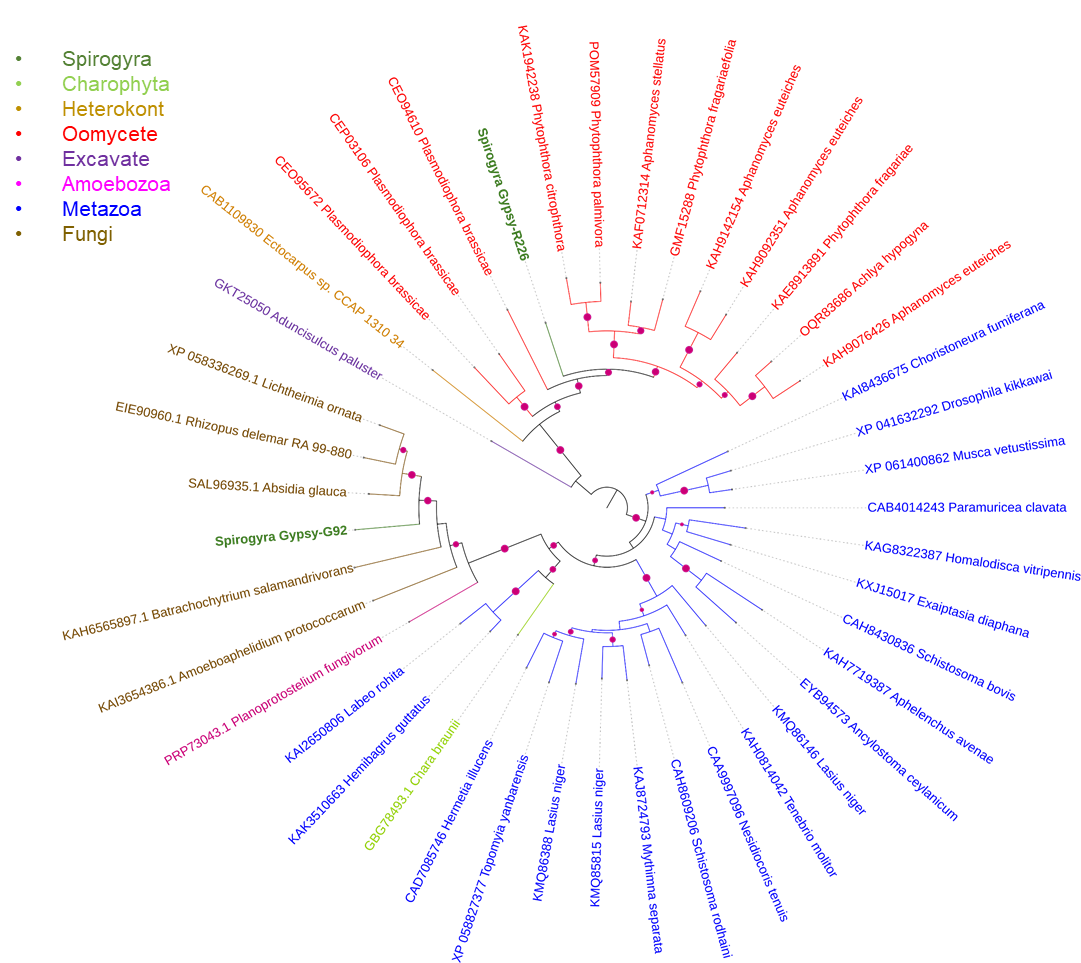

**Supplementary Figure S4: Phylogenetic tree of the Gypsy reverse transcriptase domain showing a putative horizontal gene transfer.** Homologs of *S. pratensis* RT sequences were searched against the GenBank nr proteins database and filtered versions thereof to target or exclude the putative horizontal gene transfer donor (Mucoromycotina and SAR) and recipient clades (Viridiplantae below Tracheophyta). The transposase protein domains were aligned with MAFFT v7.475 and the phylogenetic trees were constructed with IQ-TREE version 1.6.12 (-m MFP+MERGE -bb 1000).

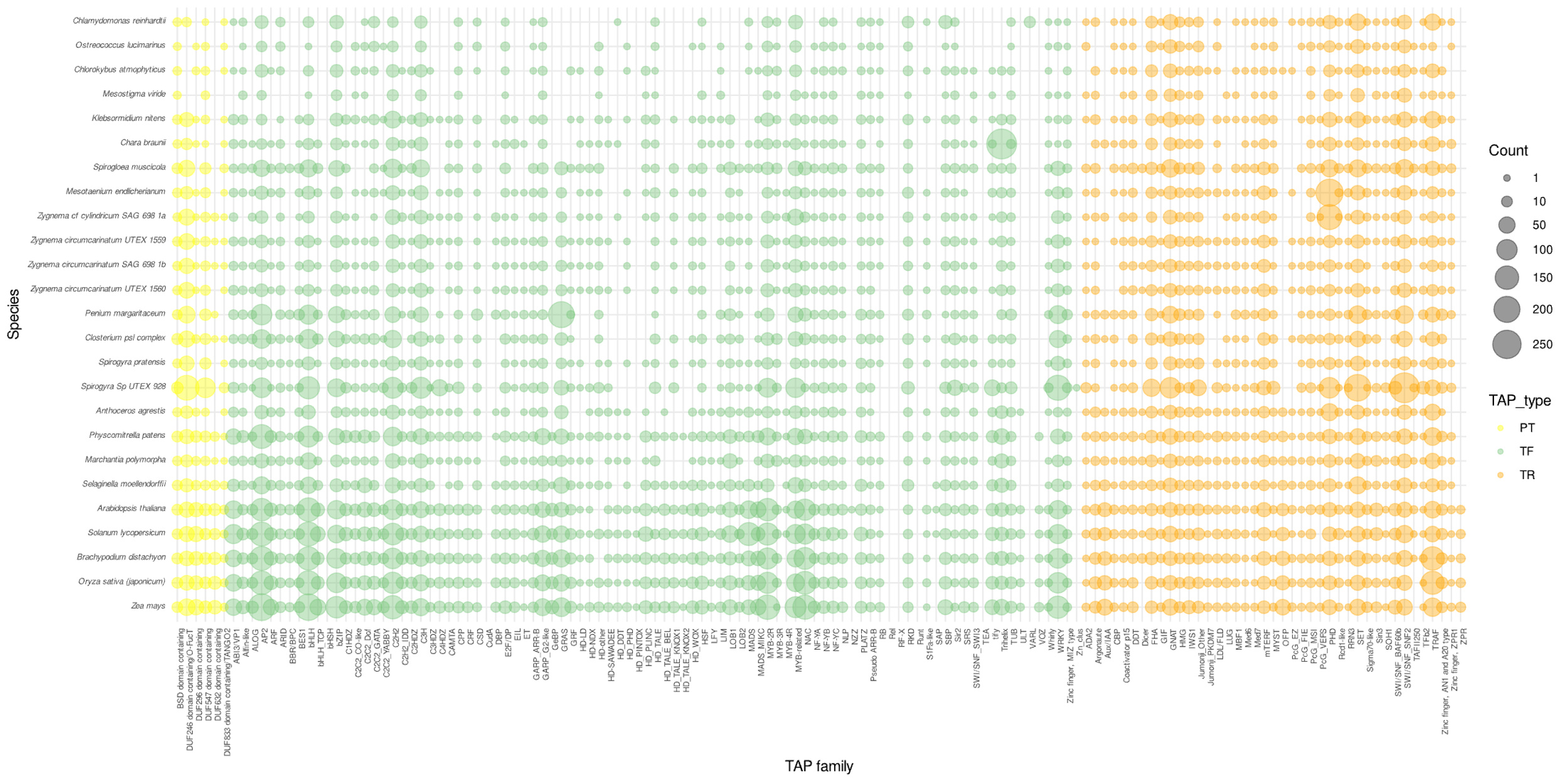
**Supplementary Figure S5: TAP bubble plot.** Number of homologs of transcription associated proteins (TAPs) identified in *Spirogyra* using TAPscan.

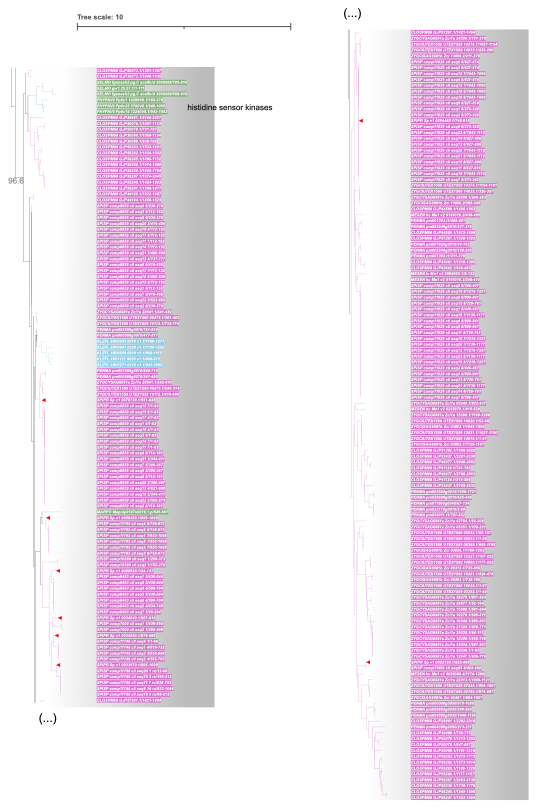

**Supplementary Figure S6: Zoom into Resp_reg motif tree clade containing expansions in Zygnematophyceae**. Zygnematophyceae are coloured in pink, embryophytes in green and other streptopyhte algae in light-blue. Proteins of *Spirogyra pratensis* are marked with a red triangle. Branch support values are non-parametric SH-aRLT. Motif tree was computed using 25 representative viridiplantae species.

**Supplementary Figure S7: Zoom into Resp_reg motif tree showing the clade of type-A RRs**. Zygnematophyceae are coloured in pink, embryophytes in green, other streptopyhte algae in light-blue and chlorophytes in dark-blue. Proteins of *Spirogyra pratensis* are marked with a red triangle. Branch support values are non-parametric SH-aRLT. Motif tree was computed using 25 representative viridiplantae species.

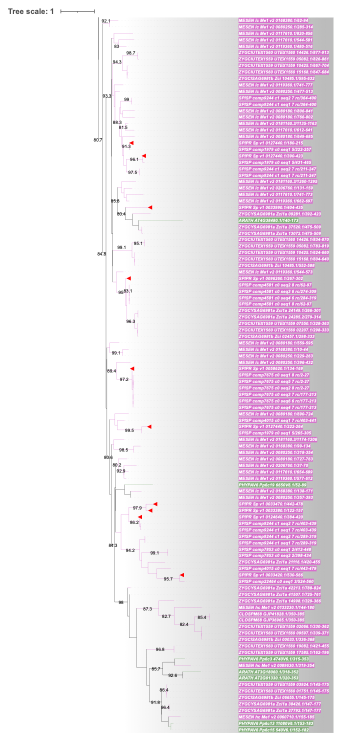

**Supplementary Figure S8: Zoom into WD40 clade containing expansions in Zygnematophyceae**. Zygnematophyceae are coloured in pink and embryophytes in green. Proteins of *Spirogyra pratensis* are marked with a red triangle. Branch support values are non-parametric SH-aRLT. WD40 motif tree was computed with reduced input using only species of Zygnemtophyceae and Embryophyta.

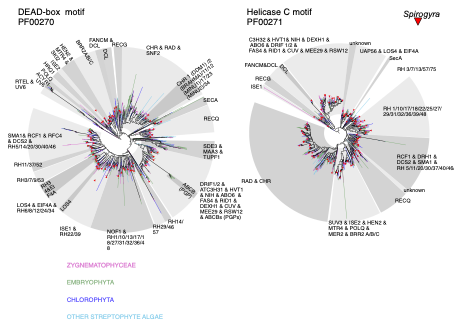

**Supplementary Figure S9: DEAD-box and Helicase C motif trees.** Phylogenetic tree of DEAD-box and Helicase C motifs of different Zygmamatophyceae (in pink), Embryophyta (in green), Chlorophyta (in dark blue) and other streptophyte algae (in light blue). Clades are annotated based on Arabidopsis IDs. *Spirogyra pratensis* sequences are marked with a red triangle. Motif trees were computed using 25 representative viridiplantae species.

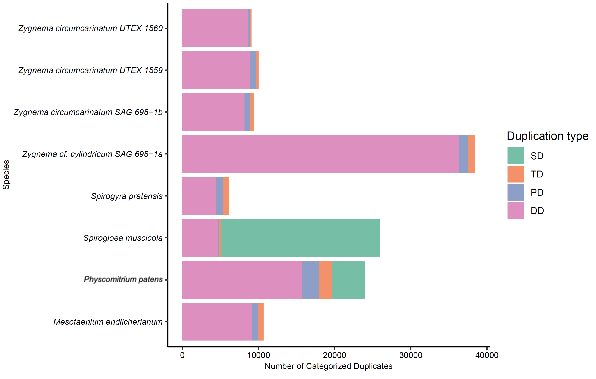

**Supplementary Figure S10: Gene duplication events.** Gene duplication events in *S. pratensis* were assessed using doubletrouble. Genes were classified into four duplication categories: segmental duplication (SD), tandem duplication (TD), proximal duplication (PD), and dispersed duplication (DD).

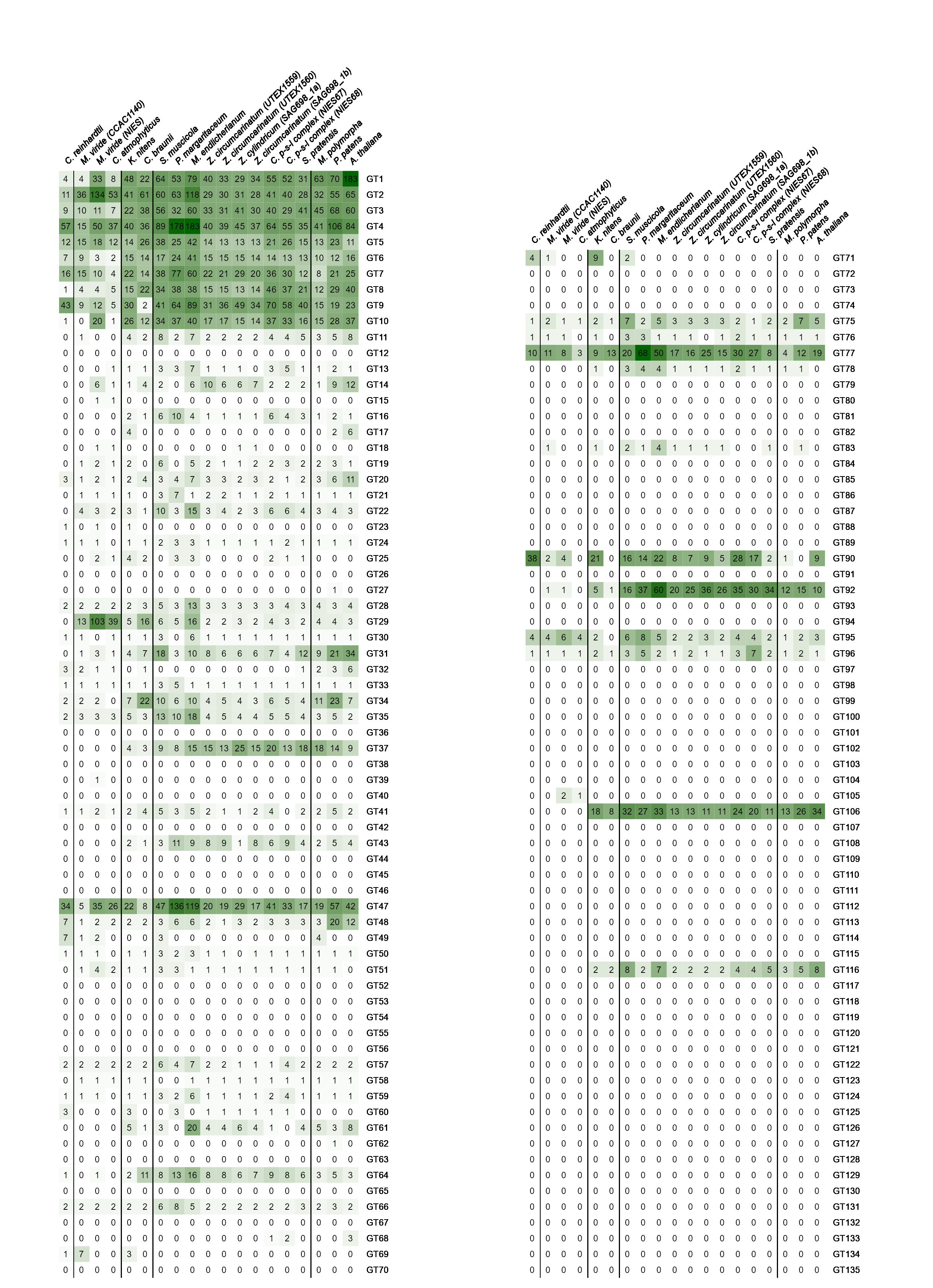

**Supplementary Figure S11: Summary of the glycosyltransferases (GTs) found in *Spirogyra*.** Heatmap showing homologs of *Spirogyra* and other viridiplantae glycosyltransferases. Homologs were identified by computing phylogenetic trees (see Supplementary Figures S38-S58).

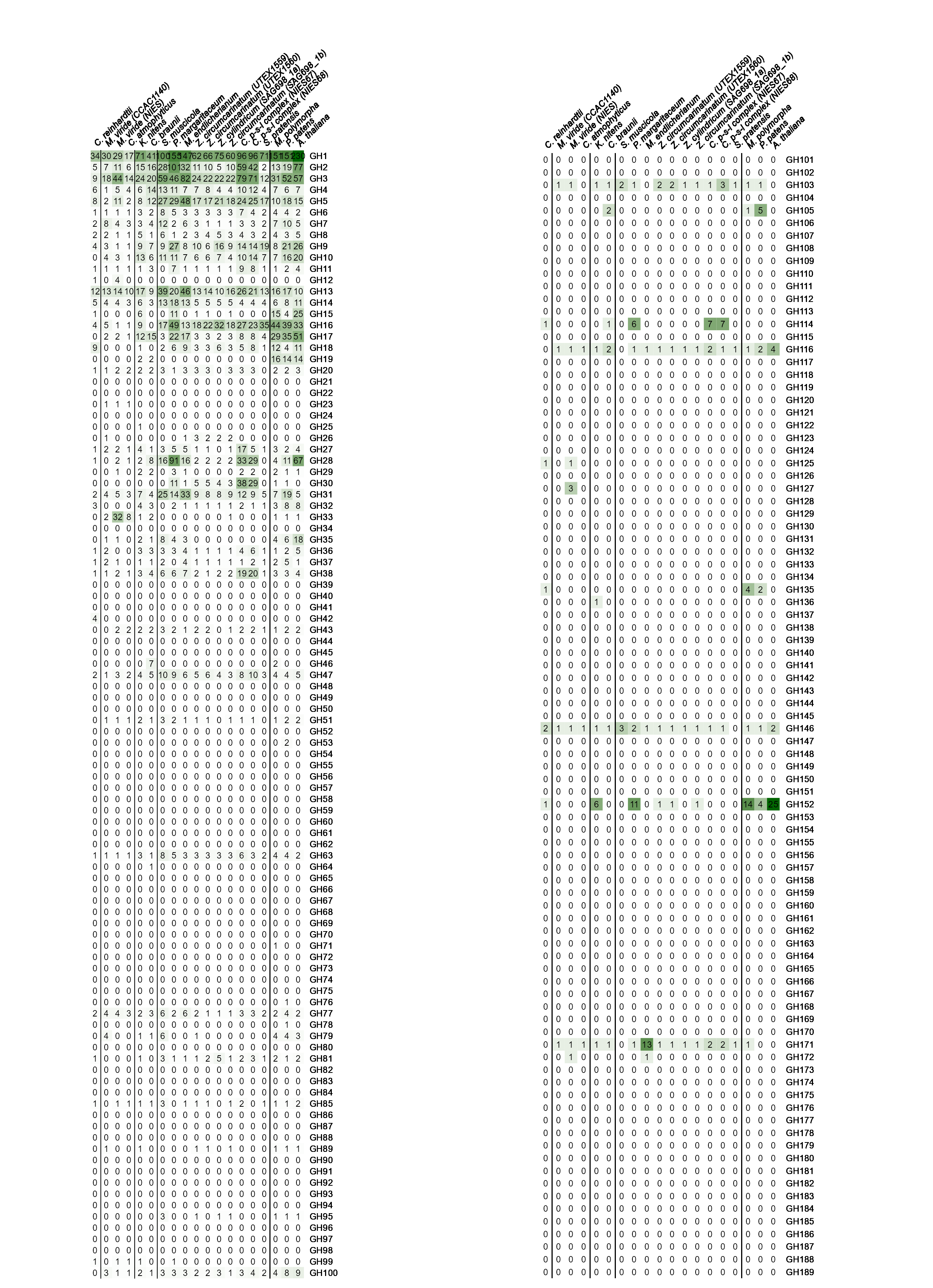
**Supplementary Figure S12: Summary of the glycosylhydrolases (GHs) found in *Spirogyra.*** Heatmap showing homologs of *Spirogyra* and other viridiplantae glycosylhydrolases. Homologs were identified by computing phylogenetic trees (see Supplementary Figures S38-S58).

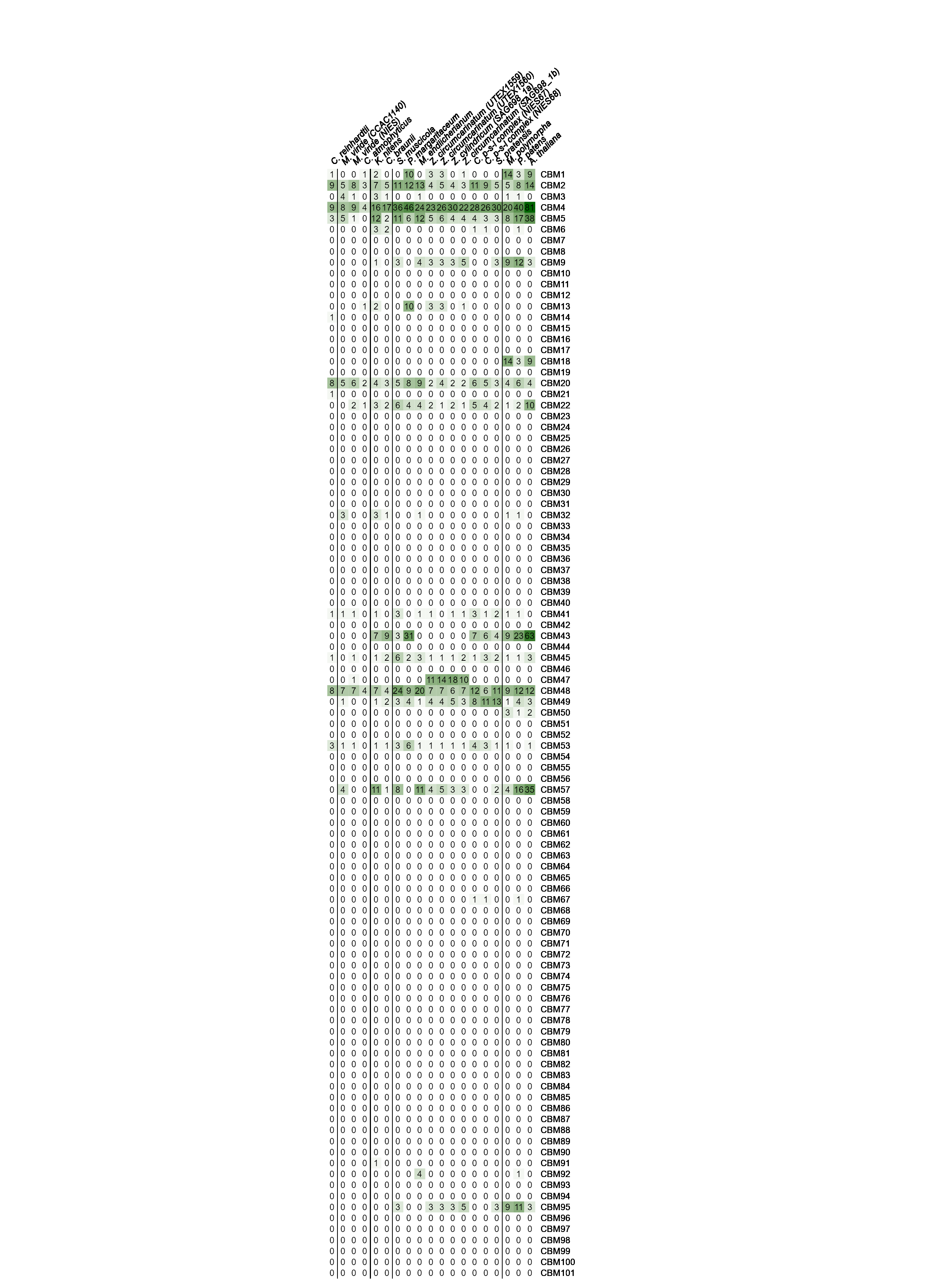
**Supplementary Figure S13: Summary of the carbohydrate-binding modules (CBMs).** Heatmap showing homologs of *Spirogyra* and other viridiplantae carbohydrate-binding modules. Homologs were identified by computing phylogenetic trees (see Supplementary Figures S38-S58).

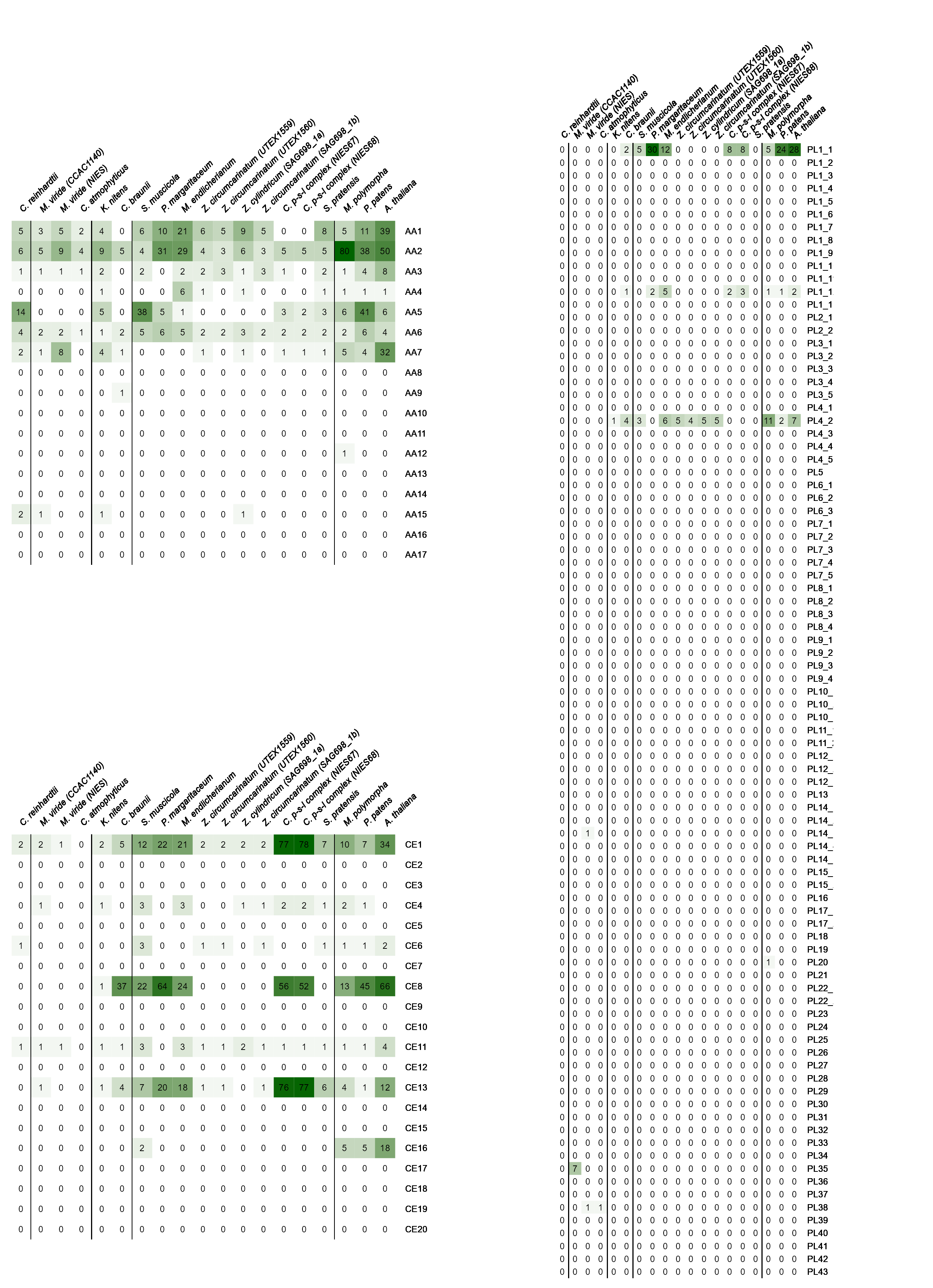

**Supplementary Figure S14: Summary of the enzymes with auxiliary activity (AAs), pectate lyases (PLs) and carbohydrate esterases (CEs) found in *Spirogyra.*** Heatmap showing homologs of *Spirogyra* and other viridiplantae enzymes with auxiliary activity, pectin lyases and carbohydrate esterases. Homologs were identified by computing phylogenetic trees (see Supplementary Figures S38-S58).

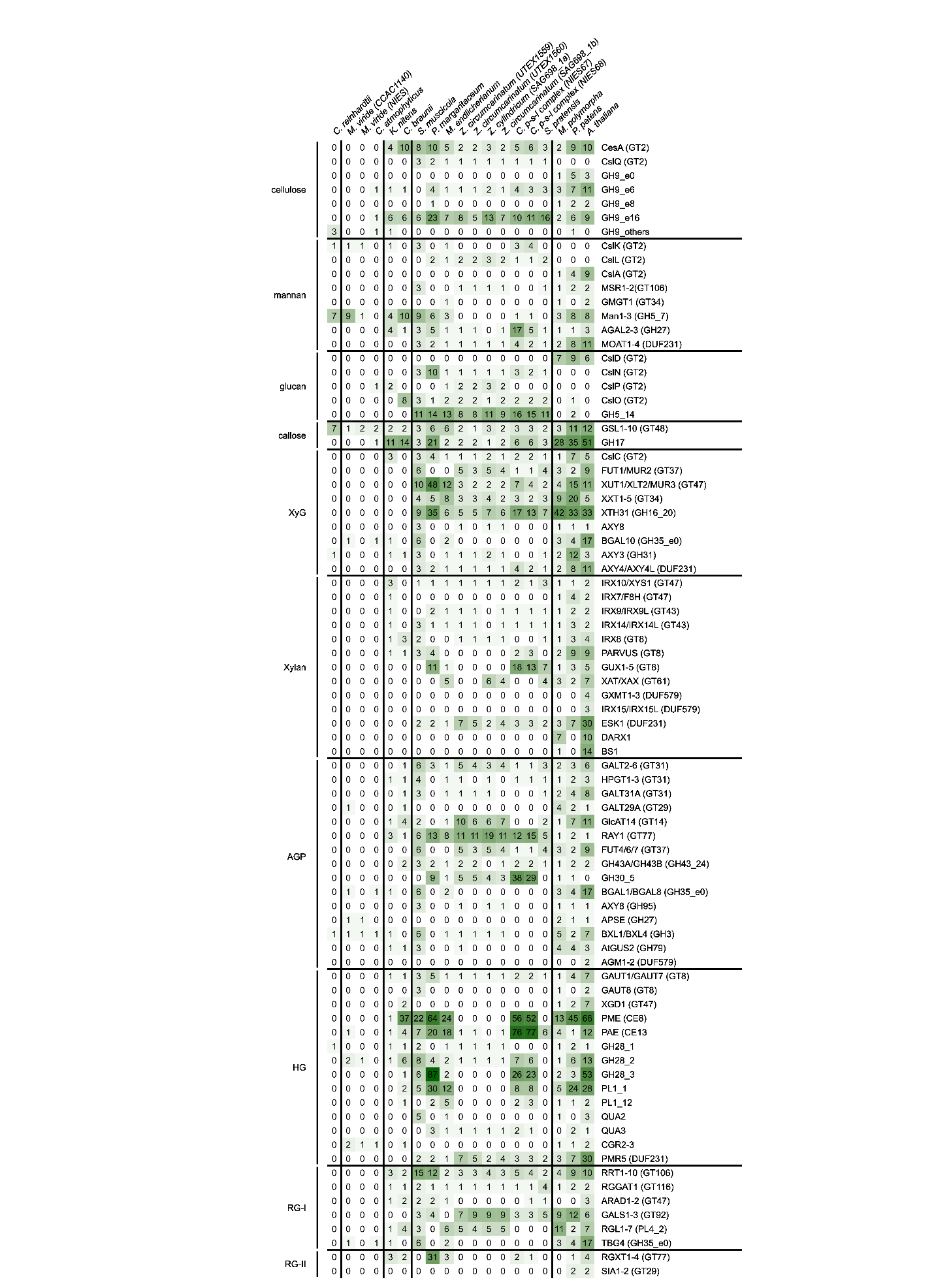
**Supplementary Figure S15: Summary of CAZymes found in *Spirogyra.*** Heatmap showing homologs of *Spirogyra* and other viridiplantae carbohydrate-active enzymes (CAZymes). Homologs were identified by computing phylogenetic trees (see Supplementary Figures S38-S58).

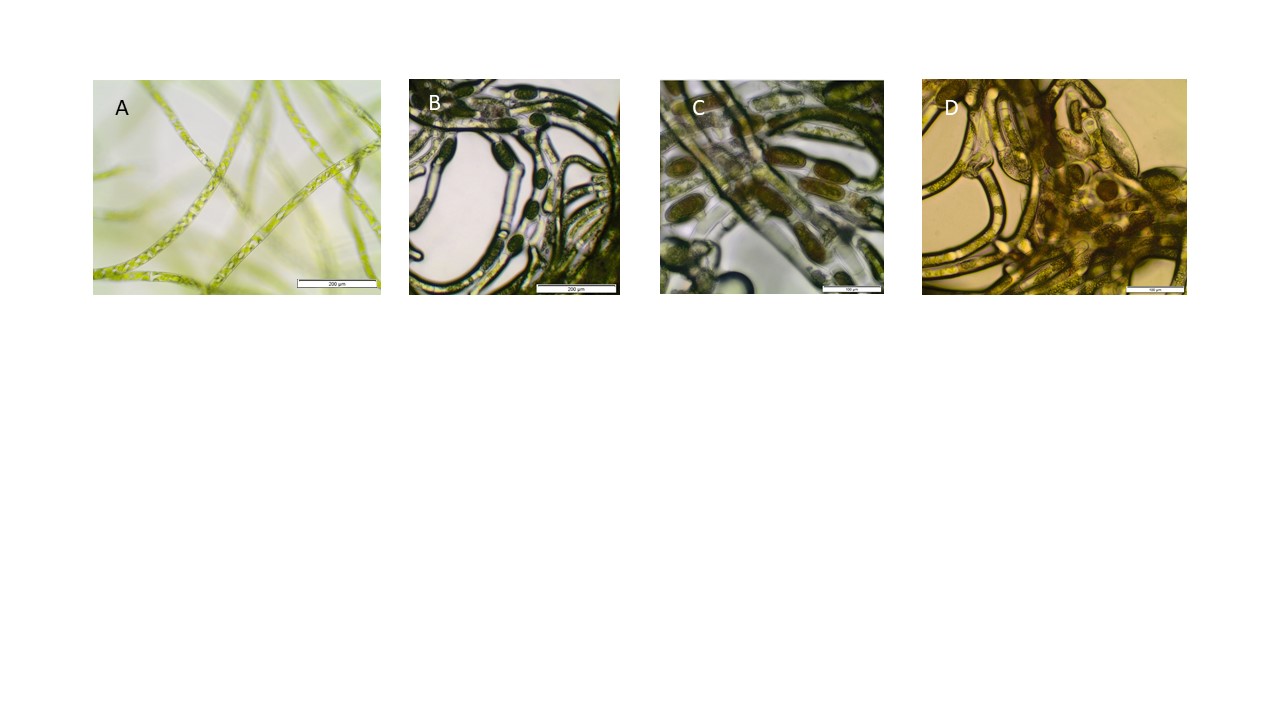

**Supplementary Figure S16: Light microscopic images of S. *pratensis* MZCH#10213 life cycle stages**. (A) vegetative cells, after 3 days on solid medium, (B) young spores 10 days after 10 days on solid C-medium, spores are green and most of remaining cells viable; (C) immature spores after 14 days, spores turn brown but still contain green residues, (D) mature spore after 31 days, mostly colored brown, storage vesicles visible. Scale bar in (A) and (B) 200 µm, in (C) and (D) 100 µm).

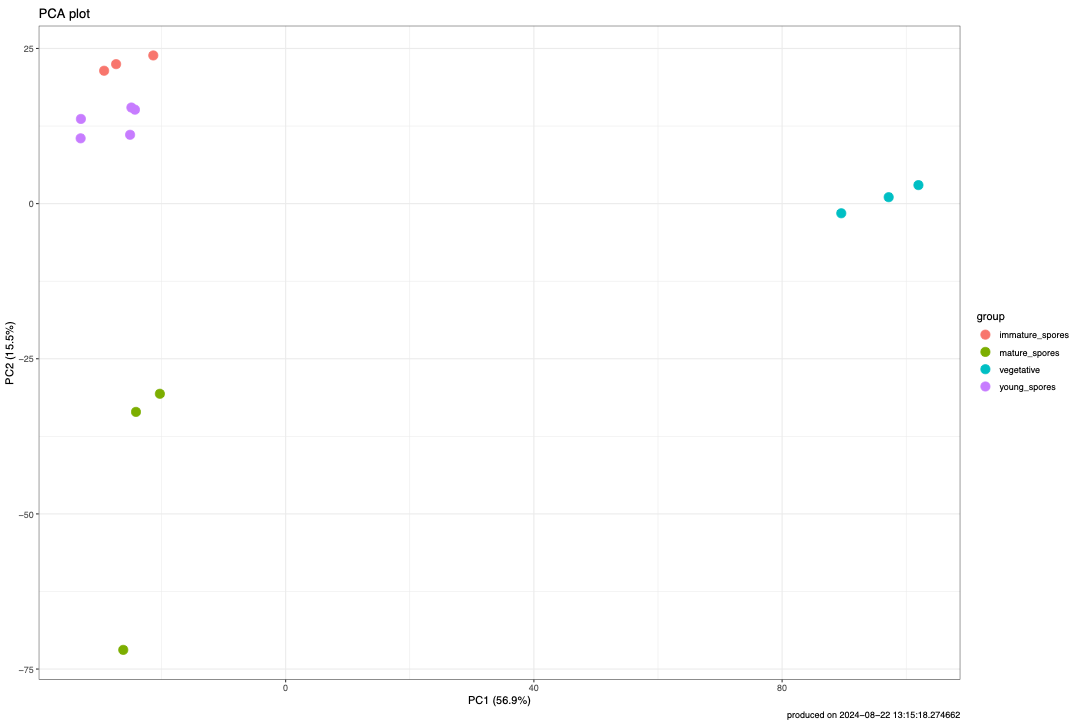

**Supplementary Figure S17: Principal component analysis of RNA-seq data of developmental stages.** PCA of the qsmooth normalized log2 count per million (cpm) values of the RNA-seq data generated from different developmental stages of *Spirogyra.*

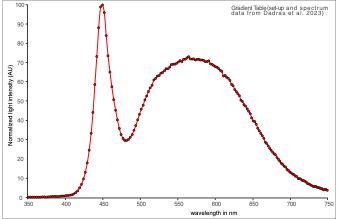

**Supplementary Figure S18: Light Spectrum of the gradient table set-up.** Average light spectrum of gradient table set-up. Set-up and spectrum data is the same as in Dadras et al. 2023).

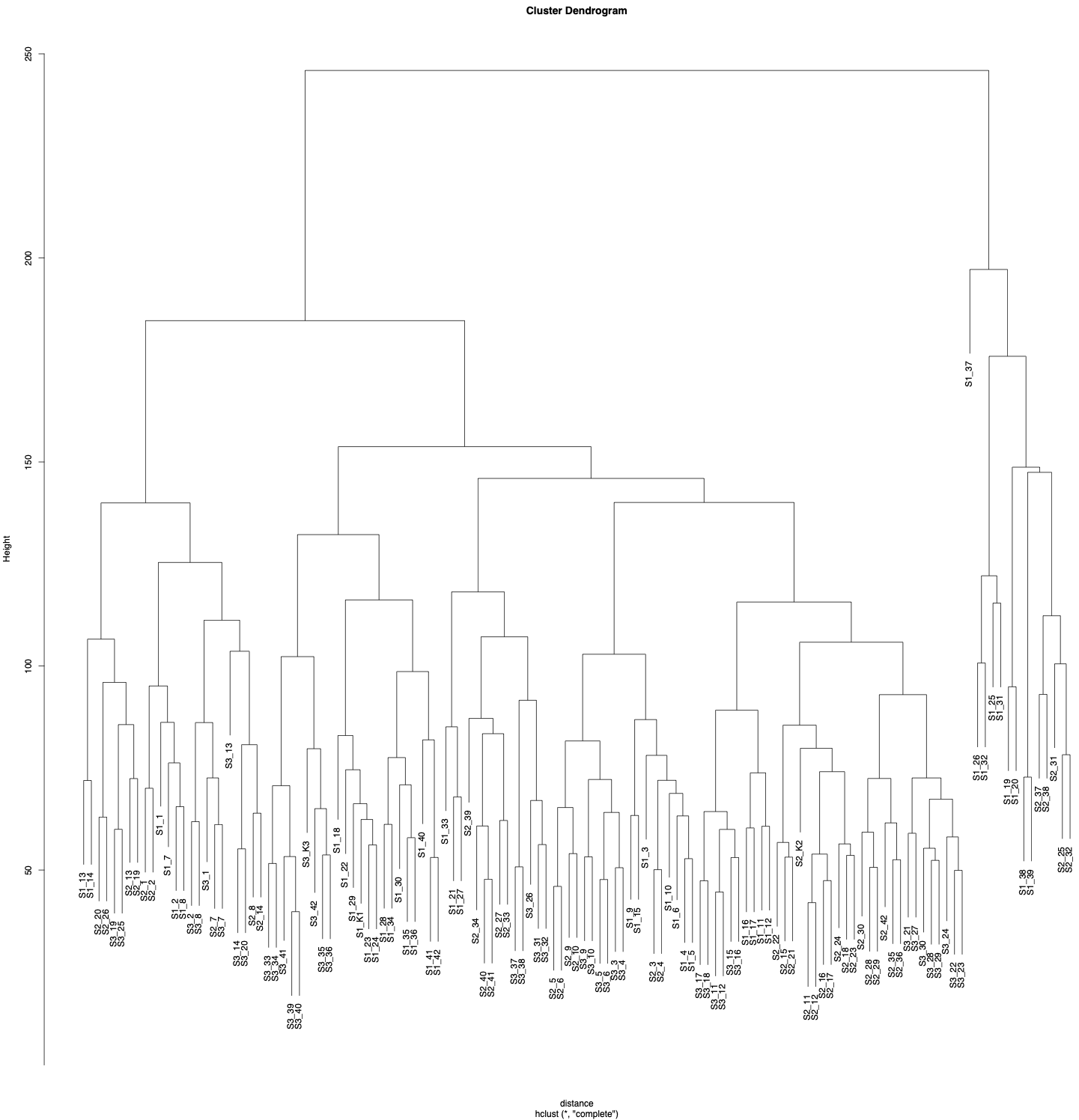

**Supplementary Figure S19: Hierarchical clustering of gradient table RNA-seq samples.** Sample S1_37 (29°C; 530 µmol photons s^–1^ m^–2^) was identified as an outlier and removed from further analysis.

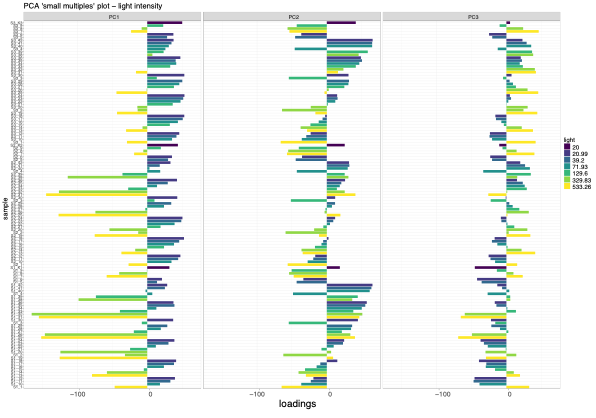

**Supplementary Figure S20: Small multiple PCA, impact of light intensity on RNA-seq data, gradient-table experiment.** Influence of the experimental variable light intensity across all RNA-seq samples of gradient table set-up on the principal components 1 to 3. Principal component 1 is strongly influenced by light intensity.

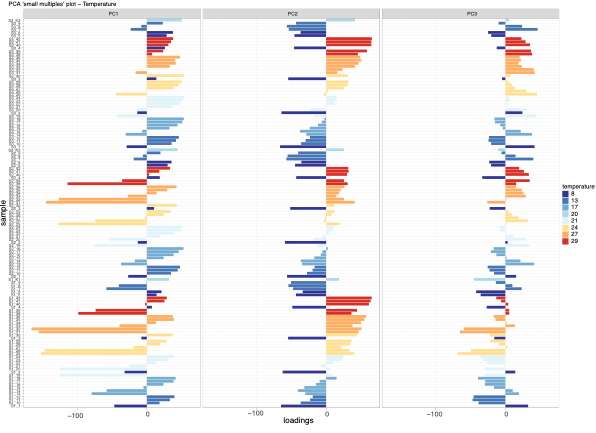

**Supplementary Figure S21: Small multiple PCA, impact of temperature on RNA-seq data, gradient-table experiment.** Influence of the experimental variable temperature across all RNA-seq samples of gradient table set-up on the principal components 1 to 3. Principal component 2 is strongly influenced by temperature.

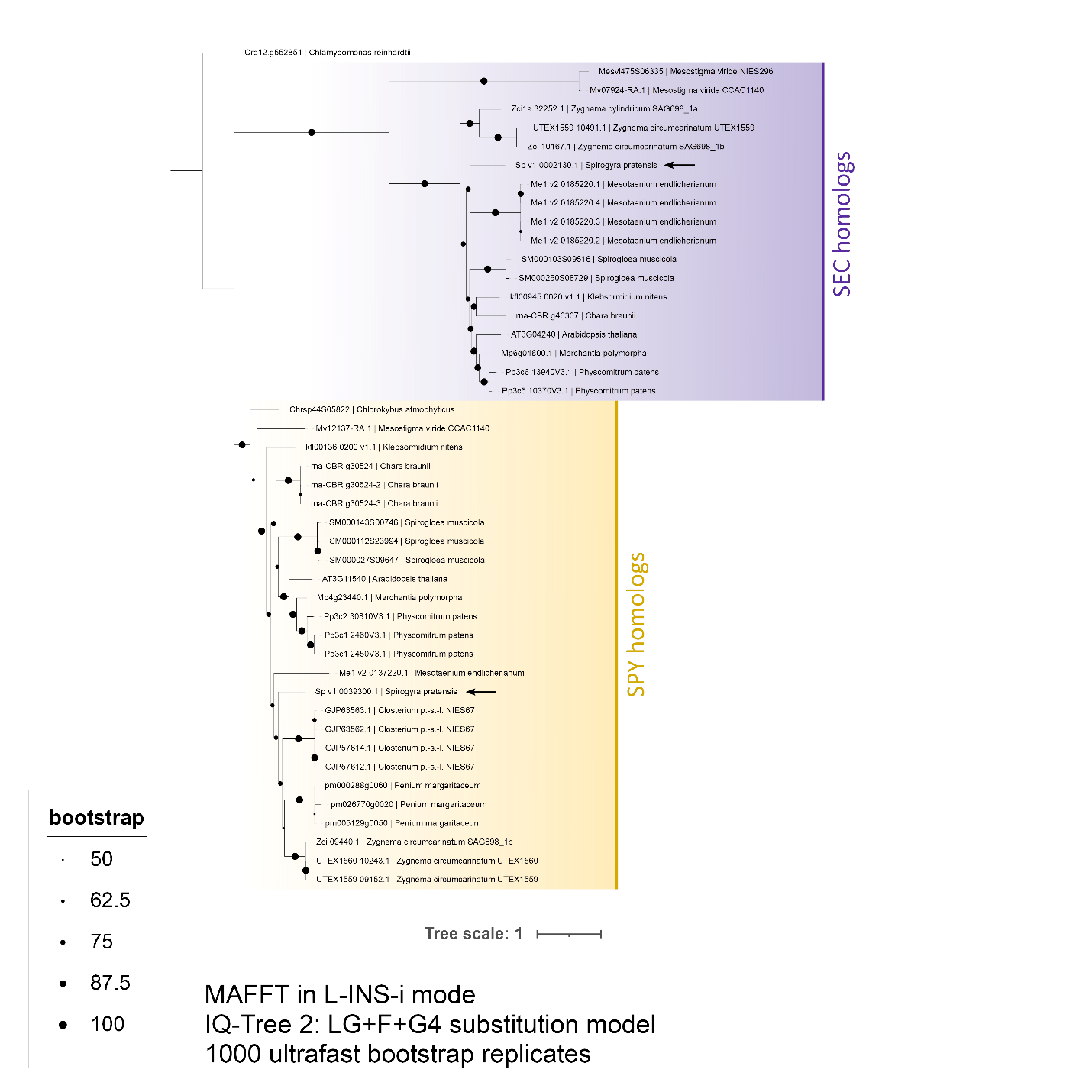

**Supplementary Figure S22: Phylogeny of glycosyltransferases from family 41.** *Spirogyra* homologs are marked with an arrow.

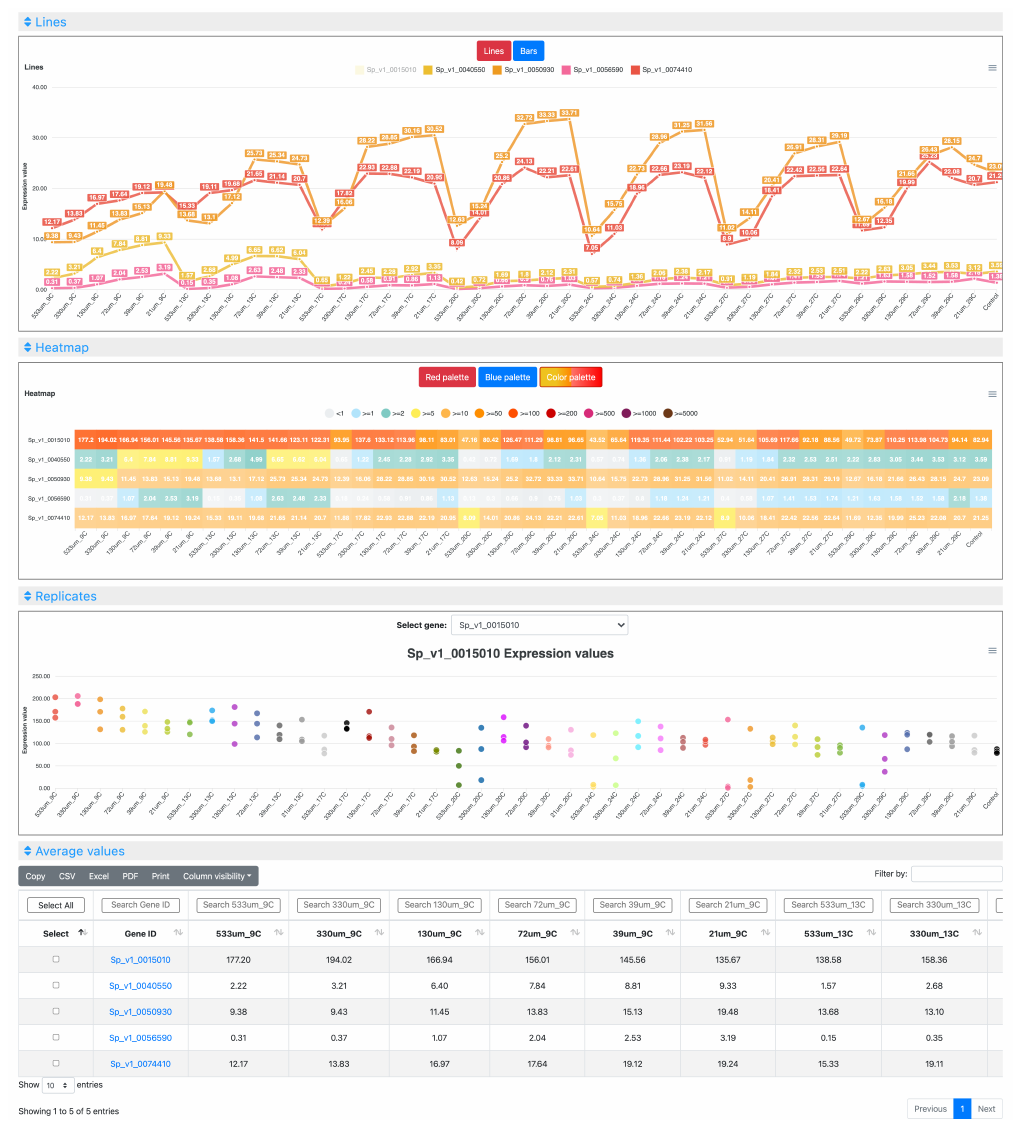

**Supplementary Figure S23: *S. pratensis* gene expression atlas at MAdLandExpression, showing expression of five genes along the temperature and light intensity gradient experiment**. From top to bottom: “Lines" and “Heatmap" plots alllow the comparison of multiple genes simultaneously; the “Replicates" plot show the behavior of each replicate of the selected gene across all experimental conditions; and the "Average values" table provides expression data visualization and downloading, along with functional annotations.

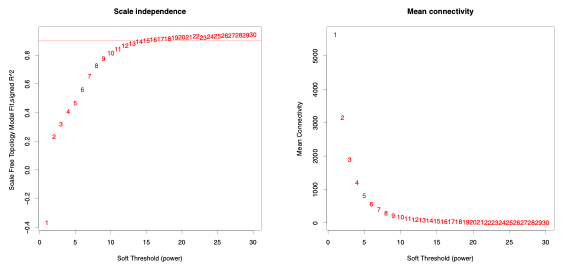

**Supplementary Figure S24: Picking a soft threshold power based on scale free topology and mean connectivity.** Thresholding powers of 1 to 20 were plotted against scale free topology and mean connectivity. A soft threshold power of 14 was chosen to construct the WGNCA.

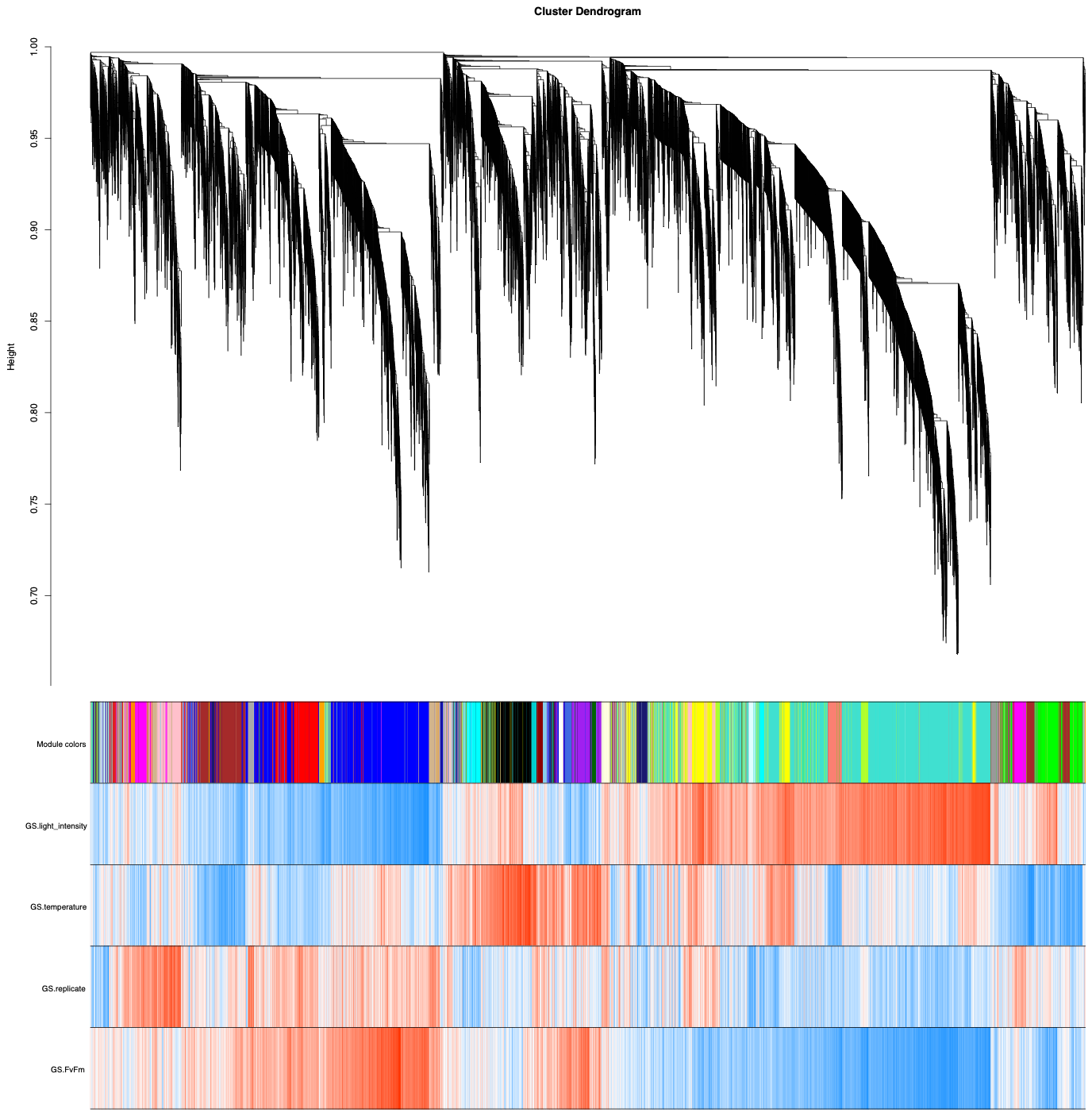

**Supplementary Figure S25: Gene Cluster Dengrogram and Module Membership.** Dendrogram showing hierarchical clustered genes used for WGCNA construction. The heatmap below the dendrogram shows the gene significance of genes in relation to the experimental parameters light intensity, temperature, biological replicate and Fv/Fm. Positive correlations are shown in red, negative correlations are shown in blue.

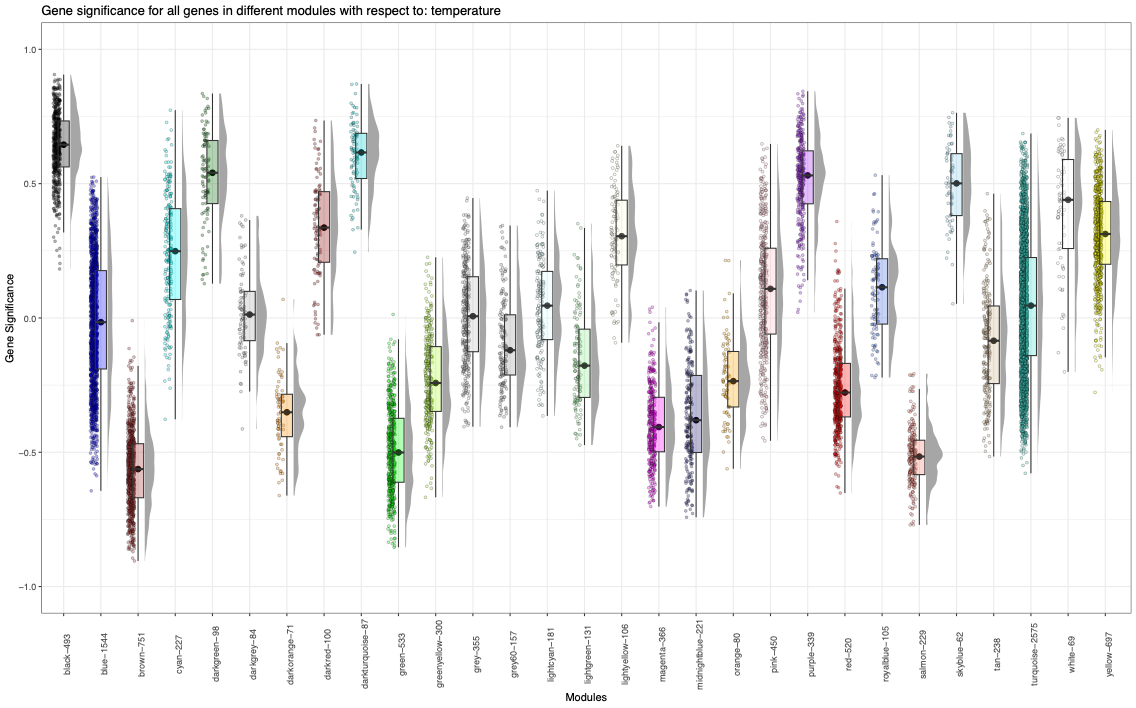

**Supplementary Figure S26: Gene significance for all genes split into WGNCA modules in relation to temperature.** Box plots show the mean gene significance and interquartile range of the gene significance for all WGCNA modules in respect to the experimental condition temperature. Each data point symbolizes a gene.

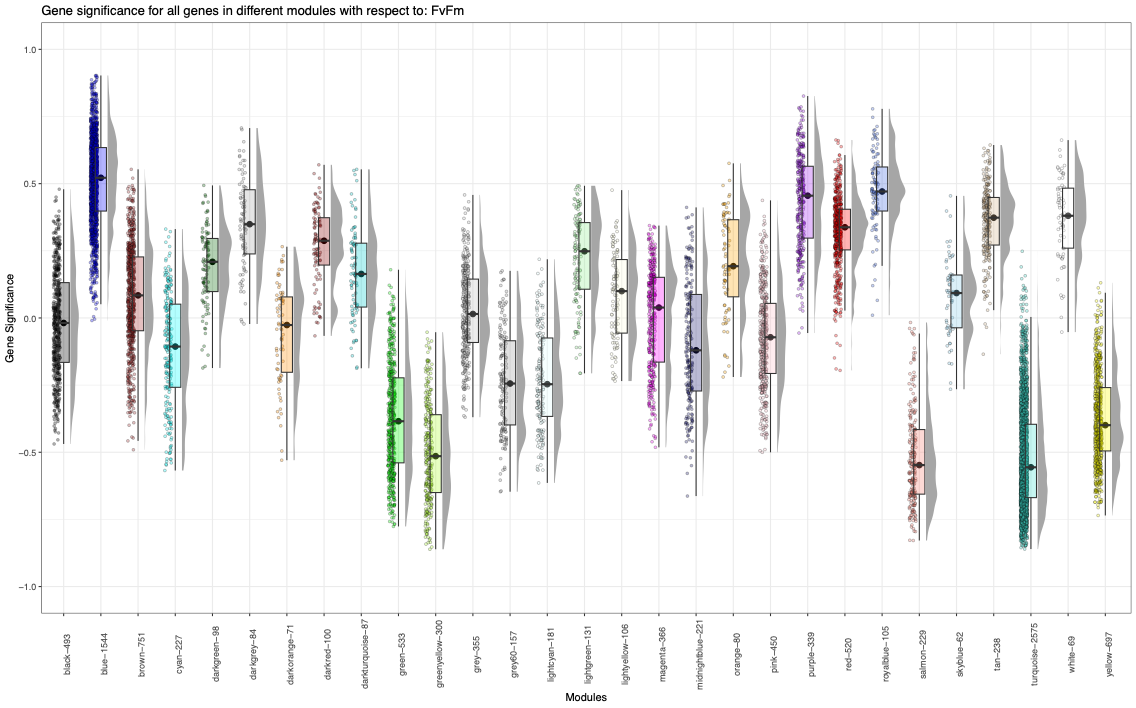

**Supplementary Figure S27: Gene significance for all genes split into WGNCA modules in relation to Fv/Fm.** Box plots show the mean gene significance and interquartile range of the gene significance for all WGCNA modules in respect to the experimental condition Fv/Fm. Each data point symbolizes a gene.

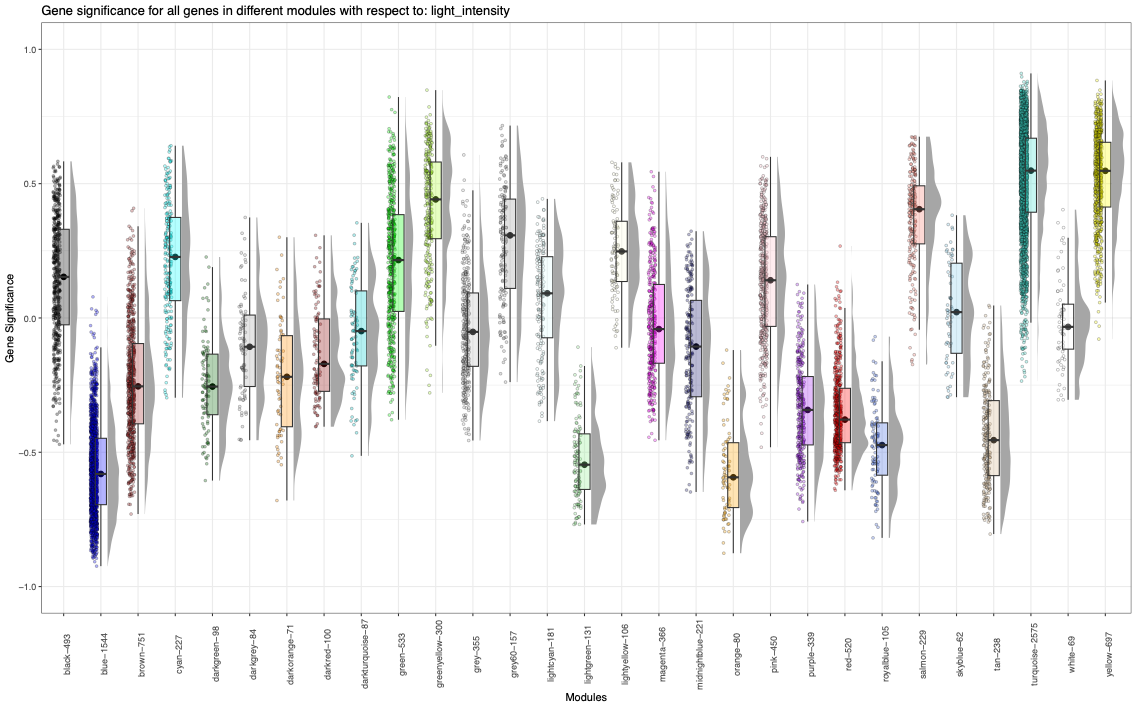

**Supplementary Figure S28: Gene significance for all genes split into WGNCA modules in relation to light intensity.** Box plots show the mean gene significance and interquartile range of the gene significance for all WGCNA modules in respect to the experimental condition light intensity. Each data point symbolizes a gene.

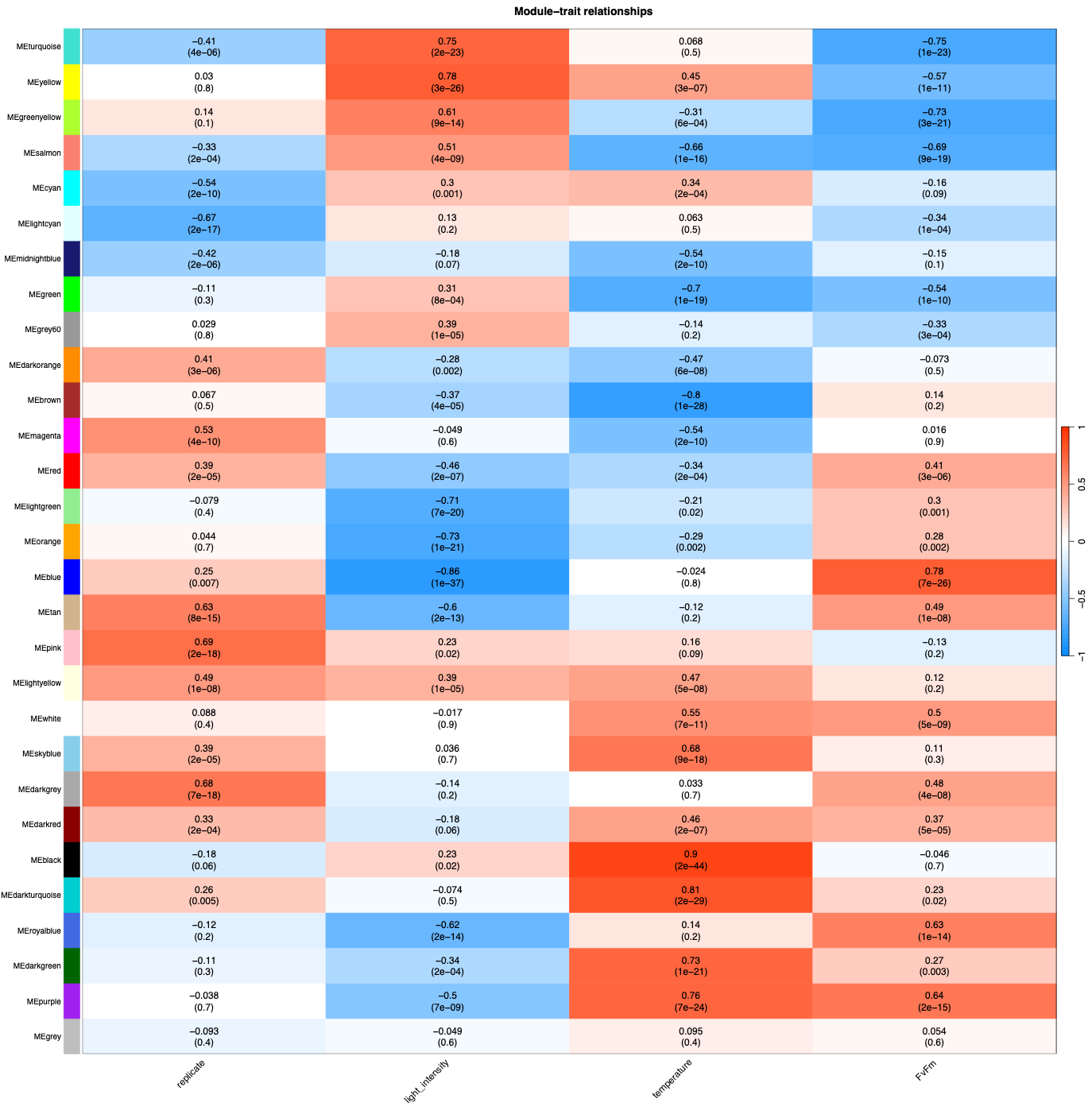

**Supplementary Figure S29:** **Module-trait relationship of WGCNA.** Eigengenes of the WGCNA modules were correlated to the variables light intensity, temperature, Fv/Fm and biological replicate. The heatmap shows the correlation as well as the p-value of a Student’s t-test.

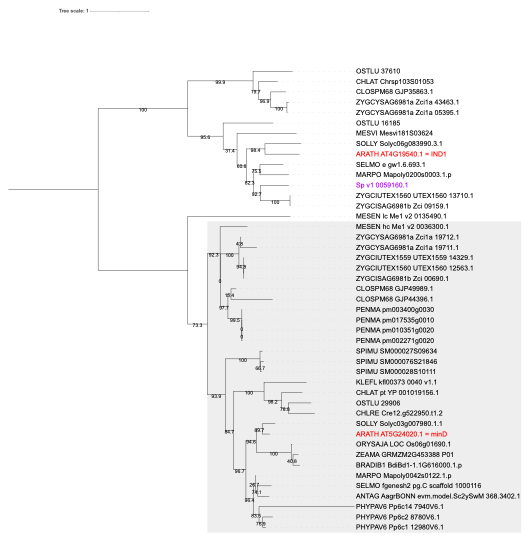

**Supplementary Figure S30: MinD phylogenetic tree.** Homologs of *Arabidopsis* minD were identified using blastP, aligned using mafft and phylogenetic trees were calculated using iqtree2. *Arabidopsis* proteins are marked in red, *Spirogyra* proteins in purple. Branch support values are non-parametric SH-aRLT. Tree is midpoint rooted. Supplement to Figure 5e (bubble plot).

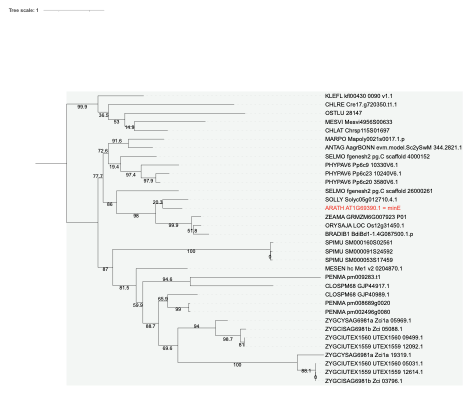

**Supplementary Figure S31: MinE Phylogenetic tree.** Homologs of *Arabidopsis* minE were identified using blastP, aligned using mafft and phylogenetic trees were calculated using iqtree2. *Arabidopsis* proteins are marked in red, *Spirogyra* proteins in purple. Branch support values are non-parametric SH-aRLT. Tree is midpoint rooted. Supplement to Figure 5e (bubble plot).

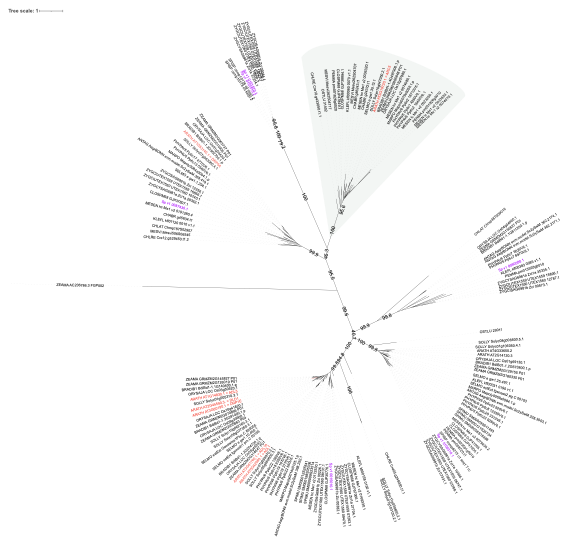

**Supplementary Figure S32: ARC5 phylogenetic tree.** Homologs of *Arabidopsis* ARC5 were identified using blastP, aligned using mafft and phylogenetic trees were calculated using iqtree2. *Arabidopsis* proteins are marked in red, *Spirogyra* proteins in purple. Branch support values are non-parametric SH-aRLT. Tree is midpoint rooted. Supplement to Figure 5e (bubble plot).

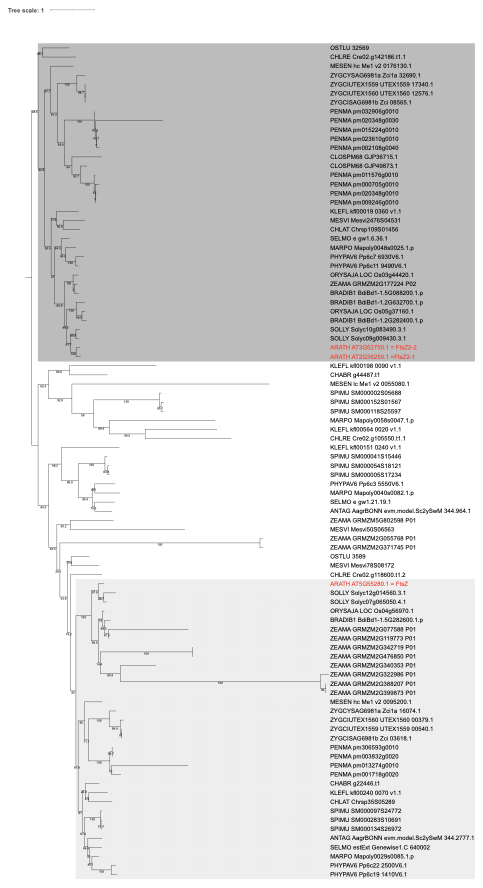

**Supplementary Figure S33: FtsZ phylogenetic tree.** Homologs of *Arabidopsis* FtsZ were identified using blastP, aligned using mafft and phylogenetic trees were calculated using iqtree2. *Arabidopsis* proteins are marked in red, *Spirogyra* proteins in purple. Branch support values are non-parametric SH-aRLT. Tree is midpoint rooted. Supplement to Figure 5e (bubble plot).

**Supplementary Figure S34: PDV phylogenetic tree.** Homologs of *Arabidopsis* PDV1 and PDV2 were identified using blastP, aligned using mafft and phylogenetic trees were calculated using iqtree2. *Arabidopsis* proteins are marked in red, *Spirogyra* proteins in purple. Branch support values are non-parametric SH-aRLT. Tree is midpoint rooted. Supplement to Figure 5e (bubble plot).

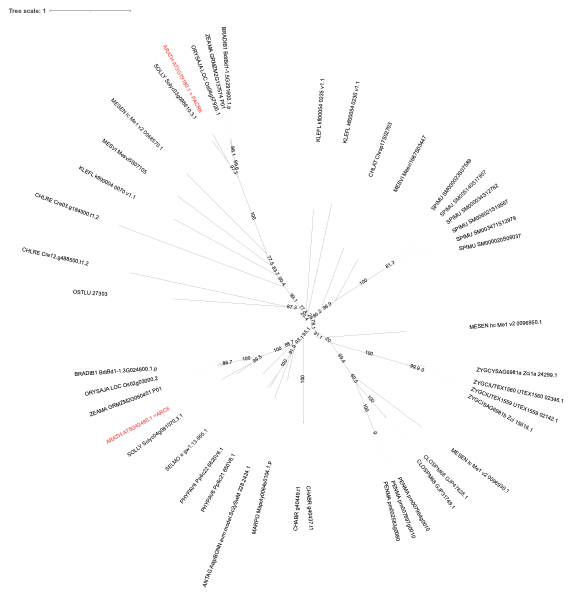

**Supplementary Figure S35: ARC6 and PARC6 phylogenetic tree.** Homologs of *Arabidopsis* ARC6 and PARC6 were identified using blastP, aligned using mafft and phylogenetic trees were calculated using iqtree2. *Arabidopsis* proteins are marked in red, *Spirogyra* proteins in purple. Branch support values are non-parametric SH-aRLT. Supplement to Figure 5e (bubble plot).

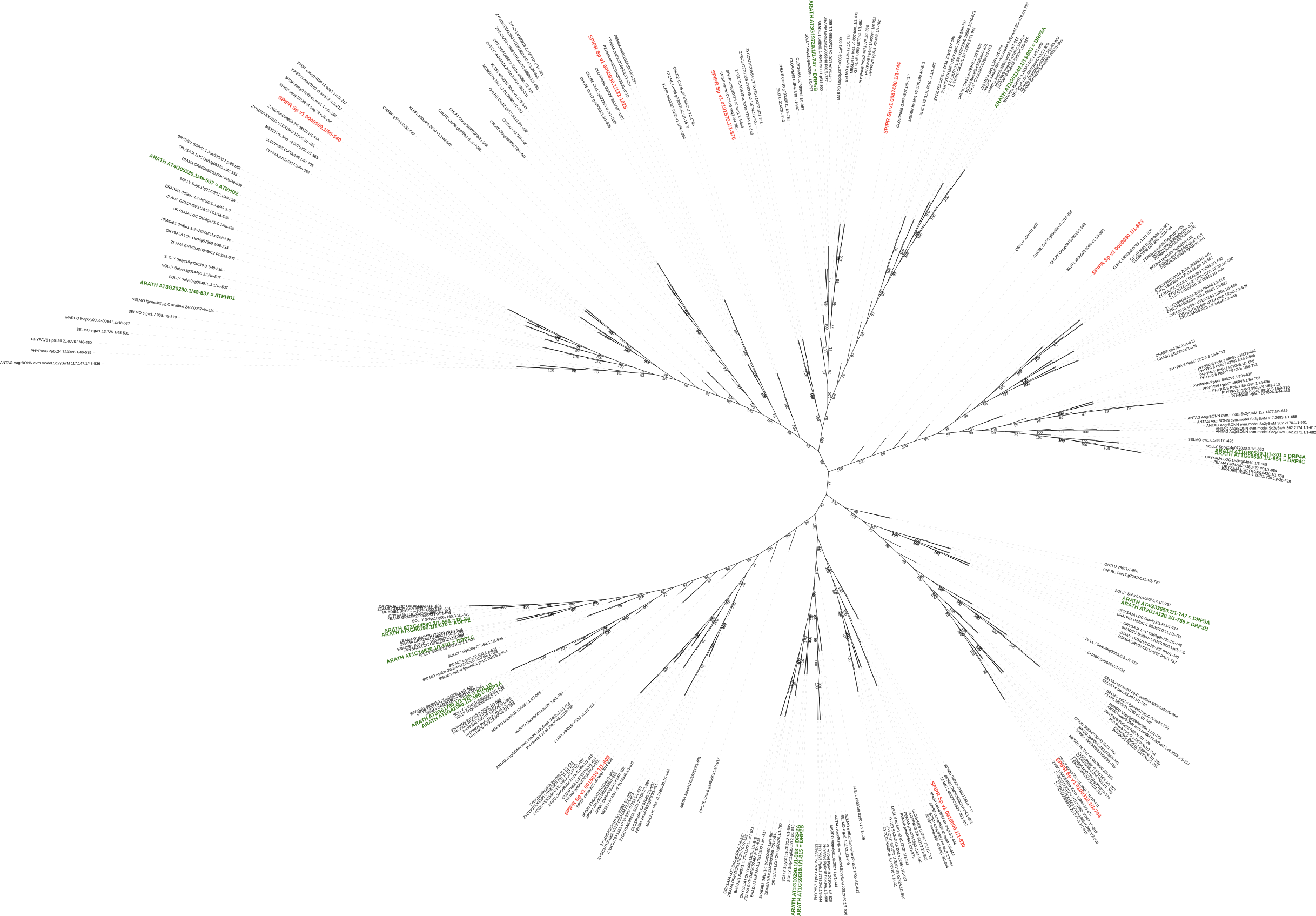
**Supplementary Figure S36: Dynamin tree (unrooted).** *S. pratensis* and *A. thaliana* gene Ids are highlighted in red and green, respectively.

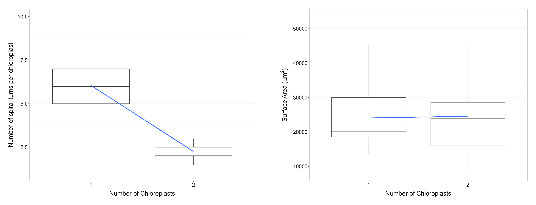
**Supplementary Figure S37:** **Quantification of spiral turns and surface area of *Spirogyra* chloroplasts.** Left panel: Number of chloroplasts negatively correlates with the number of spiral turns per chloroplast (R^2^ = 0.69, *p < 0.001*, regression equation y= -3.80x + 9.84). Right panel: No significant difference between the chloroplast surface area in cells with one or two chloroplasts (p=0.84).

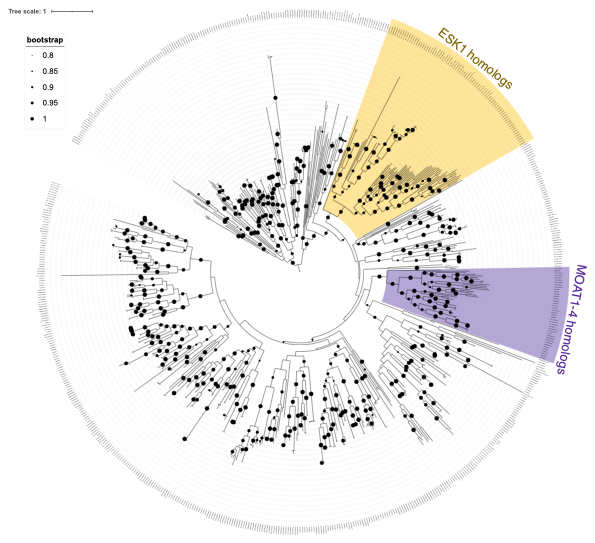
**Supplementary Figure S38: CAZyme (DUF231) ESK1 and MOAT1-4 tree.** *Spirogyra* sequences were compared with a set of Viridiplantae species. For that, protein sequences of those families were aligned using MAFFT (in L-INS-i mode). IQTREE2 (-st AA -m TEST -bb 1000 -alrt 1000) or FASTTREE (LG+CAT, version 2.1.10) were used for inference of maximum-likelihood trees. Supplementary Data for Supplementary Figure S11-S15.

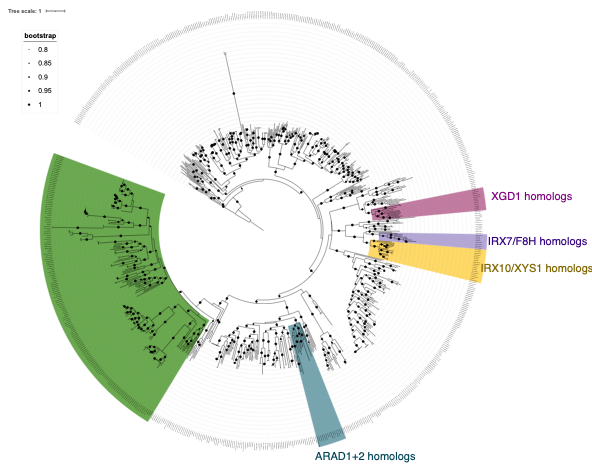

**Supplementary Figure S39: CAZyme (GT47) XGD1, IRX7/F8H, IRX10/XYS1 and ARAD1+2 tree.** *Spirogyra* sequences were compared with a set of Viridiplantae species. For that, protein sequences of those families were aligned using MAFFT (in L-INS-i mode). IQTREE2 (-st AA -m TEST -bb 1000 -alrt 1000) or FASTTREE (LG+CAT, version 2.1.10) were used for inference of maximum-likelihood trees. Supplementary Data for Supplementary Figure S11-S15.

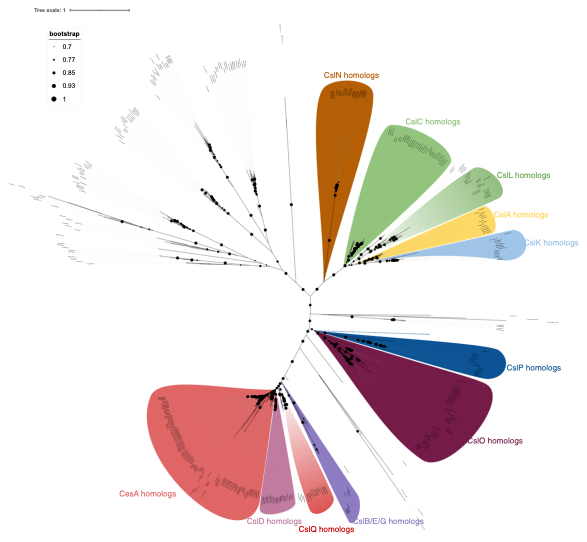

**Supplementary Figure S40: CAZyme (GT2) CesA, CsID, CsIQ, CsIB/E/G, CslO, CsiP, CsIK, CsIA, CsIL, CsIC and CSiN tree.** *Spirogyra* sequences were compared with a set of Viridiplantae species. For that, protein sequences of those families were aligned using MAFFT (in L-INS-i mode). IQTREE2 (-st AA -m TEST -bb 1000 -alrt 1000) or FASTTREE (LG+CAT, version 2.1.10) were used for inference of maximum-likelihood trees. Supplementary Data for Supplementary Figure S11-S15.

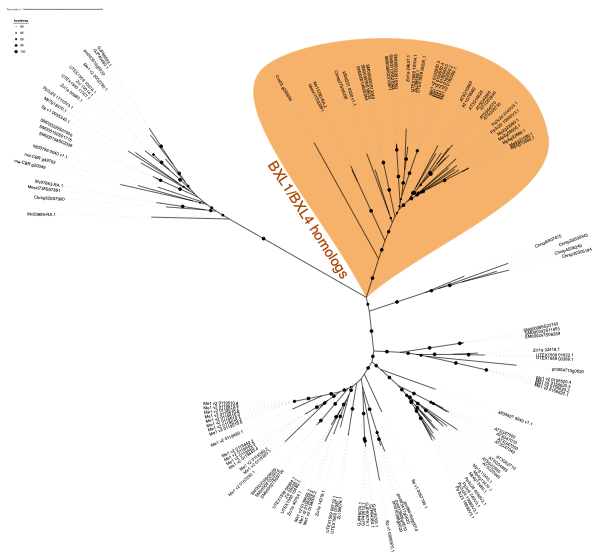

**Supplementary Figure S41: CAZyme (GH3) BXL1/BXL4 tree.** *Spirogyra* sequences were compared with a set of Viridiplantae species. For that, protein sequences of those families were aligned using MAFFT (in L-INS-i mode). IQTREE2 (-st AA -m TEST -bb 1000 -alrt 1000) or FASTTREE (LG+CAT, version 2.1.10) were used for inference of maximum-likelihood trees. Supplementary Data for Supplementary Figure S11-S15.

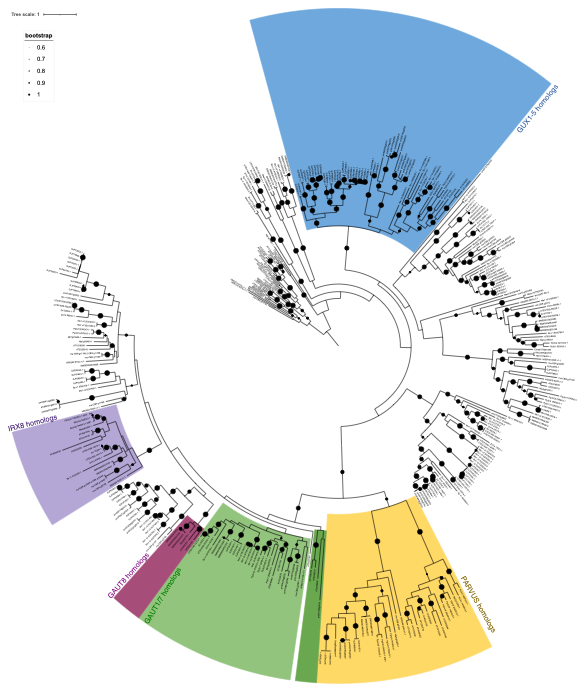

**Supplementary Figure S42: CAZyme (GT8) IRX8, GAUT8, GAUT1/7, PARVUS, GUX1-5 tree.** *Spirogyra* sequences were compared with a set of Viridiplantae species. For that, protein sequences of those families were aligned using MAFFT (in L-INS-i mode). IQTREE2 (-st AA -m TEST -bb 1000 -alrt 1000) or FASTTREE (LG+CAT, version 2.1.10) were used for inference of maximum-likelihood trees. Supplementary Data for Supplementary Figure S11-S15.

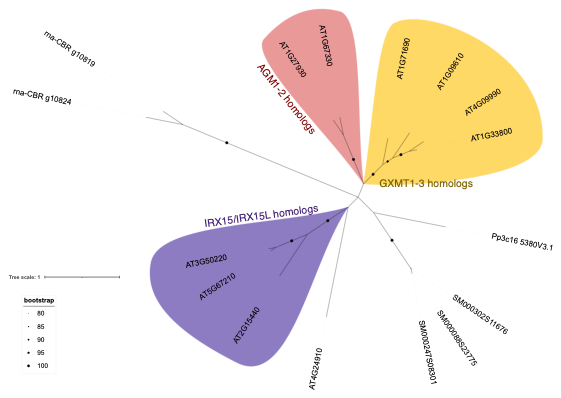

**Supplementary Figure S43: CAZyme (DUF579) IRX8, GAUT8, GAUT1/7, PARVUS, GUX1-5 tree.** *Spirogyra* sequences were compared with a set of Viridiplantae species. For that, protein sequences of those families were aligned using MAFFT (in L-INS-i mode). IQTREE2 (-st AA -m TEST -bb 1000 -alrt 1000) or FASTTREE (LG+CAT, version 2.1.10) were used for inference of maximum-likelihood trees. Supplementary Data for Supplementary Figure S11-S15.

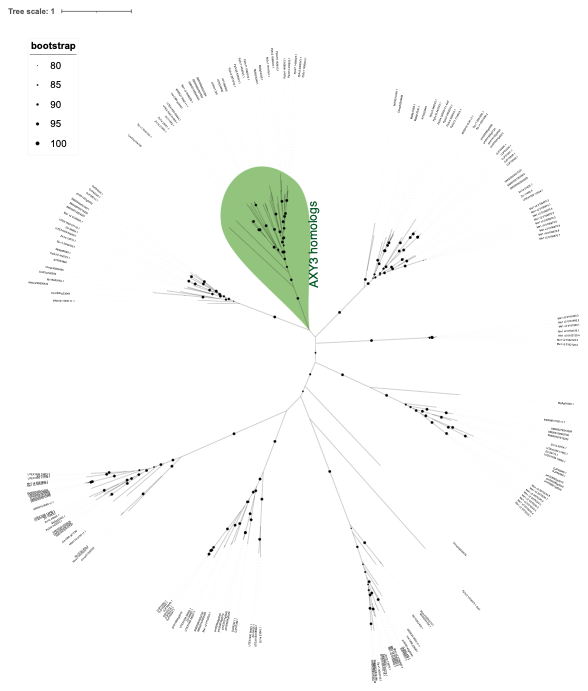

**Supplementary Figure S44: CAZyme (GH31) AXY3 tree.** *Spirogyra* sequences were compared with a set of Viridiplantae species. For that, protein sequences of those families were aligned using MAFFT (in L-INS-i mode). IQTREE2 (-st AA -m TEST -bb 1000 -alrt 1000) or FASTTREE (LG+CAT, version 2.1.10) were used for inference of maximum-likelihood trees. Supplementary Data for Supplementary Figure S11-S15.

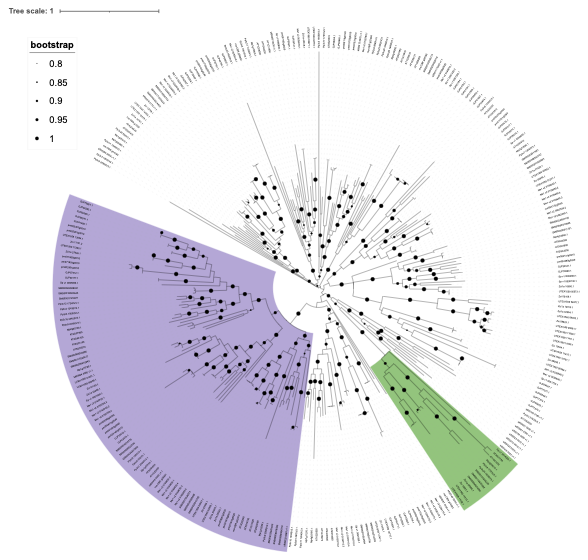

**Supplementary Figure S45: CAZyme (GT106) tree.** *Spirogyra* sequences were compared with a set of Viridiplantae species. For that, protein sequences of those families were aligned using MAFFT (in L-INS-i mode). IQTREE2 (-st AA -m TEST -bb 1000 -alrt 1000) or FASTTREE (LG+CAT, version 2.1.10) were used for inference of maximum-likelihood trees. Supplementary Data for Supplementary Figure S11-S15.

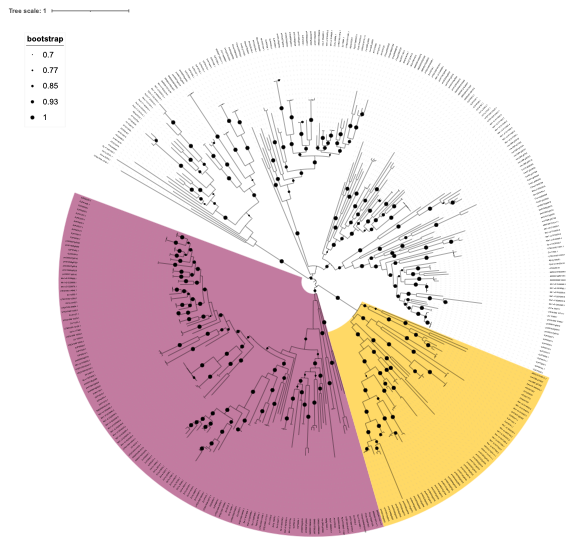

**Supplementary Figure S46: CAZyme (GT77) tree.** *Spirogyra* sequences were compared with a set of Viridiplantae species. For that, protein sequences of those families were aligned using MAFFT (in L-INS-i mode). IQTREE2 (-st AA -m TEST -bb 1000 -alrt 1000) or FASTTREE (LG+CAT, version 2.1.10) were used for inference of maximum-likelihood trees. Supplementary Data for Supplementary Figure S11-S15.

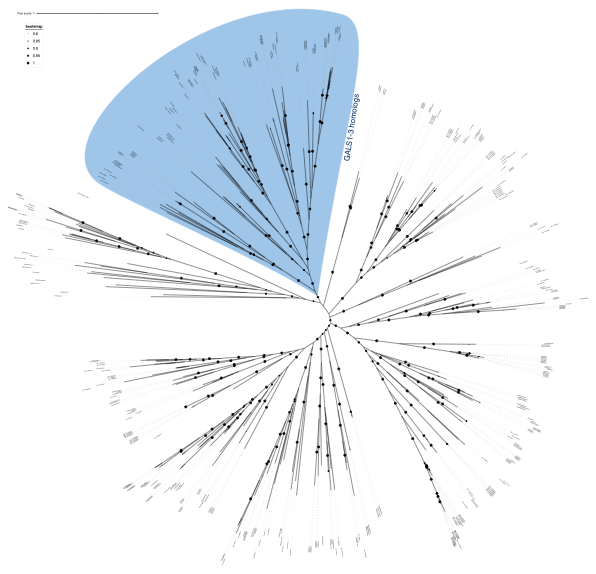

**Supplementary Figure S47: CAZyme (GT92) GALS1-3 tree.** *Spirogyra* sequences were compared with a set of Viridiplantae species. For that, protein sequences of those families were aligned using MAFFT (in L-INS-i mode). IQTREE2 (-st AA -m TEST -bb 1000 -alrt 1000) or FASTTREE (LG+CAT, version 2.1.10) were used for inference of maximum-likelihood trees. Supplementary Data for Supplementary Figure S11-S15.

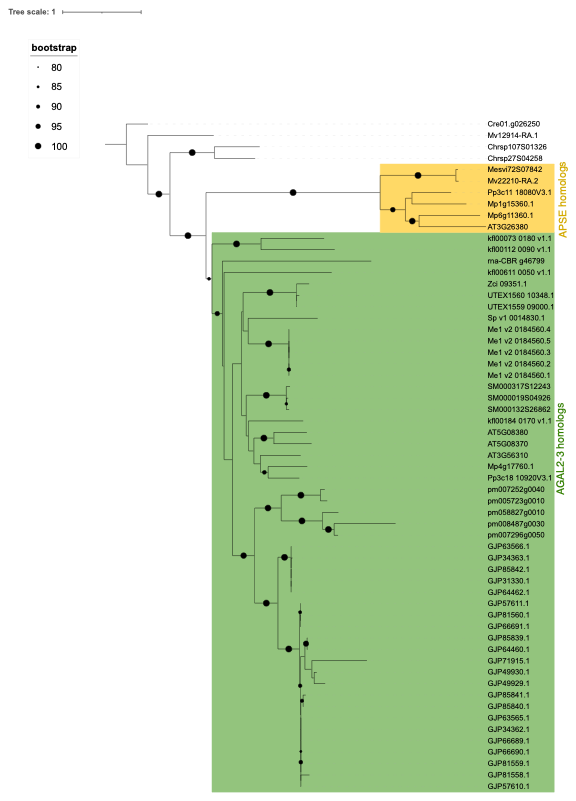

**Supplementary Figure S48: CAZyme (GH27) AGAL2-3, APSE tree.** *Spirogyra* sequences were compared with a set of Viridiplantae species. For that, protein sequences of those families were aligned using MAFFT (in L-INS-i mode). IQTREE2 (-st AA -m TEST -bb 1000 -alrt 1000) or FASTTREE (LG+CAT, version 2.1.10) were used for inference of maximum-likelihood trees. Supplementary Data for Supplementary Figure S11-S15.

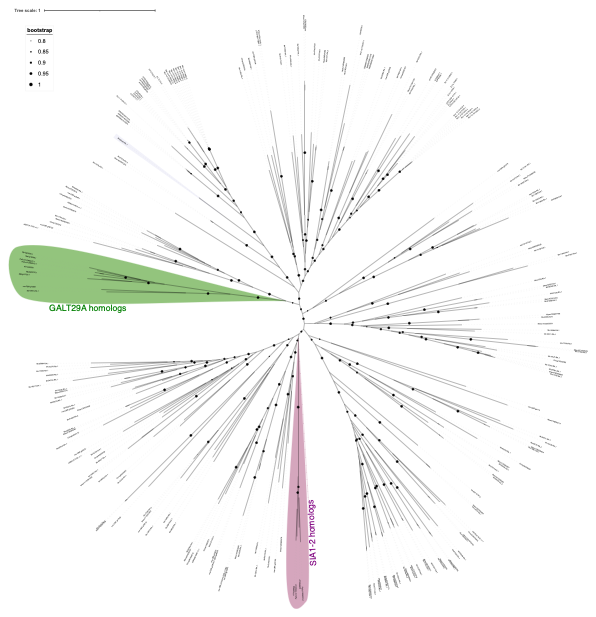

**Supplementary Figure S49: CAZyme (GT29) SIA1-2, GALT29A tree.** *Spirogyra* sequences were compared with a set of Viridiplantae species. For that, protein sequences of those families were aligned using MAFFT (in L-INS-i mode). IQTREE2 (-st AA -m TEST -bb 1000 -alrt 1000) or FASTTREE (LG+CAT, version 2.1.10) were used for inference of maximum-likelihood trees. Supplementary Data for Supplementary Figure S11-S15.

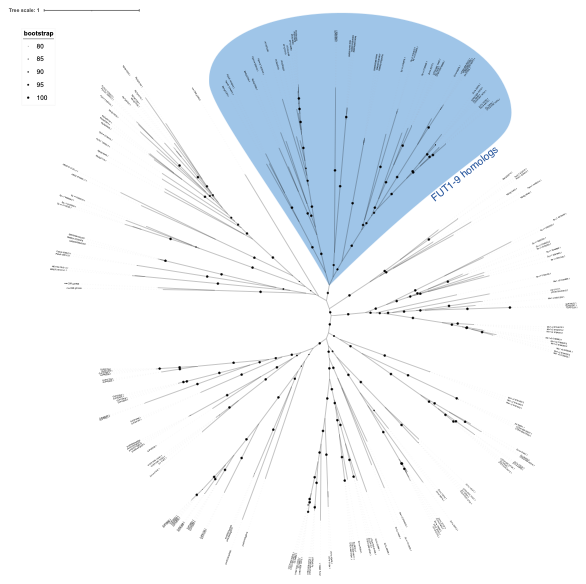

**Supplementary Figure S50: CAZyme (GT37) FUT1-9 tree.** *Spirogyra* sequences were compared with a set of Viridiplantae species. For that, protein sequences of those families were aligned using MAFFT (in L-INS-i mode). IQTREE2 (-st AA -m TEST -bb 1000 -alrt 1000) or FASTTREE (LG+CAT, version 2.1.10) were used for inference of maximum-likelihood trees. Supplementary Data for Supplementary Figure S11-S15.

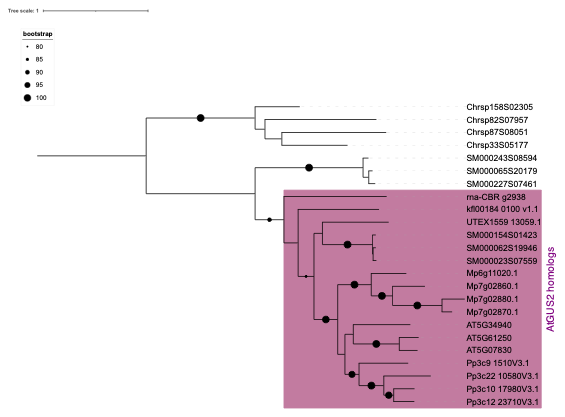

**Supplementary Figure S51: CAZyme (GH79) GUS2 tree.** *Spirogyra* sequences were compared with a set of Viridiplantae species. For that, protein sequences of those families were aligned using MAFFT (in L-INS-i mode). IQTREE2 (-st AA -m TEST -bb 1000 -alrt 1000) or FASTTREE (LG+CAT, version 2.1.10) were used for inference of maximum-likelihood trees. Supplementary Data for Supplementary Figure S11-S15.

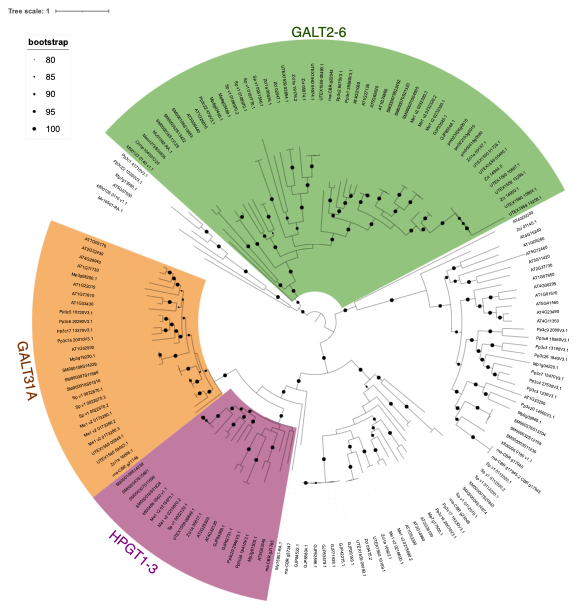

**Supplementary Figure S52: CAZyme (GT31) HPGT1-3, GALT31A and GALT2-6 tree.** *Spirogyra* sequences were compared with a set of Viridiplantae species. For that, protein sequences of those families were aligned using MAFFT (in L-INS-i mode). IQTREE2 (-st AA -m TEST -bb 1000 -alrt 1000) or FASTTREE (LG+CAT, version 2.1.10) were used for inference of maximum-likelihood trees. Supplementary Data for Supplementary Figure S11-S15.

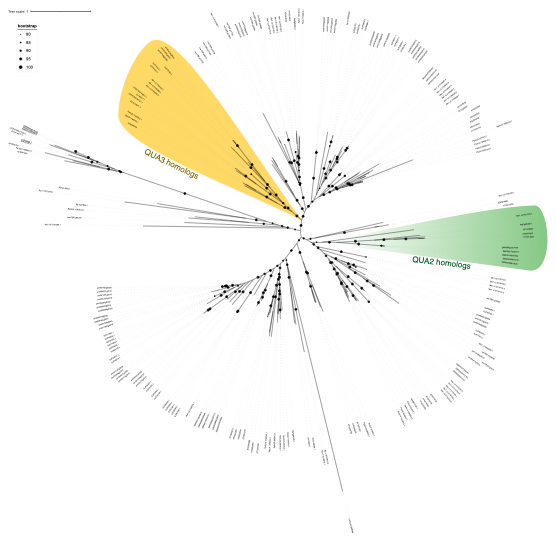

**Supplementary Figure S53: CAZyme QUA2 and QUA3 tree.** *Spirogyra* sequences were compared with a set of Viridiplantae species. For that, protein sequences of those families were aligned using MAFFT (in L-INS-i mode). IQTREE2 (-st AA -m TEST -bb 1000 -alrt 1000) or FASTTREE (LG+CAT, version 2.1.10) were used for inference of maximum-likelihood trees. Supplementary Data for Supplementary Figure S11-S15.

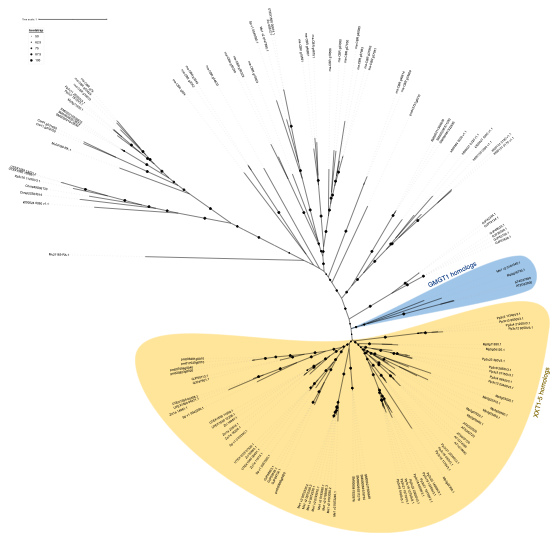

**Supplementary Figure S54: CAZyme (GT34) XXT1-5 and GMGT1 tree.** *Spirogyra* sequences were compared with a set of Viridiplantae species. For that, protein sequences of those families were aligned using MAFFT (in L-INS-i mode). IQTREE2 (-st AA -m TEST -bb 1000 -alrt 1000) or FASTTREE (LG+CAT, version 2.1.10) were used for inference of maximum-likelihood trees. Supplementary Data for Supplementary Figure S11-S15.

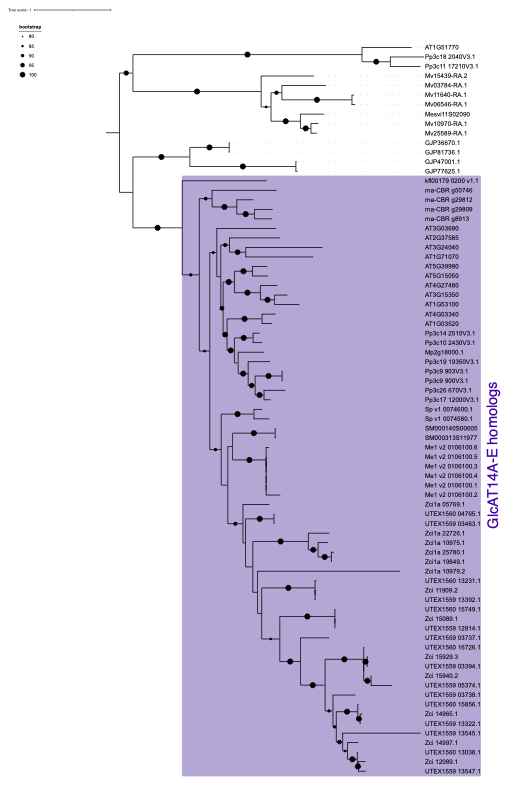

**Supplementary Figure S55: CAZyme (GT14) GlcAT14A-E tree.** *Spirogyra* sequences were compared with a set of Viridiplantae species. For that, protein sequences of those families were aligned using MAFFT (in L-INS-i mode). IQTREE2 (-st AA -m TEST -bb 1000 -alrt 1000) or FASTTREE (LG+CAT, version 2.1.10) were used for inference of maximum-likelihood trees. Supplementary Data for Supplementary Figure S11-S15.

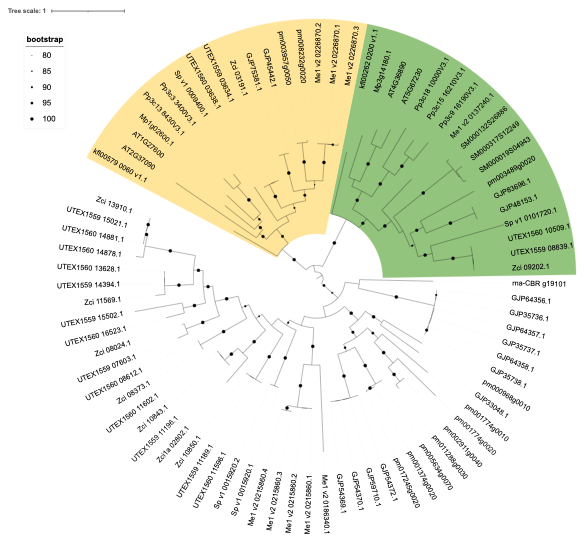

**Supplementary Figure S56: CAZyme (GT43) tree.** *Spirogyra* sequences were compared with a set of Viridiplantae species. For that, protein sequences of those families were aligned using MAFFT (in L-INS-i mode). IQTREE2 (-st AA -m TEST -bb 1000 -alrt 1000) or FASTTREE (LG+CAT, version 2.1.10) were used for inference of maximum-likelihood trees. Supplementary Data for Supplementary Figure S11-S15.

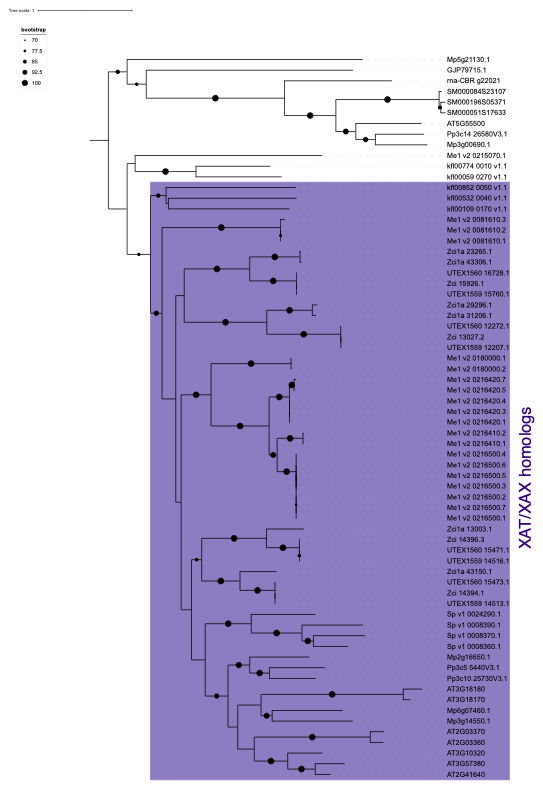

**Supplementary Figure S57: CAZyme (GT61) XAT/XAX tree.** *Spirogyra* sequences were compared with a set of Viridiplantae species. For that, protein sequences of those families were aligned using MAFFT (in L-INS-i mode). IQTREE2 (-st AA -m TEST -bb 1000 -alrt 1000) or FASTTREE (LG+CAT, version 2.1.10) were used for inference of maximum-likelihood trees. Supplementary Data for Supplementary Figure S11-S15.

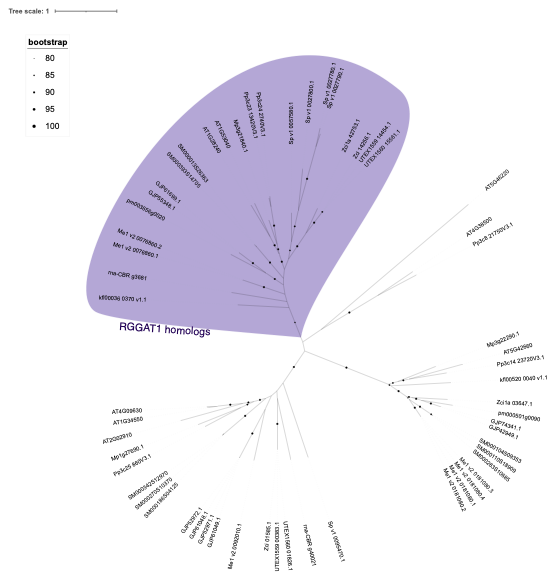

**Supplementary Figure S58: CAZyme (GT116) RGGAT1 tree.** *Spirogyra* sequences were compared with a set of Viridiplantae species. For that, protein sequences of those families were aligned using MAFFT (in L-INS-i mode). IQTREE2 (-st AA -m TEST -bb 1000 -alrt 1000) or FASTTREE (LG+CAT, version 2.1.10) were used for inference of maximum-likelihood trees. Supplementary Data for Supplementary Figure S11-S15.
